# A comprehensive engineering strategy improves potency and manufacturability of a near pan-neutralizing antibody against HIV

**DOI:** 10.1101/2024.10.14.618178

**Authors:** Mohammad M. Sajadi, Abdolrahim Abbasi, Zahra Rikhtegaran Tehrani, Christine Siska, Rutilio Clark, Woo Chi, Michael S. Seaman, Dieter Mielke, Kshitij Wagh, Qingbo Liu, Taylor Jumpa, Randal R. Ketchem, Dung N. Nguyen, Willaim D. Tolbert, Brian G. Pierce, Ben Atkinson, Derrick Deming, Megan Sprague, Andrew Asakawa, David Ferrer, Yasmin Dunn, Sarah Calvillo, Rui Yin, Johnathan D. Guest, Bette Korber, Bryan T. Mayer, Alicia H. Sato, Xin Ouyang, Scott Foulke, Parham Habibzadeh, Maryam Karimi, Arash Aslanabadi, Mahsa Hojabri, Saman Saadat, Roza Zareidoodeji, Mateusz Kędzior, Edwin Pozharski, Alonso Heredia, David Montefiori, Guido Ferrari, Marzena Pazgier, George K. Lewis, Joseph G. Jardine, Paolo Lusso, Anthony DeVico

## Abstract

Anti-HIV envelope broadly neutralizing antibodies (bnAbs) are alternatives to conventional antiretrovirals with the potential to prevent and treat infection, reduce latent reservoirs, and/or mediate a functional cure. Clinical trials with “first generation” bnAbs used alone or in combination show promising antiviral effects but also highlight that additional engineering of “enhanced” antibodies will be required for optimal clinical utility, while preserving or enhancing cGMP manufacturing capability. Here we report the engineering of an anti-CD4 binding-site (CD4bs) bnAb, N49P9.3, purified from the plasma of an HIV elite-neutralizer. Through a series of rational modifications we produced a variant that demonstrates: enhanced potency; superior antiviral activity in combination with other bnAbs; low polyreactivity; and longer circulating half-life. Additional engineering for manufacturing produced a final variant, eN49P9, with properties conducive to cGMP production. Overall, these efforts demonstrate the feasibility of developing enhanced anti-CD4bs bnAbs with greatly improved antiviral properties as well as potential translational value.

## Introduction

Currently, antiretroviral agents (ARVs), most of which target HIV replication enzymes, are the only available resources for the prevention and treatment of AIDS. Although highly effective, these drugs present noteworthy limitations including differential penetration into tissue sanctuaries such as CNS, numerous side effects relevant to lifelong administration, and the inability to provide long-term, drug-free remission or functional cure. Advances in newer integrase strand transfer inhibitor (ISTI) ARVs provide superior efficacy and less frequent delivery (e.g. bi-or tri-monthly) versus reverse transcriptase inhibitors, yet also present potential long-term safety concerns. For example, numerous reports link ISTI with diabetes and metabolic syndrome [1–6]. The capsid inhibitor lenacapavir has the capability of Q6 month injection, and to date, its only drawbacks are the lag time prior to full efficacy (about a month), as well as injection site reactions [7].

BnAbs against the HIV envelope (Env) trimer are promising alternatives or ARV-complementing agents for HIV prevention and treatment; reduction of latent virus reservoirs; and/or cure strategies. Such antibodies arise for as yet unknown reasons only in a small percentage of individuals who are chronically infected with HIV [8] [9], often after a years-long period of accumulating substantial somatic hypermutation [10]. BnAbs target an array of conserved sites (reviewed in [11]) within the Env trimer: the apex comprised of gp120 V1/V2 amino acids and glycans; the gp120 V3 loop domains comprising the N332-glycan; high-mannose regions in the gp120 outer domain [12]; the CD4-binding site (CD4bs) of gp120 [13]; the membrane proximal region (MPER) of gp41 (reviewed in [14]); and a combination of gp120-gp41 interface sequences [15]. Although each class has specific strengths and limitations, a general advantage of bnAbs is that they lack toxicity and may act against HIV through multiple effector mechanisms including direct virus inactivation as well as Fc-mediated phagocytosis and killing of infected cells [16, 17].

Clinical testing of “first generation” human bnAbs has yielded promising results regarding safety and efficacy. The AMP trial of the anti-CD4bs antibody, VRC01, found that participants that received infusions of recombinantly produced VRC01 IgG were protected against viruses that were potently neutralized by VRC01 [18]. However, overall efficacy in the trial was negligible as VRC01 lacked sufficient neutralization potency to protect against the majority of circulating viral strains [18]. The recently completed TITAN trial of 3BNC117 (anti-CD4bs) and 10-1074 (anti-V3) highlighted the promise of dual bnAb treatment in eliciting long-term viral suppression [19]. The TATELO trial of VRC01 and 10-1074 in a pediatric cohort demonstrated that dual bnAb treatment safely maintained HIV suppression for up to 24 weeks in 44% of children after cessation of conventional ARVs [20]. Such findings suggest that bnAbs could offer a feasible strategy of ART-sparing in children to avoid long-term toxicities and adherence issues. Furthermore, through a process still under investigation, bnAb treatment appears to mediate a so-called “vaccinal effect” that promotes more efficient host immune control of infection in addition to mediating neutralization [21]. Thus, bnAb treatment could enable functional cure strategies [22]. Notably, it is becoming increasingly clear that at-risk subjects and people living with HIV (PLWH) are most receptive to a range of treatment options, including bnAbs, that suit individual needs [23].

While these studies support a rationale for bnAb-based strategies to prevent or control HIV/AIDS, they also highlight why full clinical value will require greater breadth and potency of action. In trials where bnAb Fab domains resembling their natural counterparts are used to screen resistant viruses, up to 46% of subjects may be excluded based on prescreening that identified the presence of resistant virus [24]. Fortunately, the amount of information now available supports rational engineering of bnAbs to achieve greater potency, breadth, and pharmacokinetic (PK) characteristics, while allowing feasible cGMP production at clinical scale. Here we report such engineering of a near pan-neutralizing anti-CD4bs bnAb, N49P9.3, isolated from the plasma of donor N49, an HIV elite neutralizer [25]. We reasoned that the natural characteristics of a circulating plasma antibody would promote translational development. We describe modifications that enhance the breath and potency of Fab and Fc-mediated antiviral effects, improve PK in animal models and facilitate cGMP manufacture. Such constructs should increase the prospects for successful bnAb-based treatment and prevention options in HIV/AIDS.

## Material and Methods

### Antibody isolation and production

The B-cell database of volunteer N49, previously assembled [25], was searched for all clones with characteristic CD4bs deletion in CDRL3, and found sequences aligned to identify those in the N49P9 family. The identified VH or VL region clones were cloned into an expression vector upstream to a human IgG1*03 backbone for heavy chain and either a k or λ light chain expression vector. The paired plasmids were used to transiently transfect 293 Freestyle cells. Transfectant supernatants were used for purification of the mAbs using protein A affinity chromatography.

### ELISA

The HIV binding ELISA was performed with monomeric BAL-gp120 antigen. Bal-gp120 was directly coated (4 µg /ml) on the plates, but otherwise ELISA was the same as previously described [25]. Each assay performed in duplicate at multiple dilutions and EC50 calculated using GraphPad Prism 10 software (Boston, MA).

### Neutralization assay

HIV-1 neutralization testing was performed using a luciferase-based assay in TZM.bl cells, as previously described [26]. Briefly, three-or five-fold serial dilution of plasma, supernatant, or monoclonal antibodies were tested against a panel of Env-pseudotyped viruses in duplicate. Following a 48 hour incubation, the reduction in luciferase expression following a single round of virus infection was measured using Bright-Glo luciferase reagent and GloMax luminometer (Promega, Madison, WI). The concentrations of antibody that inhibited 50% or 80% of viral infection (IC50 and IC80 titers, respectively) were calculated using 5-parameter curve fitting (LabKey NAb Tool) [27].

### Clinical polyreactivity testing

Antibodies were tested in a CLIA-certified clinical lab (University of Maryland Medical Center) for rheumatoid factor, anti-centromere antibody, anti-Jo1 antibody, anti-Sjögren’s-syndrome-related antigen A (SSA), anti-Sjögren’s-syndrome-related antigen B (SSB), anti-topoisomerase I (SCL-70), anti-ribonucleoprotein (RNP), and anti-smooth muscle (SM) antibodies. Anti-nuclear antibody (ANA) testing in Hep 2 cells was carried out with Bio-Rad ANA Screening Test (Hercules, CA). For all assays, mAbs were tested at 25 µg /ml, with the positive and negative controls that were assay-specific.

### ADCC assay

#### Env-IMC construction and virus production

To construct Env-IMCs which contain the ectodomain of the Envelope of interest (HxB2 positions 41-687) and an isogenic NL4.3 backbone as previously described [28], partial *envelope* genes were amplified by PCR to produce amplicons which contained a 20 base pair overlap with the pNL4.3 backbone on both the 5’ (HxB2 positions 6353-6335) and 3’ (HxB2 positions 8283-8299) ends. The pNL4.3 plasmid (BEI Resources, Cat. no. ARP-114) was then PCR amplified to generate a product which excluded the Env ectodomain nucleotides but included the signal peptide and endodomain regions of NL4.3 *envelope*. The pNL4.3 and partial *envelope* PCR products were then fused using NEBuilder HiFi DNA Assembly (NEB, Ipswich MA) per the manufacturer’s recommendations to produce an NL4.3-*env.*ecto chimeric HIV infectious molecular clone (Env-IMC) which contained a complete, in-frame *envelope*. Viruses were produced by transfecting the Env-IMCs into HEK293T cells using Dreamfect Gold transfection reagent (OZ Biosciences, San Diego CA). Virus-containing supernatants were harvested 48 h following transfection, clarified by 0.45 µm filtration and adjusted to 10% FCS (Nucleus Biologics).

#### Target cell infection

1×10^6^ D660 cells (CEM.NKR_CCR5_ cells which contain GFP and Renilla *luciferase* genes under the control of the HIV-1 LTR; kindly provided by Dr. John Kappes, University of Alabama at Birmingham) were infected with 1mL of virus. Briefly, 2µL of Lentiblast (OZ Biosciences) was added to 1mL of virus and incubated for 10 minutes.

D660 cells were then pelleted by centrifugation at 485xg for 5 minutes and the virus/Lentiblast mixture was added to the cells. Cells and virus were centrifuged at 1200xg for 2 hours at 30°C. After centrifugation, cells were resuspended in R10 to a final concentration of 5×10^5^ cells/mL and incubated for 72 hours.

#### Renilla Luciferase-based ADCC assay

A modified version of the LucR-based ADCC assay previously described [29] was conducted: the day prior to the ADCC assay, cryopreserved PBMCs to be used as effectors in the assay were thawed in R10, counted and assessed for viability and resuspended in R10 overnight. On the day of the assay, infected D660 cells were counted, assessed for viability (viability was ≥ 80% to be used in the assay) and the concentration was adjusted to 2 x 10^5^ viable cells/mL (5 x 10^3^ cells/well). PBMCs were then counted, assessed for viability, pelleted and resuspended in the infected D660 cells at a concentration of 6 x 10^6^ PBMCs/mL (1.5 x 10^5^ PBMCs/well; effector: target cell ratio of 30:1). Heat-inactivated autologous plasma was serially diluted. The effector/ target cell mix and antibody dilutions were plated in opaque 96-well half-area plates, centrifuged at 300 x g for 1 minute after 30 minutes incubation at room temperature, and then incubated for 5.5 hours at 37°C, 5.5% CO_2_ to allow ADCC-mediated cell lysis to proceed. After 5.5 hours, ViviRen substrate (Promega) was diluted 1:500 in R10 and added 1:1 to the assay wells. The substrate generates luminescence only in live, infected cells; not in dead or lysed cells. The final readout was the luminescence intensity generated by the presence of residual intact target cells that have not been lysed by the effector population in the presence of ADCC-mediating antibodies. The percentage of specific killing was calculated using the formula

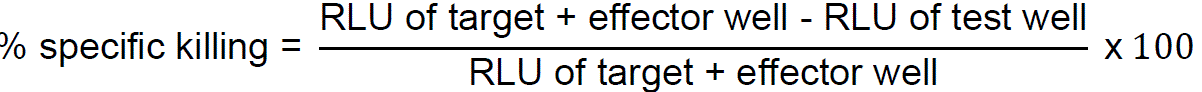

In the analysis, the RLU of the target + effector wells represent lysis by NK cells in the absence of any source of antibody. The specific killing activities in the ADCC assay were summarized for each mAb and virus by computing the area under the dilution curve (AUC) as the mean of the specific killing activities across the dilution series. Specific killing activities less than 10% were assigned a value of 0 prior to AUC calculation.

### Antibody combination analysis

#### BnAb and bnAb combination modeling

To compare individual bnAbs, we predicted IC99 and IIP for each bnAb versus each virus using the mathematical modeling previously outlined [30]. Briefly, neutralization curve slopes were calculated using IC50 and IC80 titers and these with IC50 titers characterize the Hill curve that determines the full neutralization curve, which then allows calculation of IC99 (concentration at which 99% of the viral sample is neutralized) and IIP (Log10 fold reduction in a single round of replication in the presence of bnAb). For bnAb combinations the Bliss-Hill model [30] was used that can predict the full neutralization for the bnAb combination using individual bnAb IC50 and IC80 titers. This mathematical description of the full curve can be solved to obtain IC50, IC80 and IC99 titers and IIP for the combination. Titers for bnAb combinations are calculated assuming each bnAb in the combination is present at the same concentration, and they are reported as the sum of concentrations of each bnAb (e.g. if a 2-bnAb combination has an IC50 = 1 µg/ml, then each bnAb is present at 0.5 µg/ml; similarly if a 3-bnAb combination has IC50=1 µg/ml, then each bnAb is present at 0.33 µg/ml). For ranking, 6 summary metrics were used for individual bnAbs: 1) IC80 geometric mean, 2) IC80 < 1 µg/ml breadth, 3) IC99 geometric mean, 4) IC99 < 10 µg/ml breadth, 5) median IIP, 6) IIP>5Log10 breadth. For bnAb combinations, additionally IC80 breadth with 2 or more bnAbs active was also considered. Each individual bnAb or bnAb combination was compared to that of other individual bnAbs or combinations with the same number of bnAbs. For each metric, rank was calculated as 1 + the number of bnAb/combinations strictly better (not better than or equal to) than the bnAb or combination in question.

#### Antibody engineering for potency

In order to enhance binding affinity and avidity, we first undertook a strategy to engrafting the heavy chain framework 3 loop (“-FR”) from VRC03 to enable quaternary interaction with a neighboring gp120 protomer, as previously reported for other antibodies [31, 32]. Thus, N49P9.6 was engineered into N49P9.6-FR, each Fab arm with an unconventional bnAb architecture that presents two paratopes reactive with two neighboring protomers on the Env trimer, mimicking the quaternary binding mode of CD4 [33]. To optimize the N49P9.6-FR antibody, we utilized multiple rational and structure-based design approaches to improve its biophysical properties and antigen targeting. Based in part on previous successful HIV antibody design efforts [34, 35], we assessed germline revertant mutations and consensus residue mutations from VRC01-class antibodies using the Therapeutic Antibody Profiler (TAP) tool [36], which assesses antibody developability using structure-based metrics, as well as Rosetta [37], which was used to calculate antibody stability or binding energy changes (ΔΔGs) for individual substitutions based on modeling. Favorable substitutions based on one or both of those methods were prioritized for experimental testing. We additionally tested a substitution to remove a possible N-glycosylation site in the heavy chain V domain, given that N-glycan removal was previously shown to be effective for antibodies [38], as well as a CDRL2 loop residue swap from the related VRC07 antibody [34]. Finally, two substitutions were selected based on computational scanning mutagenesis of the heavy chain framework-3 loop in Rosetta and favorable predicted Env targeting and/or thermostability.

#### Antibody engineering for developability

Sequence stability violations were determined for N49P9.6-FR using a Just proprietary Residual Artificial Neural network for the Design of Antibodies (RANDŸ). VH and VL were designed based on all possible amino acid combinations at the identified sites. These VH and VL designs were DNA synthesized at Twist Bioscience (San Francisco, CA) as library pools for the VH and VL regions, separately. The V-regions were cloned into a phagemid vector and M13 Fab display libraries were generated as described [39].

The gp160 target (CH505TFchim.6R.SOSIP.664v4.1) was randomly biotinylated on primary amines using the EZ-link NHS-Biotin kit per manufacturer’s recommendations (ThermoFisher Scientific, Waltham, MA). Multiple rounds of Fab phage panning were performed using a MagBead Kingfisher platform. The first panning round used 100nM biotinylated gp160 in PBS to allow Fab-phage binding. Four, 10min PBS washes were performed followed by phage elution at pH 2.5 via 0.2M glycine. Subsequent rounds had multiple arms where temperature challenge was conducted in one arm whereas GnHCl challenges were conducted in the other.

Temperatures ranged from 65°C to 75°C and GnHCl concentrations ranged from 0.25 to 1.0 M to increase selection of more stable Fabs. Each library challenge was for 10min. Additionally, the gp160 target concentration was decreased from 20 nM in the second round to 0.8nM in the last round. Sanger sequencing and polyclonal ELISA using the same gp160 as used in the panning was performed after rounds 1-3 to confirm Fab enrichment. 48-96 Fabs from all arms were converted to IgG1 in a batch, two step, high-throughput cloning process. These sequenced plasmids were transiently transfected in duplicate into HEK293 cells for small-scale expression in 96well blocks as per manufacturer’s recommendations (ThermoFisher Scientific, Waltham MA). After 4 days of expression, the supernatants were harvested and antibody secretory titers measured via BLI. These crude supernatants were then assessed for neutralization activity in up to 10 pseudovirus panel (first tested against 5 pseudoviruses, and if no loss of neutralization, tested against additional 5 pseudoviruses). Loss of neutralization defined as > 5 fold loss of IC50 or IC80 against any single pseudovirus. The top candidates as determined by preservation of neutralization activity were scaled-up in the 293 transient expression system as described but rather in 24well blocks or 125mL flasks to provide enough material for further downstream analytics. The supernatants were harvested after a 4-day expression and the titers measured via BLI. The secreted antibodies were Protein A purified. Samples were buffer exchanged against 10 diavolumes of 20mM sodium phosphate, 150mM sodium chloride, pH 7.1 (PBS) using a centrifugal filter with a 30 kDa molecular weight cut off (Amicon). After buffer exchange, samples were normalized to 1 mg/ml using a Lunatic protein concentration plate format instrument (Unchained Labs, Pleasanton, CA). The antibodies were then advanced for biophysical characterization and further neutralization activity assessment.

#### Generation of stable pool for final candidate variants

Horizon GS KO host cells were cultured in CD OptiCHO growth medium containing 4 mM glutamine. One day prior to transfection, host cells were passaged at 1E6 cells/mL to maintain the cells in exponential growth phase. HT method transfection was performed to create stable pool replicates, each receiving a total of 10ug of N49P9.6-FR expression vector and transposase mRNA using ECM-830 BTX electroporator. Immediately after transfection, cells were transferred to a 24 DWP containing 2 mL of warm glutamine supplemented CD OptiCHO for a 3-day recovery. Selection of transfectants began with culturing the cells in glutamine-free CD OptiCHO medium. Stable pools reached 98% viability by selection day 11 and were subsequently subjected to a 10-day fed batch production for early material generation, evaluation of productivity and protein quality prior to single-cell cloning.

#### Differential Scanning Fluorimetry (DSF)

Thermal transition temperature(s) and weighted shoulder scores were determined by DSF according to the method previously described [40]. A single biological sample was divided, and the assay ran twice per molecule type. Additional information was also obtained from a parameter termed the weighted shoulder score (WSS) which accounts for multiple pieces of information from the unfolding curve [41].

#### Low pH stability

Stability during a low pH hold was determined according to the method previously described [40]. The increase in high molecular weight of the low pH exposed sample as compared with the control sample was reported.

Any increase above 10% high molecular weight after low pH is considered undesirable and will present additional manufacturing challenges.

#### Chemical unfolding

The chemical unfolding assay was completed as previously described [42], with some modifications. After a 3-day incubation in 32 guanidine hydrochloride (GND) concentrations, the samples were measured on a Fluorescence Innovations SUPR-UV plate reader (excitation: 275 nm, emission: 300-440 nm). The measured fluorescence intensity at 362 nm was corrected for scattering and stray light, the unfolding curve was generated by graphing each corrected intensity against the GND concentration and the inflection point was reported. Inflection point range between 1.5 to 2.5 molar GND has been observed internally, with higher inflection points being reflective of higher conformational stability.

#### Relative solubility

Solubility was assessed according to the method previously described [43]. Analysis was done in PBS buffer (20mM sodium phosphate and 150mM sodium chloride pH 7.1) and a final PEG 10,000 concentration ranging from 7.2% to 9.6%. Percent recovery relative to a 0% PEG control was determined and average recovery across the PEG concentration range was reported.

#### Self-Interaction Nanoparticle Spectroscopy (SINS)

SINS measurements were performed according to the method previously described [44]. Briefly, gold nanoparticles (Ted Pella, Redding, CA) were conjugated overnight with an 80:20 ratio of anti-human and anti-goat antibodies (Jackson Immuno Research, West Grove, PA). Unreacted sites were blocked using an aqueous 0.01% (w/v) polysorbate 80 solution. Conjugated gold nanoparticles were then concentrated by centrifugation and removal of 95% of the supernatant. Analysis was carried out in PBS (20mM phosphate, 150mM NaCl, pH 7.1) at a protein concentration of 0.05 mg/ml reacted with 5 µl of concentrated conjugated gold nanoparticles. After a 2-hour incubation, absorbance spectrum from 400-600 nm was collected using a Spectrostar Nano plate reader at 2nm steps. The wavelength maximum of the spectrum peak is reported. Three assay replicates were made from each of the pooled expressed antibodies.

#### Standup Monolayer Absorption Chromatography (SMAC)

SMAC measurements were performed according to the method previously described [45]. Retention times were determined using a Dionex UPLC equipped with a Zenix column (Sepax Technologies) and a running buffer comprised of 150mM sodium phosphate pH 7.0.

#### Polyreactivity

Polyreactivity testing was performed according to the method previously described [46]. An ELISA assay was used to test against 4 different antigens: KLH, Insulin, dsDNA and Cardiolipin (CL). Samples are diluted to 1 µg/mL and a secondary anti-human antibody is used to detect the amount of protein that has bound to the different antigens. After substrate addition, absorbance is measured at 405 nm and subtracted against a PBS blank sample. An absorbance range between 0 to 3 has been observed.

#### Viscosity

Samples were buffer exchanged into 10mM acetate, 9% sucrose, pH 5.2 and prepared at various concentrations from 85 mg/mL to 175mg/mL as measured in triplicate by Solo VPE (Repligen, Waltham Massachusetts). All viscosity measurements were performed at 20°C on an MCR-92 cone and plate rheometer (Anton Paar, Graz Austria) equipped with a CP20-0.5 cone. A shear rate sweep from 50 to 2000 1/s was performed using a sample volume of 25uL per measurement. Viscosity results for sheer rates between 500 and 1000 1/s were averaged and reported per protein concentration.

### X-ray Crystallography

#### Protein expression and purification

BG505 SOSIP.664 and Clade A/E gp120_93TH057_core_e_ were expressed in HEK293 GnTI^-^ cells. Cells were first seeded at 1 x 10^6^ cells/ml (viability >90%) into an expression flask and then transfected with plasmids encoding for the BG505 SOSIP.664 or Clade A/E gp120_93TH057_core_e_ envelope trimer and furin at a molar ratio of 4:1 (SOSIP:furin), as previously described [47, 48]. Transfected cells were incubated for seven days at 37° C, in a humidified atmosphere of 8% CO_2_ on a shaker rotating at 125 rpm. Cell supernatants were then collected, centrifuged and filtered through a 0.22 µm PES membrane to remove cell debris. BG505 SOSIP.664 trimer was purified by N49P9.1 affinity chromatography, i.e. N49P9.1 IgG covalently bound to protein A. BG505 SOSIP.664 was eluted from the column with 3 M magnesium chloride. Eluted protein was immediately diluted in DPBS and then the buffer was exchanged to DPBS using Amicon microcon centrifugal concentrators (Millipore Sigma). Clade A/E gp120_93TH057_core_e_ was purified by 17b affinity chromatography, i.e. 17b IgG covalently linked to protein A. Eluted protein was immediately mixed with 0.1 M Tris-HCl pH 8.5 to raise the pH and the buffer was then exchanged to DPBS using Amicon microcon centrifugal concentrators. Clade A/E gp120_93TH057_core_e_ and BG505 SOSIP.664 was further purified by size exclusion chromatography (SEC) using a Superdex 200 Increase 10/300 GL column (Cytiva) equilibrated with Dulbecco’s phosphate buffered saline (DPBS) buffer. Fabs were produced by papain digestion of IgGs. Briefly IgGs were incubated at 37° C with immobilized papain (Thermo Fisher) for 3-4 hours. Fabs were then separated from Fcs and undigested IgGs by passage over a HiTrap protein A column (Cytiva). Fabs were further purified by gel filtration chromatography on a Superdex 200 16/60 column equilibrated in DPBS. Purified Fabs were then concentrated for use in complex formation.

#### X-ray complex formation and purification

Clade A/E gp120_93TH057_core_e_ that had been produced in GnTI^-^ cells had its glycans removed with EndoH_f_ (New England Biolabs) before its use in complex formation. Briefly, ten units of EndoH_f_ were added per microgram of protein. The protein was incubated at 37° C overnight and the EndoH_f_ then removed by passage over an amylose column. Complexes were made by mixture of deglycosylated gp120_93TH057_core_e_ with an excess of Fab at a ratio of 1.1:1 (Fab:gp120). Complexes were purified by gel filtration chromatography on a Superdex 200 16/60 column equilibrated with 10 mM Tris-HCl pH 7.2 and 100 mM ammonium acetate. Purified complexes were concentrated to approximately 5-10 mg/ml for use in crystallization trials.

#### Crystallization

Initial crystallization trials were done with commercially available sparse matrix crystallization screens from Molecular Dimensions (Proplex Eco and MacroSol Eco) using the hanging-drop, vapor diffusion method. The screens were incubated at 21° C and monitored periodically for the production of protein crystals. Conditions that produced crystals were then reproduced and optimized. N49P9.1 Fab-gp120_93TH057_core_e_ complex crystals grew from 12% PEG 6000 and 0.1 M Bis-Tris pH 6.5. N49P9.3 Fab-gp120_93TH057_core_e_ complex and N49P9.3-FR Fab-gp120_93TH057_core_e_ complex crystals grew from 10% PEG 4000, 0.1 M 4-(2-hydroxyethyl)-1-piperazine ethanesulfonic acid (HEPES) pH 7.5, and 0.1 M magnesium chloride. Prior to data collection crystals were briefly soaked in crystallization condition supplemented with 20% 2-methyl-2,4-pentanediol (MPD) as a cryoprotectant and then flash frozen in liquid nitrogen.

#### Data collection and refinement

Diffraction data of the N49P9.3 Fab-gp120_93TH057_core_e_ complex was collected at the Stanford Synchrotron Radiation Light Source (SSRL) beam line 9.2 and data for the N49P9.1 Fab-gp120_93TH057_core_e_ and N49P9.3-FR Fab-gp120_93TH057_core_e_ complexes were collected at SSRL beam line 12-2. Both beam lines were equipped with Pilatus area detectors. The N49P9.1 Fab-gp120_93TH057_core_e_ complex crystals belonged to space group I2_1_2_1_2_1_ with cell dimensions of a=104.8 Å, b=110.5 Å, and c=152.2 Å and diffracted to 3.15 Å resolution. N49P9.3 Fab-gp120_93TH057_core_e_ complex crystals belonged to space group P1 with cell dimensions of a=62.2, b=68.2, c=115.1, α=90.3°, β=102.3°, and ɣ=90.3° and diffracted to 3.4 Å resolution. N49P9.3-FR Fab-gp120_93TH057_core_e_ complex crystals belonged to space group P1 with cell dimensions of a=60.4, b=65.4, c=112.4, α=90.0°, β=104.9°, and ɣ=90.0° and diffracted to 2.15 Å resolution. The N49P9.1 complex structure was solved by molecular replacement with Phaser from the CCP4 suite based on the coordinates of gp120 from PDB ID 3TGT and the VRC01 Fab from PDB ID 4RFE for the N49P9.1 Fab. The N49P9.3 and N49P9.3-FR complex structures were solved using the refined coordinates from the N49P9.1 complex structure (PDB ID 6OZ3). Refinement was done with Refmac and/or Phenix, coupled with manual refitting and rebuilding using COOT, as described in [49]. The quality of the final refined models were monitored using the program MolProbity, as described in [49]. All illustrations were prepared with the PyMol Molecular Graphic suite (http://pymol.org) (DeLano Scientific, San Carlos, CA, USA). Data collection and refinement statistics are shown in **Table S5**. Final structures have been deposited in the Protein Data Bank (PDB) with accession codes 6OZ3, 7SX6, and 7SX7. N49 structures were compared to VRC03 (PDB ID 6CDI), VRC01-FR (PDB ID 6NNF), and N6-FR (PDB ID 6NM6).

### Cryo Electron Microscopy (CryoEM)

#### CryoEM sample preparation and data collection

Protein complex was made by incubating an excess of PGT121 and N49P9.6-FR Fabs with GnTI^-^ produced BG505 SOSIP.664 trimer overnight at room temperature at a molar ratio of 10:10:1, respectively. The complex was purified by SEC on a Superdex 200 Increase 10/300 GL column (Cytiva) in DPBS buffer. The complex peak was pooled and concentrated to 2.3 mg/ml. 5 µl of protein complex was deposited on a holey copper grid (QUANTIFOIL R1.2/1.3, 200 mesh, EMS cat# Q250-CR1.3) or a carbon film copper grid (QUANTIFOIL R 1.2/1.3, UT, 200 mesh, EMS cat# Q250-CR1.3-2NM) which had been glow-discharged for 30s at 25 mA using PELCO easiGlow (TedPella Inc.). CryoEM grids were double-blotted and vitrified in liquid ethane using Vitrobot Mark IV (Thermo Fisher) with a blot time of 1s and variable blot forces at 4°C and 100% humidity.

The cryopreserved grids were screened on a FEI Talos Artica microscope at 200 kV equipped with a FEI Falcon3EC detector using the EPU software (Thermo Fisher). The CryoEM data from high quality grids were then collected on a FEI Glacios electron microscope operating at 200 kV, equipped with a Gatan K3 direct electron detector. Micrographs were collected at a magnification of 56,000 corresponding to a pixel size of 0.889 Å with a total exposure dose of 55 or 59 e−/Å^2^.

#### CryoEM data processing, model building and refinement

Motion correction, CTF estimation, particle picking, curation, extraction, 2D classification, ab initio reconstruction, volume refinements and local resolution were performed using CryoSPARC. Particles extracted from a holey carbon grid (Q250-CR1.3 grid) dataset and a holey carbon grid with ultrathin carbon (Q250-CR1.3-2NM) dataset were combined after the particle extraction step in order to provide top and bottom views of the complex. Combined particles were subjected to 2D reference-free classification in CryoSPARC and good 2D classes were selected and subjected to multiple rounds of 2D classification.

Initial models of BG505 SOSIP from PDB ID 6CDI, PGT121 from PDB ID 5CEZ and N49P9.3-FR from PDB ID 7SX7 (this manuscript) were used as templates. Initial model-to-map fitting cross correlation was carried out in UCSF ChimeraX. Several rounds of model refinement were carried out in Phenix (real-space refinement) and Coot (manual refinement). Geometry validation and evaluation were performed by EM-Ringer and Molprobity. Resolution of the structure was further assisted by published crystallographic information positioning the PGT 121 Fab on the BG505 SOSIP.664 trimer [50]. The final model was refined to acceptable geometry with a Chimera CC score of 0.71. The statistics of data collection, reconstruction and refinement are described in **Table S6** and the structure has been deposited in the PDB with accession code 7UOJ.

### In vivo testing

#### PK and half-life testing

PK testing was carried out in FcRn−/− hFcRn transgenic mice (Jackson Laboratory). These mice, which harbor a knockout allele of the FcRn α-chain (Fcgrt^tm1Dcr^) and express the human FcRn α-chain, have been extensively studied and used for measuring IgG half-life of human mAbs and Fc engineered mutants. Mice (9 weeks old, equal male and female) were injected IP with 10mg/kg of antibody, and 7 serial blood draws at 0 hours, 8 hours, and days 1, 3, 7, 10,14, 21, and 28 days and IgG Elisa specific for Human IgG measured. Non-compartmental analysis (NCA) was used to calculate the mean of the individual-level terminal half-life estimates for each animal and mAb. Concentrations below the lower limit of the assay were excluded from the analysis. The NCA approach estimates the slope of the terminal elimination phase (λz), where λz was calculated via a linear regression between Y=log(concentrations) and X=time after injection. The time interval was fixed to use all available time points starting with day 3 post-injection.

#### Generation of humanized mice

For generation of humanized CD34 mice, newborn mice from strains NSG-SGM3 or NSG-IL15 were exposed to total body radiation (100 cGy) per mouse in a RS-2000 x-ray radiator, followed by intrahepatic injection of 10^5^ CD34+ cells under anesthesia (ice for ∼ 3 minutes until gross movements cease). Human CD34+ cells, isolated from cord blood, were purchased from Lonza (Walkersville, MD). On week 12 after transplantation, mice were checked for human cell reconstitution by double staining with human FITC-conjugated anti-human CD45 antibody and APC-conjugated anti-mouse CD45 antibody (BD Pharmingen). Mice successfully transplanted (i.e. 20-50 % human CD45+) were chosen for further experiments.

#### Immunoprophylaxis study

NSG-SGM3 or NSG-IL15 mice were injected with various doses of bnAbs or Synagis. Three hours later, mice were bled for measurement of bnAb levels and, immediately after bleeding, challenged with minimum doses of transmitter founder HIV-1 1086c required for 100% infection of control animals. These doses were 30 and 75 TCID_50_ units of a virus stock titrated in primary PBMCs for CD34-NSG-SGM3 and CD34-NSG-IL15 mice, respectively. Plasma HIV RNA was quantified by Taqman qPCR at weekly intervals from Week 1 to week 4.

#### Plasma HIV RNA quantification by TagMan qPCR

Plasma viremia was measured using TaqMan Fast Virus 1-Step master mix (Thermo Fisher) in an automated Step One Plus Real-Time PCR detection system (Bio-Rad). The assay uses the previously described HIV primer pair 6F/84R and Taqman probe [51]. Viral RNA was isolated using Qiagen Viral RNA mini kit. This protocol is intended for isolation of viral RNA from plasma volumes of at least 140 µl. TaqMan qPCR amplification of HIV RNA isolated from 140 µl of patient’s plasma has a limit of detection (LOD) of 40 copies/ml. In humanized mice, available plasma volumes are small, typically ∼ 40 µl, requiring a 3.5 dilution with PBS to reach the minimum 140 µl needed for RNA isolation. Accordingly, the LOD of the TaqMan qPCR assay in mouse plasma samples is ∼ 150 copies/ml.

## Results

### Antibody isolation and primary engineering

As previously reported, we assembled a cohort of PLWH whose plasma demonstrated the presence of bnAbs against HIV [25, 52–54]. One subject (N49) harbored particularly broad and potent plasma neutralizing activity. Such neutralization potency and breadth were maintained in circulation over a 12-year period as shown by testing of temporal plasma samples against a multi-tier panel of pseudoviruses in the TZM-bl assay format (not shown). The N49 plasma sample [25] collected in 2012 neutralized 99% of 117 pseudoviruses comprising tier 1-3 isolates in a global panel. Using a combination of affinity chromatography, isoelectric focusing and mass spectroscopy, we isolated bnAb H and L chain sequences, which were assembled against a B-cell gene database derived from the same subject. This exercise traced broadly neutralizing activity to two families anti-CD4bs bnAbs termed N49P7 and N49P9. Both families use IGHV1-2 heavy chain (subclass IgG1) and λ light chains, with deletions in CDRL1 and CDRL3, but are distinguished by use of different λ light chain families and CDRH3 lengths [25]. Further mining of the N49 B-cell database identified other antibodies with closely matched CDRH3 clonal variants with superior neutralizing activity, including N49P9.3. All wildtype N49 P-series bnAbs carry an amino acid mutation in the first position of the heavy and/or light chain constant regions. This is due to the final position of the J segment determining the first nucleotide of the first amino acid in the constant region. We previously noted that reversion of this mutation to the germline constant region yielded slightly improved neutralization in several N49 P-series bNAbs. Accordingly, a P to A mutation (P1.4A CH1) in the first position of the heavy chain constant region, and a R to G mutation (R1.5G IGLC7) in the first position of the light chain constant region were introduced in N49P9.3, yielding the antibody N49P9.6.

Using a previously reported design strategy [31, 32, 55], N49P9.6 was engrafted with the acidic and elongated heavy chain framework-3 loop of the anti-CD4bs antibody VRC03 (**Figure 1**) to facilitate an additional quaternary interaction with the CD4bs of an adjacent gp120 protomer in the Env trimer [31, 33]. This modification has been shown to both increase the neutralization potency of anti-CD4bs bnAbs and improve the overall biochemical behavior of the antibodies [31, 32, 55]. The resulting variant, N49P9.6-FR, was then subjected to further structural and functional analyses. For comparison, a related variant, N49P9.3-FR, which differs from N49P9.6-FR by having wildtype mutations in the first position of the constant regions, was generated by the same design strategy.

**Figure 1.**
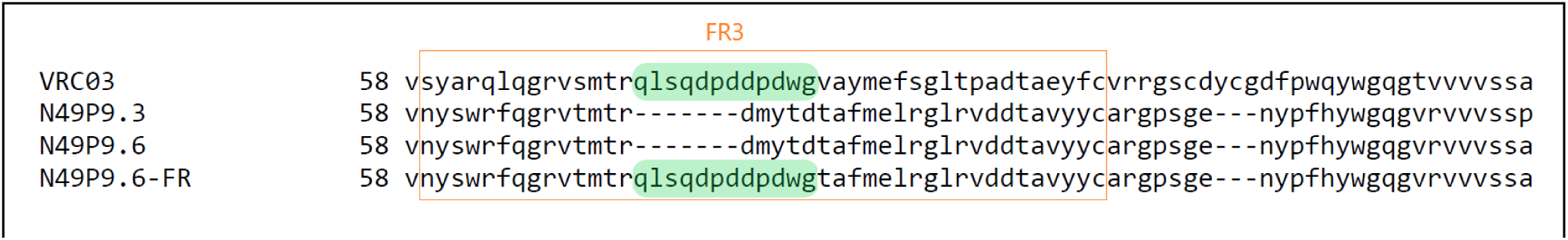
Engraftment of heavy chain framework-3 loop from VRC03 into N49P9.6 to engineer N49P9.6-FR. The heavy chain framework-3 loops of the bnAbs are set within the orange box, and the part of VRC03 engrafted into N49P9.6 is shaded in green.

Sequential engineering from N49P9.3 to N49P9.6-FR yielded incrementally improved neutralization breadth and potency as determined against a global 118 pseudovirus panel representing multiple clades and neutralization tiers (**Figure 2, Figure S1**). N49P9.6-FR demonstrated 97% coverage at a median IC50 potency of 0.01 µg/ml and IC80 potency of 0.03 µg/ml. Overall, the median IC50 value for N49P9.6-FR indicated 5.3-fold and 4.1-fold greater potency compared to N49P9.3 or N49P9.6 (P<.0001 and P<.0001, respectively) (**Figure 2)**. Similarly, the median IC80 values for N49P9.6-FR indicated 5-fold, and ∼2-fold greater potency versus N49P9.3 or N49P9.6 (P<.0001 and P<.0001, respectively).

**Figure 2.**
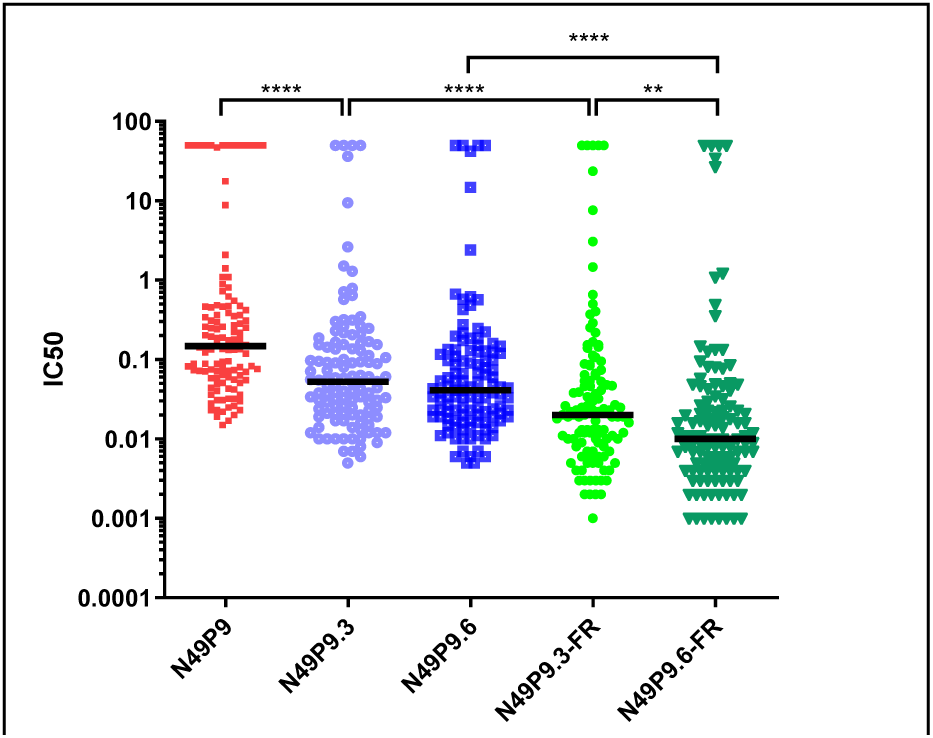
Improvement in neutralization through engineering. Variants of N49P9.3 show lower IC50 values compared to parental forms in 118 multi-tier multi-clade pseudovirus panel. N49P9.6-FR has an IC50 and IC80 of 0.01 and 0.03 μg/ml, respectively. IC50 values between groups compared by Mann-Whitney U test: ** = P<.01; **** = P<.0001

N49P9.6-FR was next compared to other CD4bs, V2 apex and V3 glycan bnAbs for a range of key qualities reflecting breadth and potency. For this purpose, data were either collected or downloaded from the CATNAP database [56], which comprises a global neutralization panel of 96 pseudoviruses (derived from the 118 pseudovirus panel) against which all lead bnAbs have been tested. Comparing IC80 titers for individual bnAbs, N49P9.6-FR had the best geometric mean of 0.069 µg/ml, a 3-fold improvement over the next best score for VRC01.23LS. The AMP trials suggested that an IC80 <1 µg/ml might be required for effective prevention. N49P9.6-FR showed the highest breadth at this stringent cutoff – 88.5% – among all the bnAbs (**Figure 3**, **Table S2**). The AMP trials also suggested that complete neutralization of viruses (as opposed to 80% neutralization) could be required for clinical efficacy. To this end, we predicted IC99 titers for each individual bnAb and found that N49P9.6-FR showed the highest potency and breadth. We also included analysis of the instantaneous inhibition potential (IIP) representing ≥5Log10 reduction in viral replication, which Jilek et al. showed [57] was associated with clinical success of antiviral agents in HIV-1 therapeutic settings. Individually, bnAbs have met this bar for very few HIV variants [30, 58]. IIP predictions estimated N49P9.6-FR to produce the highest median value and the highest breadth, albeit modest, for ≥5Log10 IIP against 14.6% of viruses. Thus, N49P9.6-FR consistently showed superior and best-in-class in vitro neutralization potency and breadth versus other CD4bs bnAbs, including VRC01.23LS, VRC07-523LS, N6, and 1-18 (**Figure 3**, **Table S1**).

**Figure 3.**
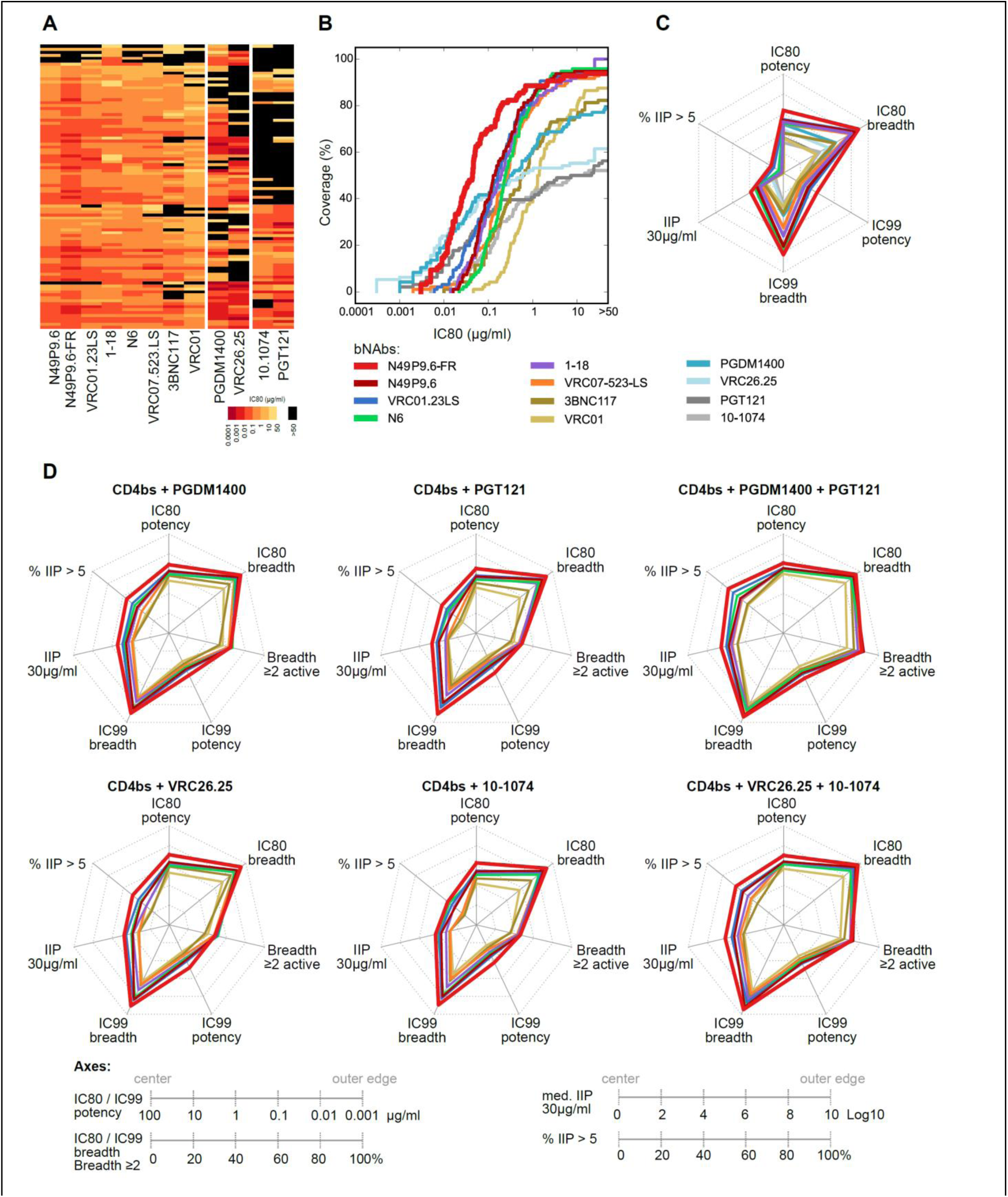
N49P9.6-FR outperforms other bnAbs individually and as part of bnAb combinations. (A) Heatmap of IC80 titers for N49P9.6, N49P9.6 FR and other best-in-class CD4bs, V2 apex and V3 glycan bnAbs. (B) IC80 breadth-potency curves for bnAbs in (A). (C) Spiderplots showing multimetric comparison of bnAbs using IC80 breadth & potency, IC99 breadth & potency and IIP median and breadth (IIP > 5Log10). Axes ticks are defined in legend in (D), and for each metric, radially outwards is better performance. (D) Similar to (C) except evaluating 2– and 3-bnAb combinations. Each CD4bs bnAb is combined with V2, V3 or both bnAbs, and IC80, IC99 and IIP are predicted.

While N49P9.6-FR exhibited a neutralization breadth and potency on par with the best-in-class bnAbs, we do note that there are viruses in this panel that are poorly neutralized or resistant to N49P9.6-FR, and the mutations that confer this resistance could be present in the large viral reservoirs in PLWH. One of the common strategies to address resistant viruses is the use of cocktails of bnAbs that recognize different epitopes on Env and are susceptible to different escape mutations. Thus, it was important to determine the functionality of N49P9.6-FR in combination with other bnAb classes. Several previous studies showed that such combinations significantly improve neutralization potency and breadth, particularly with respect to IIP, as needed for clinical success [30, 58–60]. Accordingly, we predicted neutralization data for 2– and 3-bnAb combinations in our dataset using a Bliss-Hill model [30], and systematically compared them using the above IC80, IC99 and IIP metrics. In addition, we also included coverage with 2 or more bnAbs active (able to neutralize), as this would theoretically improve coverage of the high within-host diversity that is present in each HIV-1 infected person. When combined with the same V2, V3 or both bnAbs (**Figure 3**, **Table S2 and S3**), N49P9.6-FR outperformed all other CD4bs bnAbs. The best 2-bnAb combination was N49P9.6-FR + PGDM1400 and best 3-bnAb combination was N49P9.6-FR + PGDM1400 + PGT121. The latter was predicted to neutralize 94.8% viruses at IC80 < 1 µg/ml (total bnAb concentration, not individual), 93.8% at IC99 < 10 µg/ml, and 71.9% at IIP > 5Log10; thus, highlighting the advantage of combining higher number of bnAbs. These results together suggest that N49P9.6-FR is an excellent candidate for clinical testing either individually or as part of a bnAb combination for HIV-1 prevention and/or therapy.

Several bnAbs, including those in the anti-CD4bs class, demonstrate variable levels of autoreactivity and polyreactivity [61, 62], which may confound vivo pharmacokinetics and clinical testing. Accordingly, N49P9.6-FR was tested for autoreactivity and polyreactivity against a standard clinical panel of human antigens including ANA, rheumatoid factor, centromere, Jo1, SSA, SSB, SCL-70, RNP, and SM. N49P9.6-FR tested negative for all antigens.

To increase the antibody in vivo half-life, we exploited a strategy of introducing two mutations, M428L and N434S, into the Fc domain of IgG1. These mutations were shown to increase Fc-binding affinity for the neonatal Fc receptor (FcRn) and increase plasma half-life [63, 64]. The resulting variant, N49P9.6-FR-LS, had a neutralization profile like its parental strain, N49P9.6-FR (less than a 2-fold difference in median IC50 and IC80) (**Figure S1**). Importantly, N49P9.6-FR-LS had 100% breadth when tested against a 60 pseudovirus panel from the placebo group from the AMP trial (Clades C, A1/B, B/F, F1/B, F, F1, F2) (**Table S4**), with an IC50 of 0.047 ug/ml, and IC80 of 0.164 ug/ml. Both forms were able to mediate potent ADCC against a panel of infectious molecular clones encoding recently circulating Envelopes sampled from placebo infections in the phase II/III trials, HVTN703/HVTN704 and HVTN705 (**Figure S2**). N49P9.6-FR-LS was then tested for plasma half-life in hFcRn mice, which harbor a knockout allele of the FcRn α-chain (Fcgrt^tm1Dcr^), express the human FcRn α-chain, and are extensively used for measuring *t*_1/2_ of human mAbs and Fc-engineered mutants [65]. Importantly, the *t*_1/2_ in this model strongly correlates with *t*_1/2_ in humans (superior to NHP, hemizygous hFcRn transgenic mice, and wildtype mouse models) [65]. As shown in **Table 1**, both the –FR insertion and the LS mutations lengthened the antibody half-life, as previously reported [32]. N49P9.6-FR-LS had a longer half-life in FcRn mice than any of the control CD4bs bnAbs tested, including VRC01-LS (**Table 1**).

**Table 1:**
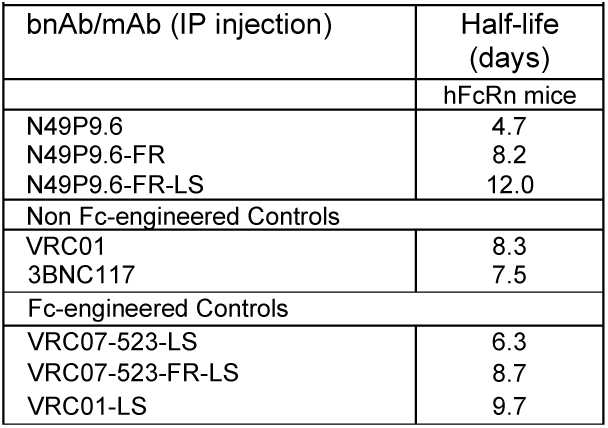
Estimated half-life by mAb by non-compartmental analyses (NCA)

| Table 1: Estimated half-life by mAb by non-compartmental analyses (NCA) |  |
| --- | --- |
| bnAb/mAb (IP injection) | Half-life (days) |
|  | hFcRn mice |
| N49P9.6 | 4.7 |
| N49P9.6-FR | 8.2 |
| N49P9.6-FR-LS | 12.0 |
| Non Fc-engineered Controls |  |
| VRC01 | 8.3 |
| 3BNC117 | 7.5 |
| Fc-engineered Controls |  |
| VRC07-523-LS | 6.3 |
| VRC07-523-FR-LS | 8.7 |
| VRC01-LS | 9.7 |

### Antiviral efficacy in vivo

Analyses of bnAb protective efficacy were carried out using NSG-SGM3 and NSG-IL-15 mice reconstituted with human CD34+ stem cells. One model (HuCD34 NSG-IL-15) efficiently reconstitutes human T cells as well as NK cells [66, 67], thus allowing any bnAb-triggered Fc-dependent effector functions to occur along with Fab-mediated Env inactivation. Several studies indicate that Fc-mediated effector functions contribute to bnAb efficacy in animal models providing effector cells [17, 66]. A recently published study in reconstituted MISTRG-IL-15 mice highlighted how NK cells control HIV infection in part by utilizing bnAbs that enable ADCC [17]. To provide a rigorous test of efficacy, we used a clade C transmitted/founder virus (1086c) with moderate sensitivity to N49P9.6-FR-LS (IC80 of 1.6 µg/ml in the TZM.bl neutralization assay format; data not shown). Serial dilutions of bnAb were administered IP. The anti-RSV antibody, Synagis, served as negative control, and was administered at a dose matching the highest bnAb concentration (20 mg/kg). After 3 hours, plasma was collected to measure circulating bnAb levels, and mice were challenged IP with a dose of challenge stock previously determined to infect all untreated Hu-CD34-NSG and Hu-CD34-IL15 mice (30 and 75 TCID_50_, respectively). At one week before challenge, the fraction of human CD45+, CD56+ cells in the NSG-IL-15 mice ranged from ∼3-31% with a median of 11.1%, while in the NSG-SGM3 mice they ranged from 0.07-13% with a median of 0.7%. The CD4+/CD8+ T cell ratios in the NSG-IL-15 mice ranged from ∼0.4-6 with a median value of 2.1 and the NSG-SGM3 mice ranged from ∼1.6-14 with a median of 5.1. To secure enough NSG-SGM3 and NSG-IL-15 mice for the experiment, cohorts of mice with different human CD34+ cell donors were combined. There were statistically significant differences between cohorts for the CD45, CD4 and NK cell populations. However, in each experiment, the cohorts were equally distributed amongst the treatment groups, and those groups lacked significant differences in the cell populations.

Animals were tested for serial plasma viral loads post-challenge for 28 days (**Figure 4A).** Treated animals were scored as uninfected if plasma viremia was undetectable over one-month follow-up period post-challenge. The dose-response curve (**Figure 4B**) showed that 63% protection was achieved in Hu-CD34-SGM3 mice at the 20 mg/kg dose. In contrast, 100% protection was evident in the Hu-CD34-IL15 mice at 20mg/kg, with 72% protection seen at a dose of 1.25mg/kg **(Figure 4C)**. All Synagis treated animals were infected. As shown in **Figure 4C**, the 1.25 mg/kg dose that produced 72% protection achieved detectable plasma bnAb levels averaging ∼ 4 µg/ml at the time of challenge. The higher plasma concentrations (≥ 10 µg/ml) for the higher bnAb doses were associated with > 85% protected animals. Notably, this dose approximates a plasma IIP value of ∼3.6 using measures from single round infection assays as described [57, 68–70].

**Figure 4.**
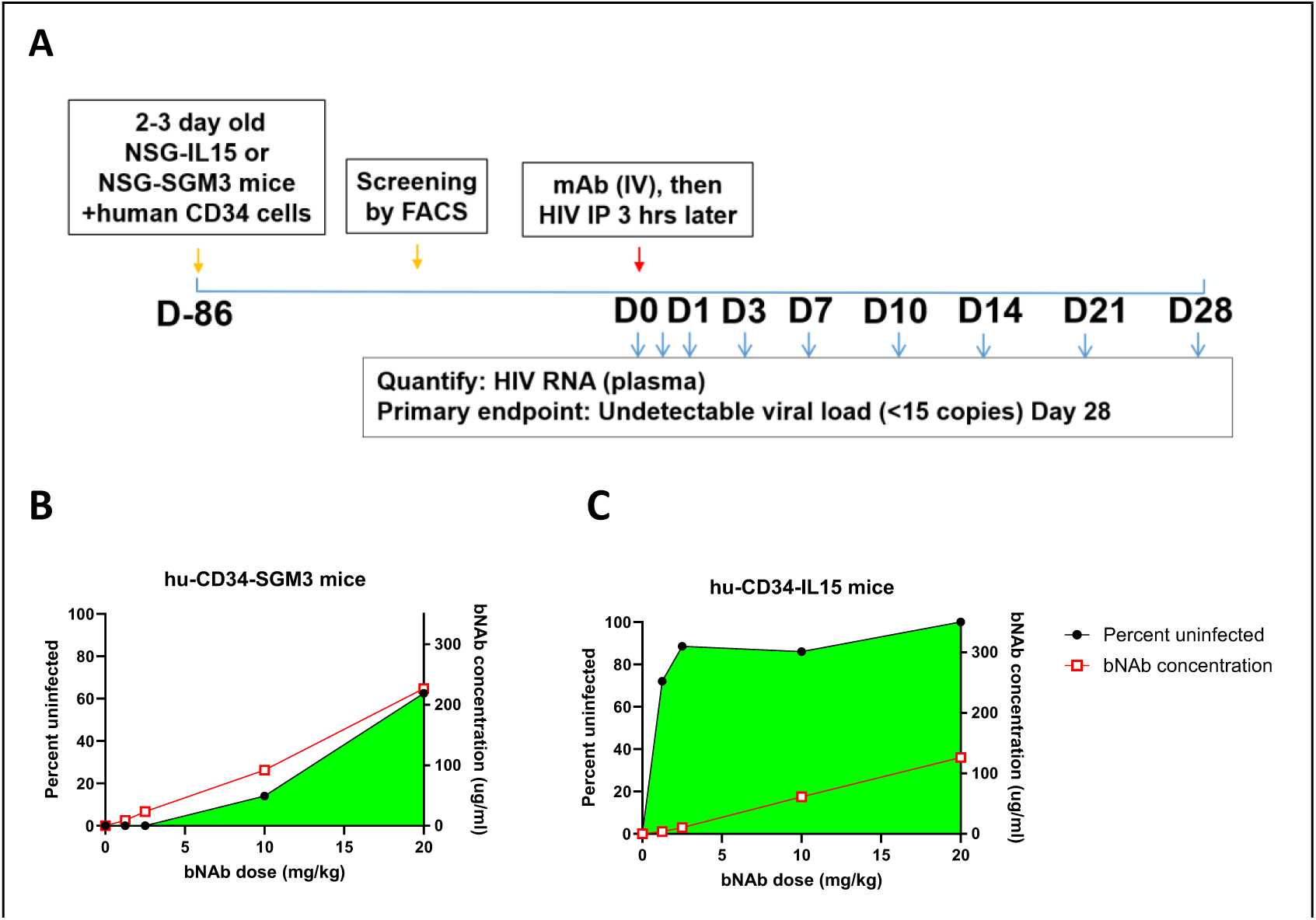
**Comparison of N49P9.6-FR-LS potency in prevention of HIV acquisition in Hu-CD34-SGM3 and Hu-CD34-IL15 mouse models**. (A) Mice were injected IV with the indicated doses of bnAb and 3 h later challenged with HIV 1086c IP. Plasma viremia was measured by RT-qPCR weekly for 4 weeks to determine infection status. (B) In SGM3 mice maximum protection of 63% was seen only at the highest bnAb dose tested (20mg/kg). (C) In IL15 mice protection was seen at all bnAb doses tested from 72% (1.25mg/kg dose) to 100% (20mg/kg dose). Immediately before virus challenge, mice were bled to determine bnAb plasma concentrations (ELISA). In each group mean plasma titers were matched to the level of protection. All control mice, injected with Synagis, became infected in both mouse strains (not shown). The experiment was repeated with similar results.

### Structural analyses

To define the molecular basis for N49P9.6-FR and the N49 P9 family of antibodies in binding to the HIV Env, we solved crystal structures of N49P9.1, N49P9.3 and N49P9.3-FR Fabs in complex with clade A/E gp120_93TH057_core_e_. Analysis of the structures reveal that only two gp120 contact residues differ between N49P9.1 and N49P9.3, one in CDRH1 and one in CDRH2 (**Figure S3, Table S5**). The first, Phe^33^ in CDRH1, packs against gp120 Ala^281^ in place of Leu^33^ in N49P9.1, but with slightly better van der Waals contacts, and modulates heavy chain Trp^50^, which also packs against gp120 Ala^281^ and forms a hydrogen bond to gp120 Asn^280^ (**Figure S3 B and C**). The second Gln^64^ in CDRH2 can form hydrogen bonds with both gp120 Arg^469^ and Asp^457^ versus Glu^64^ in N49P9.1 which can only form a hydrogen bond with Arg^469^. Gln at position 64 is also what is seen in both N49P7 and VRC01 implying that Gln can make an important contribution to binding when placed here [25, 71]. These two amino acid changes likely account for much of N49P9.3’s increase in neutralization potency and breadth relative to N49P9.1.

To interpret the above improvements and possibly inform additional refinements, we resolved the structure of N49P9.6-FR Fab in complex with a BG505 SOSIP.664 trimer by cryo-electron microscopy (cryo-EM) with Fab of PGT121 as a chaperone. A 4.02 Å experimental density map was generated using data collected from two different types of cryo-EM grids (**Figure 5**, **Figure S4-S5 and Table S6**). Refinements were informed by crystal structures of an N49P9.3-FR Fab and Fabs of other N49P9 linages in complexes with the gp120_93TH057_ core_e_ (**Figure S3**, **Table S5**). This operation was informative as N49P9.6 has only two amino acid differences outside of the variable region, relative to N49P9.3, and is nearly identical in both the N49P9.6-FR Fab-PGT121-BG505 SOSIP.664 and N49P9.3-FR Fab-gp120_93TH057_core_e_ complexes, with most contacts to the primary protomer in the trimer matching those observed in binding to the gp120 monomer (**Figure 5D** and **Figures S3** and **S5**). The only exception is an additional contact of heavy chain framework-3 residue Gln^75^ of N49P9.6-FR to the glycan on Asn^197^ of gp120 seen only in the trimer complex (**Figure 5A**). A similar contact was previously observed in the complex of N49P9.6 Fab with BG505 SOSIP.664, where framework Asp^72^ of N49P9.6 contacted the gp120 glycan on Asn^197^ [72]. Asn^197^ is absent in the gp120_93TH057_core_e_ complexes because of the V1V2 loop truncation. Other minor differences in contacts between the gp120 core and SOSIP structures are due to lack of sequence identity at six contact residues (371, 430, 460-461, and 474-475) (**Figure 5D** and **S3B**). However the changes are conservative in nature and do not disrupt N49P9.6-FR binding. The total buried surface area (BSA) for the N49P9.3-FR Fab-gp120_93TH057_core_e_ structure is 2491 Å^2^ (1254 Å^2^ for gp120 and 1237 Å^2^ for fab) while the total BSA for the primary gp120 interface in the N49P9.6-FR-BG505 SOSIP.664 complex is 2441 Å^2^ (1184 Å^2^ for gp120 and 1257 Å^2^ for Fab).

**Figure 5.**
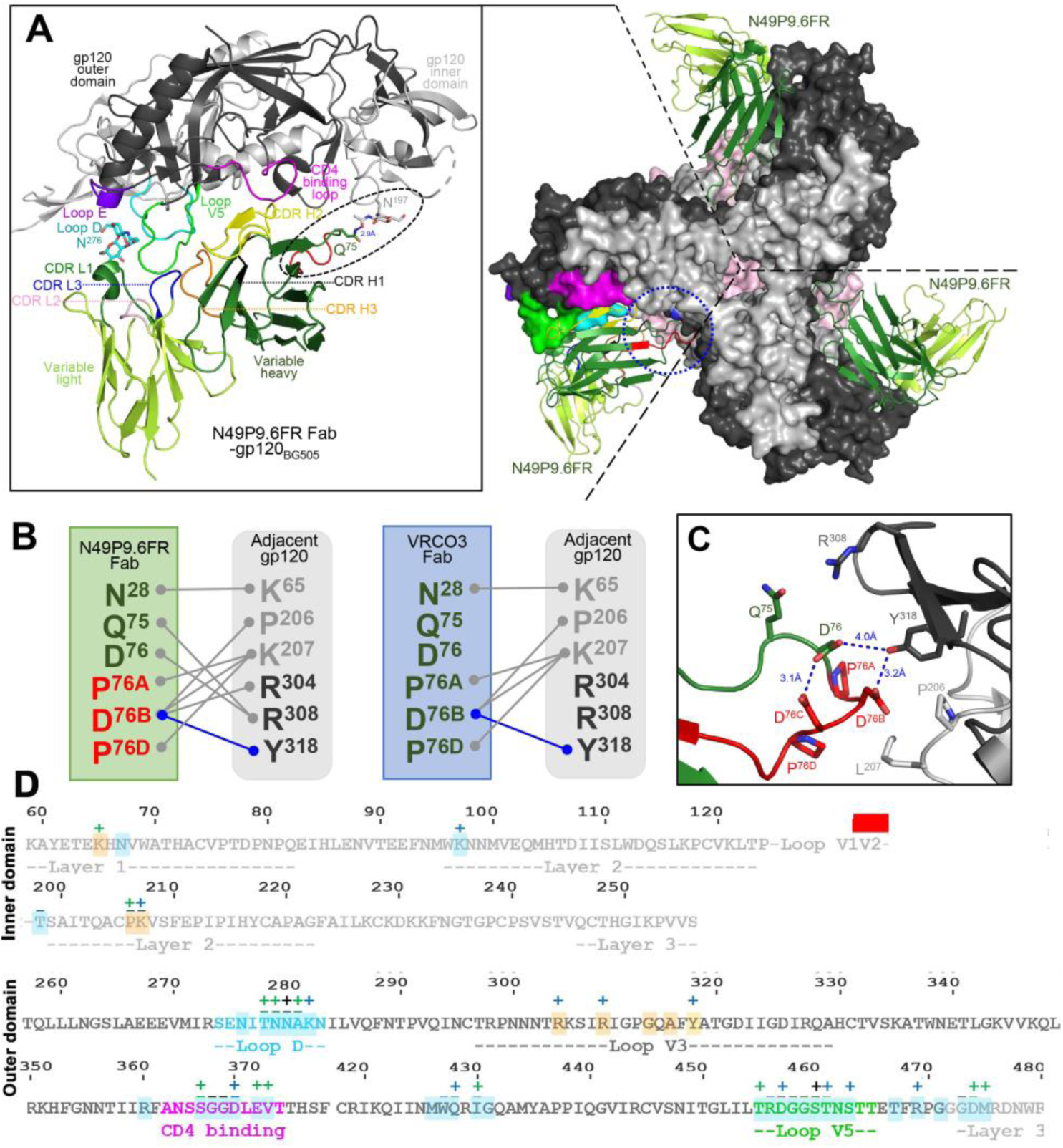
**Molecular details of the Env trimer interprotomer contacts of N49P9.6-FR**. (**A**) Top view of the complex of BG505 SOSIP.664 HIV-1 Env trimer with N49P9.6-FR (the Fabs of PGT121 were omitted for clarity). The three N49P9.6-FR Fabs (the Cryo-EM model was built with the variable regions of the Fabs only, dark and light green ribbon) are shown bound to Env trimer (inner and outer gp120 domains colored light and dark gray, respectively). On the right, one N49P9.6-FR-gp120BG505 complex was extracted from the trimer to show the details of the Fab contacts to the primary gp120 protomer. One additional primary gp120 contact of N49P9.6-FR that is only seen in the Env trimer that is mediated by the region proceeding the introduced frame sequence is encircled. On the bottom, a zoomed-in view depicts the contact between N49P9.6-FR with the adjacent BG505 protomer. All contact residues within a 4-Å distance criterion cut-off are shown as sticks and H-bonds are shown as blue dotted lines. The residues of the introduced frame sequence are shown in red. (**B**) Network of interactions formed between N49P9.6-FR with the adjacent gp120BG505 protomer as defined by a 5-Å distance criterion cut-off. The contacts for VRC03 by the same criteria to the adjacent gp120BG505 protomer are shown for comparison (PDB ID: 6CDI). (**C**) A zoomed-in view depicts the contact between N49P9.6-FR with the adjacent BG505 protomer. All contact residues within a 4-Å distance criterion cut-off are shown as sticks and H-bonds are shown as blue dotted lines. The residues of the introduced frame sequence are shown in red. (**D**) Epitope footprints of Fabs mapped onto the gp120 primary sequences. Contact residues are defined by a 5 Å cutoff and marked above the sequence with (+) for side chain and (-) for main chain to indicate the type of contact. Contact types are colored as following: hydrophilic (blue), hydrophobic (green) and both (black). Buried surface residues determined by PISA are shaded blue for primary gp120 contact residues and orange for adjacent gp120 contact residues.

Although, as described above, the primary N49P9.6-FR-gp120 interface closely resembles those observed in the N49P9.3-FR-gp120 monomer and N49P9.6 Fab-SOSIP trimer complexes, the N49P9.6-FR Fab establishes a new set of interactions with the adjacent gp120 protomer (**Figure 5B** and **C**).These contacts are mediated by the VRC03 heavy chain framework-3 insertion in N49P9.6-FR and are similar, though not identical, to the network of interactions previously reported to promote greater potency in other CD4bs bnAbs including VRC03 [31, 32]. Total BSA for the adjacent protomer is 388 Å^2^ for N49P9.6-FR (194 Å^2^ for gp120 and 194 Å^2^ for Fab) which is larger than that observed previously for VRCO3 of 376 Å^2^ (179 Å^2^ for gp120 and 197 Å^2^ for Fab), N6-FR of 298 Å^2^ (138 Å^2^ for gp120 and 160 Å^2^ for Fab), and VRC01-FR of 385 Å^2^ (180 Å^2^ for gp120 and 205 Å^2^ for Fab). N49P9.6-FR primarily makes one hydrogen bond to gp120 Tyr^318^, weak salt bridges with Lys^207^, Arg^304^, and Arg^308^, and van der Waals contacts to Pro^206^ on the adjacent protomer (**Figure 5B** and **C**). VRC03 makes similar contacts to the adjacent protomer minus contacts to Arg^304^ and Arg^308^. N6-FR shifts its adjacent protomer contacts more toward Tyr^318^ while VRC01-FR shifts more towards Lys^207^ and Arg^304^. The additional adjacent protomer contacts uniquely established by N49P9.6-FR adds approximately 16% more BSA versus the related bnAbs. Notably, N49P9.6-FR’s opening of the trimer, as determined by the average rotation (**Table S7**) is distinct and intermediate between VRC03 (more open) and N6-FR or VRC01-FR (more closed). Overall, the heavy chian framework-3 insertion in N49P9.6 better mirrors VRC03 than the same insertion engrafted onto N6 or VRC01.

### Secondary Antibody Engineering for Potency

Further engineering of N49P9.6-FR-LS to improve neutralizing activity was attempted using structure-based design tools. Based in part on previous successful HIV antibody design efforts [25, 26], we assessed germline revertant mutations and consensus residue mutations from VRC01-class antibodies. Analysis of designs was performed using the Therapeutic Antibody Profiler (TAP) tool [27], which assesses antibody developability using structure-based metrics, as well as Rosetta [28], which was used to calculate antibody stability or binding energy changes (ΔΔGs) for individual substitutions based on modeling. Favorable substitutions identified by one or both of those methods were prioritized for experimental testing. We additionally tested a substitution to remove a possible N-glycosylation motif in the heavy chain V domain (S60A), given that N-glycan removal was previously shown to be beneficial for antibodies [29], as well as a CDRL2 loop residue swap (FDDK49YSGST) from the related VRC07 antibody [25]. Finally, two substitutions were selected based on computational scanning mutagenesis of the heavy chain framework-3 loop in Rosetta and favorable predicted Env targeting and/or thermostability. In total, 11 N49P9.6-FR designs were selected for characterization (**Table S8**).

All new designs were expressed by transient transfection (see Methods) and assessed for neutralizing activity (breadth and IC50, IC80 potencies) using a screening mini-panel of 21 pseudoviruses (extracted from the larger 119 global panel) comprising viruses that are moderately or completely resistant to the parental N49P9.6-FR-LS. None of the changes led to a statistically significant improvement in the IC50 or IC80. However, certain modifications led to improved neutralization against specific viruses in the mini-panel (>5 fold improvement in the IC50 and/or IC80), including the N-glycan site removal substitution, variable-constant domain hinge substitutions (heavy chain V110T and light chain T108Q), and a structure-based point substitution (light chain T70D). However, overall improvements in median potency were not significant when the modified constructs were tested on the larger 119 pseudovirus panel (not shown). Interestingly, some designs, particularly the two “YSGST” CDRL2 swap-containing variants, which included CDRL2 residues from another VRC01-class antibody (VRC07), showed substantially reduced breadth and potency versus N49P9.6-FR. Overall, the secondary engineering yielded no further improvements in breadth or potency in the assays we employed.

### Secondary Antibody Engineering for Developability

A parallel effort focused on identifying modifications in N49P9.6-FR-LS that potentially improve scalability and manufacturability under cGMP production, without loss of neutralizing activity or other desirable properties. Given the PK data above, which demonstrably increases circulating half-life in vivo (**Table 1**), all variants expressed a heavy chain sequence containing the “LS” mutation.

The first round of optimization was performed in a phage library panning format based on efficient HIV CH505 gp160 binding. Sixty-two candidates were identified during phage-Fab panning that bound at a 0.8nM target concentration after either temperature or chemical challenge (see methods). These variants were produced in a transient expression system and tested for retention of neutralization breadth and potency versus the parental constructs. Detrimental mutations were defined as ones with > 5 fold lower BaL.26 EC50 gp120 binding on ELISA and/or≥ 5 fold increase in IC50 or IC80 against any 1 virus in the 10 pseudovirus panel (extracted from the larger 119 global panel) Eleven candidates demonstrated unacceptable loss of neutralization; one of these also had > 5-fold decrease in binding by EC50.

Of the remaining 51 variants with fully preserved breadth and acceptable potency, 5 showed 2-4-fold median improvements in IC50 and IC80. The 51 variants were then further screened (see Methods) to assess conformational and colloidal stability characteristics (**Figure S7**) favoring manufacturability properties [73–75]. The variants were ranked according to each of the parameters and the variant with the most improved measure in each category (without worsening in another category) was chosen for further screening.

This exercise identified 6 variants exhibiting improved conformational and/or colloidal stability. These variants were then combined with the 5 that had demonstrated more potent neutralization for further screening. Two variants demonstrated both the conformational/colloidal and neutralization improvement (they were found in both sets), meaning a total of 9 variants underwent further screening. Stable pools were made from these 9 variants and re-tested for neutralizing activity against 60 pseudovirus panel (one half of the 119 Global Tier 1-3 panel), and conformational/colloidal properties. The neutralization IC50 and IC80 were all within 2-fold of the parental N49P9.6-FR-LS strain, while the only measures with improvement seen between the transient transfection and stable pool material was in PEG solubility and thermal hold in one variant (M-5817), and PEG solubility and T1 in a second variant (M-5809) (data not shown). Thus, the apparent neutralization gains seen with the 10 virus panel were not reflected in a larger panel.

One of the liabilities of the antibody that we identified during analysis with RANDŸ was the 3 amino acid deletion in N-terminus of the light chain. This is a common property in CD4bs bNAbs, seen in all N49 P series bNAbs and VRC01, for example [25, 55]. Expressed as recombinant proteins, the N49 P series light chains lack 2-3 amino acids encoded by the corresponding N terminal nucleotide sequences. Mutations in the leader sequence result in the truncated N-terminus in the light chain as determined by Edman degradation N-terminal sequencing and mass spectrometry of light chain enzymatic fragments (data not shown). Such data indicate post-translational enzymatic processing. Accordingly, one optimization strategy was to include the initial “ASA” sequence in the light chain N-terminus along with a leader sequence that would not cleave it. Variants containing this modification were given the germline or “_GL” designation. In the final phase of development, wildtype and “-GL” forms of M5809 and M-5917, as well as parental N49P9.6-FR-LS were each made.

These six variants underwent another round of neutralization testing (119 global panel and a 100 Clade C virus panel). These analyses again showed IC50 and IC80 that were comparable among all variants and the parental bNAb (<2-fold differences), although one was statistically significant (**Figure S8**). These six variants underwent another round of biophysical testing, and based on improvements in thermostability, PEG solubility, and viscosity compared to the parental form N49P9.6-FR-LS, variant M-5817_GL demonstrated the broadest improvement overall **(Table S9)**. Thermal stability of M-5817_GL was improved as evinced by the additional melting transition at 70°C which corresponded to a reduction in precipitation upon thermal hold **(Table S9)**. While the conformational stability was improved in the thermal stress methods, a slight reduction in the inflection point during chemical unfolding was observed **(Table S9)**. Viscosity curves (**Figure S9)** were generated for parental N49P9.6-FR-LS and M-5817_GL to validate that improvements in colloidal stability were also evident under GMP-relevant concentrations and formulation conditions (**Table S9)**. As indicated by the improved colloidal stability measured by biophysical analysis, the optimized candidate showed a significant reduction in viscosity when formulated at high concentrations. M-5817_GL (henceforth referred to as eN49P9) was the final candidate chosen for further clinical development.

## Discussion

Over the past decade, there has been an extensive effort to discover “pan neutralizing” anti-Env bnAbs arising in humans during HIV infection, based on the premise that such antibodies might effectively treat or prevent infection in special settings wherein conventional ARVs are suboptimal and/or where functional cure is a goal. Yet clinical experience such as the AMP trials [18] has shown that individual bnAbs suffer the same shortcomings as all countermeasures based on single antiviral agents; specifically, viral escape due to HIV genomic diversity and mutability. Taking lessons from the use of conventional ARVs, one approach to overcome this issue is to combine bnAbs from different specificity classes, each with nonoverlapping gaps in variant coverage [58, 60, 76]. Another approach is to engineer select, near pan-neutralizing bnAbs toward enhanced breadth and potency. It stands to reason that a combination of ebnAb specificities would provide the strongest prospects for clinical efficacy.

Any rational enhancement of bnAb function must strike a balance between increasing breadth and potency as part of an aggregate profile of stability, acceptable in vivo PK and, importantly, manufacturability under cGMP. Some of these qualities may not be selected during affinity maturation, but are of great importance in preclinical and clinical development. During engineering, improvements in one property can lead to deterioration in another. The CD4bs bNAbs are particularly challenging as they acquire their properties in part via high rates of somatic hypermutation, with over 40% change from germline being reported [25, 77, 78]. Mutations are not restricted to the CDRs, but seen in all framework regions, and even extend to the 1^st^ position of the constant region, as seen in the N49 family of bNAbs. VRC01-class antibodies have a characteristic deletion in the CDRL3 [79]. In addition to this, deletions in the N-terminus have been noted [55]. These sequences create distinct Fab architectures that enable broad recognition of Env variants via main chain and sidechain interactions [25, 77, 80]. Thus, anti-CD4bs antibodies are difficult to engineer; for example, recognized techniques for increasing potency such as a 54W/F/Y substitution in the heavy chain can lead to polyreactivity [55].

Recognizing these hurdles, we pursued a stepwise, comprehensive approach towards engineering of a near pan-neutralizing, plasma anti-CD4bs bnAb, N49P9.3. This approach yielded an enhanced variant, eN49P9 capturing exceptional breadth and potency qualities, desirable PK behavior, and biophysical properties amenable to GMP production. As noted above, such an aggregate profile holds superior potential for clinical applications.

Our starting resource was a group of bnAbs recovered from donor N49 plasma as part of an anti-Env repertoire comprising two “N49P families” with anti-CD4bs specificity [25]. All family members have distinguishing features and differ from the canonical bnAb of this class, VRC01. The N49P9 family is similar to VRC01 in that it has a similar length CDRH3. But unlike VRC01 it binds the CD4bs cavity of the Env trimer through an interaction between 54Y in CDRH2 and the Phe43 cavity in gp120, more like N6. In comparison, the N49P7 family has a more elongated CDRH3, which bypasses the Phe43 cavity, making functional contacts with conserved residues in the gp120 inner domain [25]. N49P9.3 variant demonstrated superior breadth and potency, as a potential consequence of establishing unique contacts with gp120 (**Figure S3**) not seen in other family members. Accordingly, N49P9.3 was engineered to produce N49P9.6 by reverting the 1^st^ position of the constant regions in the heavy and light chains to germline, which was then engrafted with the framework-3 loop from VRC03 to enable quaternary interaction with a second gp120 protomer, a strategy previously shown to significantly improve the potency and pharmacokinetics of several CD4bs bnAbs (26,27). Notably, this insertion led to substantial gains in neutralizing potency of N49P9.6, even greater than for N49P7-FR or VRC01.23LS [55]. Thus, the engraftment strategy particularly benefits the N49P families of bnAbs, likely because the resulting structures experience peculiar avidity gains from multi-paratope attachment to Env trimers. One reason may be that, as indicated by the cryoEM structures of Fabs (**Figure 5**), in the N49P9.6-FR structure the VRC03 framework-3 insertion contacts the adjacent gp120 protomer in an orientation that more closely resembles VRC03 itself ([32]) allowing for more extensive protomer contacts compared to N6-FR or VRC01-FR. The potential increases in binding avidity offered by such multi-paratope trimer binding may compensate for any paucity in inter-spike crosslinking caused by low Env trimer density on virion surfaces [81–90]. We note that the available literature is equivocal regarding this issue with respect to native virions. Optical techniques indicate localized, tight clustering of trimers while EM suggests more uniformly dispersed positioning [82, 84–87, 90]. Notably, one study based on optical techniques indicated that soluble CD4 or bnAb binding to Env promotes the dispersal of adjacent trimers [88]. Regardless of the scenario, it can be envisioned that increased bnAb-trimer binding avidity would enhance functional potential.

It was encouraging that modifications leading to improved neutralizing activity also boosted ADCC against cells infected by contemporary viruses encoding Env collected from the AMP trial (**Figure S2**). Importantly, these viruses were not genetically modified to include reporter genes that might alter the functionality of HIV regulatory genes. Such findings indicate that enhanced Fab targeting also augments the activities of FcR-expressing effector cells against infected cells. However, the LS mutation, which offers clear advantages for improving PK (**Table 1)** appeared to further modulate the functional activity, lowering or increasing ADCC scores, depending on the variant. The basis for such modulation merits further analyses at the structural level and suggests that trade-offs between ADCC and PK may be required.

The involvement of Fc-dependent effector activities in N49P9.6-FR-LS efficacy was further indicated by our in vivo studies (**Figure 4**), which deliberately used a challenge virus with low in vitro neutralization sensitivity to N49P9.6-FR-LS. Applying PT80 calculations developed for human trials [91], the in vitro neutralization data and NSG mouse plasma levels of N49P9.6-FR-LS indicate that titers of ∼ 317 µg/ml would be needed for 90% protection, assuming neutralizing activity as the principal bnAb function. In accordance, in studies using the hu-CD34-NSG-SGM3 mouse model, in which human NK-cell reconstitution in blood is not robust [92, 93] and neutralization is the predominant mechanism of protection, only modest efficacy (63% uninfected animals) against 1086c was seen at the highest dose (20mg/kg) where mean plasma titers were 226 µg/ml (**Figure 4A**).

In comparison, in the hu-CD34-IL15 model, where reconstituted animals routinely present 5-10 times higher levels of mature human NK cells versus NSG-SGM3 animals [67, 94] we observed substantially better efficacy (>85% uninfected animals) at much lower bnAb doses and corresponding plasma titers of 10 µg/ml, which notably does not meet the optimal neutralization IIP of 5Log10 for the challenge virus. These data indicate that for certain bnAbs such as N49P9.6-FR-LS, in vivo potency may be higher than what is solely predicted by in vitro neutralizing activity and related PT80 calculations. Of course, a true test of this promising concept awaits clinical comparisons of N49P9.6-FR-LS versus other bnAbs.

The neutralizing activity of bnAb class combinations containing N49P9.6-FR-LS was an important parameter for analysis. The field is actively pursuing such combinations for clinical use, hoping to capture the cumulative effects of polyspecific reactivity and overcome virus variability and escape. Currently, triple bnAb combinations seem to offer the best balance between feasibility, cost, and antiviral potential. Cell line-based neutralization assays with individual Env pseudoviruses already show that equimolar 2-class combinations are superior to single bnAbs with respect to breadth and potency [30, 57, 58, 95]; triple or quadruple combinations were superior to ones using two bnAbs. Multi-parameter neutralization measures were uniformly superior when N49P9.6-FR-LS was used alone or in combination as the anti-CD4bs class component (**Figure 3**), outperforming other CD4bs bNAbs evaluated in clinical trials, such as 3BNC117, N6, VRC07523-LS, VRC01.23.LS, and 1-18. Of note, N49P9.6-FR + PGDM1400 + PGT121 exceeded IIP >5Log10 against more than 70% of test viruses. Such potency is likely to be improved using combinations containing rationally enhanced versions of other bnAb classes currently under development.

A desirable scenario for combined bnAb action occurs when more than one antibody acts on the same HIV virion. In the alternative case where each bnAb binds a distinct subset of a virus swarm, escape from only one antibody could allow unbridled replication with no fallback protection. Two or more bnAbs concurrently/redundantly binding virions mitigate this risk and could also exercise cooperativity or synergy [30, 58, 95]. Therefore, it is encouraging that the N49P9.6-FR + PGDM1400 + PGT121 combination was predicted to cover > 80% of test viruses with ≥ 2 bnAbs active. In accordance, using fluorescence correlation spectroscopy (FCS) for direct virion analyses, we previously reported that anti-CD4bs and anti-V3 bnAbs bind to the same particle [96]. More recently, we used FCS and FCS FRET to detect concurrent binding of anti-CD4bs, anti-V3 and anti-V2 apex antibodies to virus particles and even single virion trimers (not shown). The latter indicates an especially advantageous setting for bnAb action that merits further evaluation.

It was intriguing, albeit disappointing, that we were unable to further improve the overall neutralization profile of N49P9.6-FR-LS via structure-guided engineering. While certain modifications led to more potent neutralization of a few variants, they reduced potency against others (**Figure S6**). One limitation of our engineering exercise was the reliance on available SOSIP trimer structures, which represent variants that may already be sensitive to potent bnAb action and thus less capable of highlighting areas for improvement. A second limitation of the approach, which relies on relatively static structures, is that it cannot capture thermodynamic features that might impact bnAb binding and function. For example, it was recently shown that anti-CD4bs bnAbs variably favor open versus closed or intermediate Env transition states [97]. Conversely, the extreme breadth and potency of bnAb N6 appeared to stem from its ability to non-selectively bind across these structures. The equal breadth and greater potency of N49P9.6-FR-LS indicates a similar nonselective binding that is accommodated by the framework 3 engraftment. At this point, it is unclear whether rare variants that are resistant to N6 and N49P9.6-FR-LS have unusual conformational features, sequences, or both. In any case, future structure-guided engineering of bnAbs may benefit from analyses that include additional thermodynamic measures, multiple intermediate forms of trimers, and/or targets based on resistant Env variants.

Our second engineering effort, which focused on enhancing manufacturability while retaining neutralization potency and breadth, was more productive. After extensive screening, one variant (eN49P9) with 6 heavy chain and 3 light chain mutations, as well as an uncleaved light chain N-terminus was identified that improved PEG solubility and thermostability while retaining other desirable biophysical characteristics. The eN49P9 variant had less than a 2 fold change in neutralization potency in the 119 multi-clade compared to N49P9.6-FR-LS.

Overall, our findings suggest that a near-pan neutralizing bnAb against the CD4bs can be engineered for both enhanced antiviral activities and manufacturability. We have successfully engineered an HIV antibody for both improved potency and manufacturability. New antibodies such as eN49P9 may be considered for future clinical testing in combination with enhanced bnAbs of other specificity classes.

## Acknowledgments

This work was supported by Bill and Melinda Gates Foundation # INV-005284 (M.M.S) and # INV-036842 (M.S.S. and K.W.); NIH grants R01AI147870 (M.M.S), 1R01AI155150-01A1 (M.M.S/A.L.D.), R01 AI174908 (M.P.), P01 AI162242 (M.P.); the Intramural Research Program of the Division of Intramural Research, NIAID (Q.L. and P.L.); and VA Merit Award IBX004525 (M.M.S.). The CryoEM center at IBBR is supported by the University of Maryland Strategic Partnership (MPOWER) and by the NIST cooperative agreement 70NANB21H105. Use of the Stanford Synchrotron Radiation Lightsource, SLAC National Accelerator Laboratory, is supported by the U.S. Department of Energy, Office of Science, Office of Basic Energy Sciences under Contract No. DE-AC02-76SF00515. The SSRL Structural Molecular Biology Program is supported by the DOE Office of Biological and Environmental Research, and by the National Institutes of Health, National Institute of General Medical Sciences.

## Disclaimer

The views expressed in this manuscript are those of the authors and do not reflect the official policy or position of the Uniformed Services University, US Army, the Department of Defense, or the US Government.

**Figure S1.**
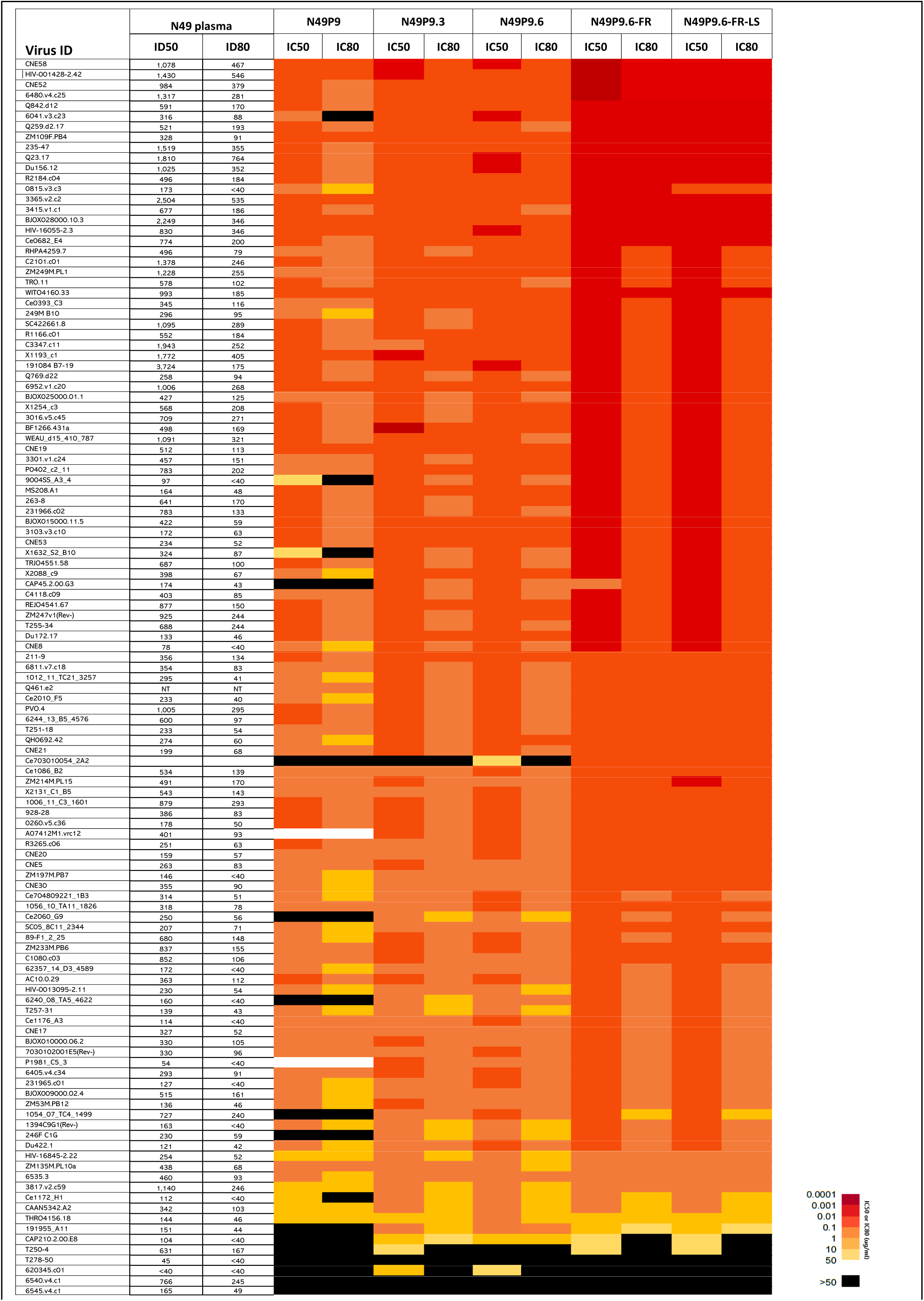
Neutralization testing of N49P9 family and variants in 118 multi-tier multiclade HIV pseudovirus assay. IC50 and IC80 values given as colors and heat map (refer to color key).

**Figure S2.**
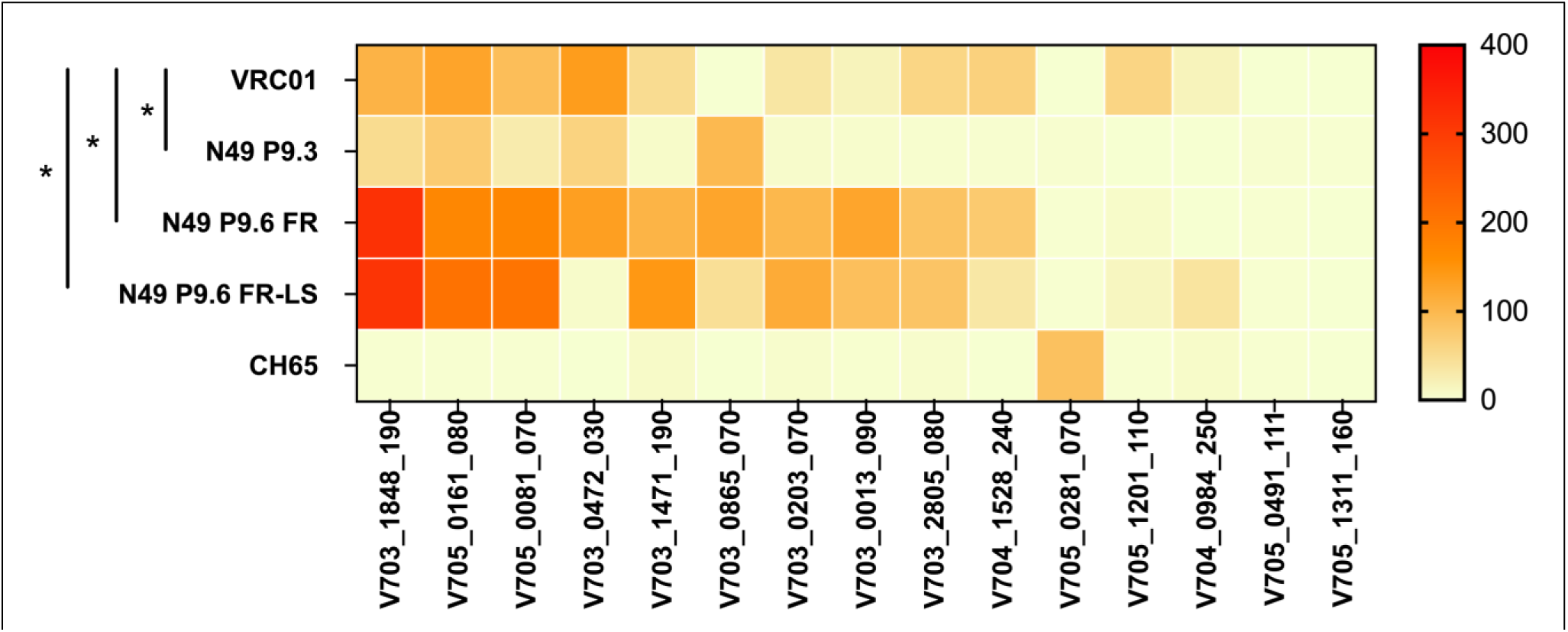
ADCC activity of N49 P series variants. The heat map represents the magnitude of responses against infectious molecular clones expressing recent Envelopes from the placebo group of phase II/III trials (HVTN703/704/705) as area under the curve (AUC) from 0 (no activity) to ≥400 (highest) as indicated by the scale on the right y-axis. Antibody-specific killing of each mAb listed on the left y-axis was conducted against each HIV IMC (x-axis) starting at 50μg/mL using a 5-fold dilution. Area-under-the-curves (AUCs) were then calculated using the trapezoid rule after subtracting the background activity. The anti-flu CH65 mAb was used as negative control. The parental antibody mAb N49P9.3 displayed significantly less ADCC activity than VRC01, while the engineered variants N49P9.6-FR, and N49P9.6-FR-LS demonstrated significantly more ADCC activity than VRC01, when tested by paired t test. * = P < .05 = P < .05

**Figure S3.**
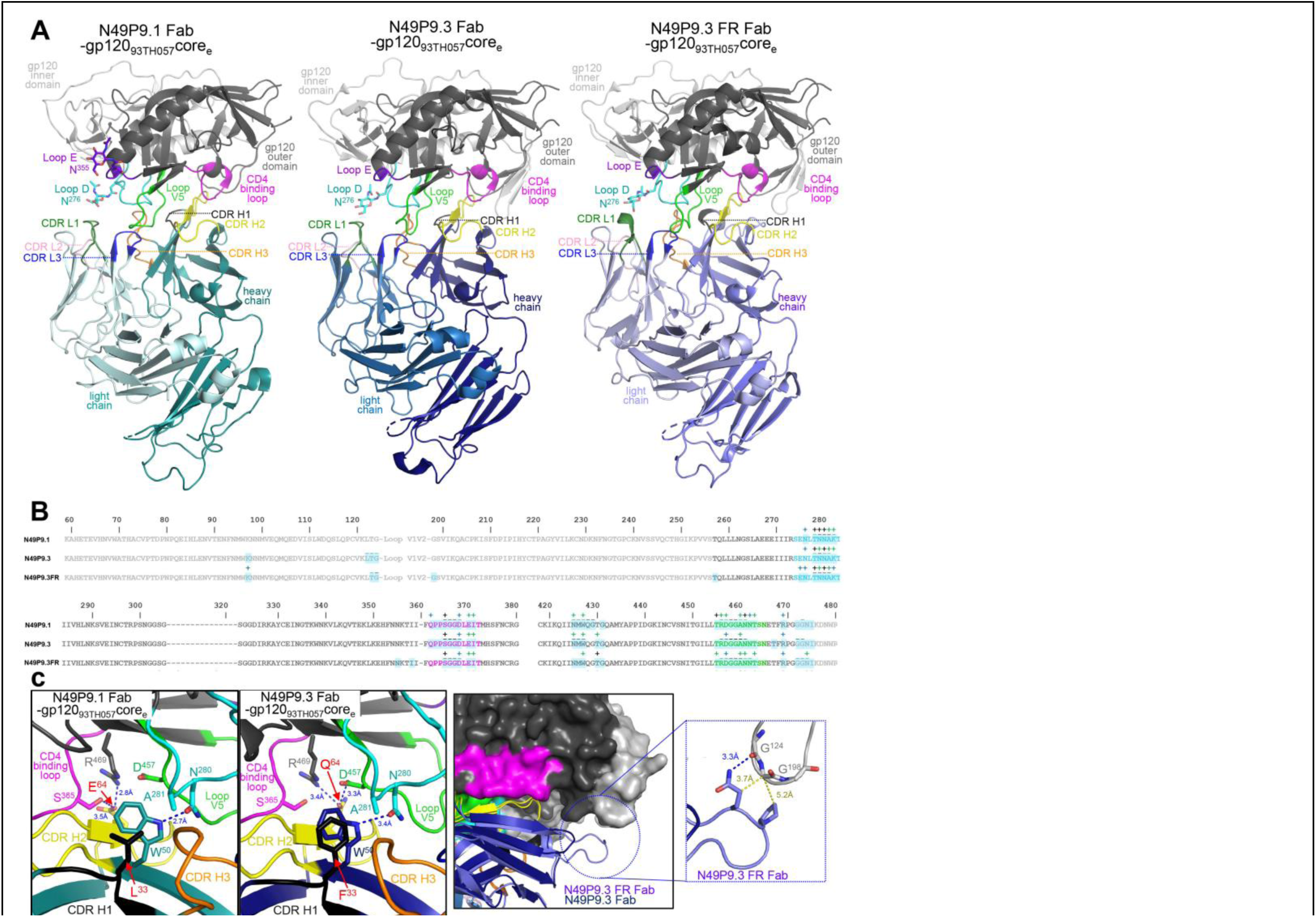
**Crystal structures of N49P9.1Fab-, N49P9.3-Fab– and N49P9.3FR Fab-gp120**_93TH057_**core**_e_ **complexes** (**a**) The overall structure of the complexes are shown as ribbon diagrams. The complementary-determining regions (CDRs) of Fabs are colored: CDR L1, green; CDR L2 light pink; CDR L3, blue; CDR H1, black; CDR H2, yellow and CDR H3, light orange. Outer and inner domains of gp120 are dark and light grey, respectively. The outer domain loops: D, E, CD4 binding, and V5 are colored in cyan, purple blue, magenta, and light green, respectively. Carbohydrates at position N^276^ (loop D) and N^355^ (loop E) are shown as sticks. (**b**) Epitope footprints of Fabs N49P9.1 and N49P9.3 mapped onto the gp120 primary sequences. Contact residues are defined by a 5 Å cutoff and marked above the sequence with (+) for side chain and (-) for main chain to indicate the type of contact: hydrophilic (blue), hydrophobic (green) and both (black). Buried surface residues determined by PISA are shaded blue for primary (**c**) Details of N49P9.3 Fab-gp120_93TH057_core_e_ interface with a blow-up view into the Fab contacts mediated by CDR H1 and 2 of N49P9.1 and N49P9.3 with colors indicated as in (a) and hydrogen bonds shown as dotted blue lines *(left panel)*. Interaction network of the frame region of N49P9.3-FR to gp120_93TH057_core_e_ *(right panel)*. H-bonds and Van der Waals contacts are shown as blue and yellow dotted lines, respectively. Residues that differ between N49P9.1 and N49P9.3 are labeled in red. Structures of N49P9.3 with and without the VRC03 FR insertion allowed us to assess the framework’s role in binding monomer gp120. Analysis of the N49P9.1 and N49P9.3 structures reveal that only two gp120 contact residues differ between the two, one in CDRH1 and one in CDRH2 (b and c). The first, Phe33 in CDRH1, packs against gp120 Ala281 in place of Leu33 in N49P9.1, but with slightly better van der Waals contacts, and modulates heavy chain Trp50, which also packs against gp120 Ala281 and forms a hydrogen bond to gp120 Asn280. The second Gln64 in CDRH2 can form hydrogen bonds with both gp120 Arg469 and Asp457 versus Glu64 in N49P9.1 which can only form a hydrogen bond with Arg469. Gln at position 64 is also what is seen in both N49P7 and VRC01 implying that Gln can make an important contribution to binding when placed here [22, 54]. These two amino acid changes likely account for much of N49P9.3’s increase in neutralization potency and breadth relative to N49P9.1. With the addition of the N49P9.3-FR structure We were able to confirm that the insertion in N49P9.3-FR makes little direct contact to gp120 core in the N49P9.3-FR Fab-gp12093TH057coree complex as expected and that the primary gp120 contacts remain intact; the VRC03 framework insertion is only expected to contact the adjacent gp120 protomer in the Env trimer [44]. The only direct contact the insert makes are to two glycines in the truncated V1V2 loop in the gp12093TH057coree construct which are not present in the original V1V2 loop sequence (Figure S3c).

**Figure S4.**
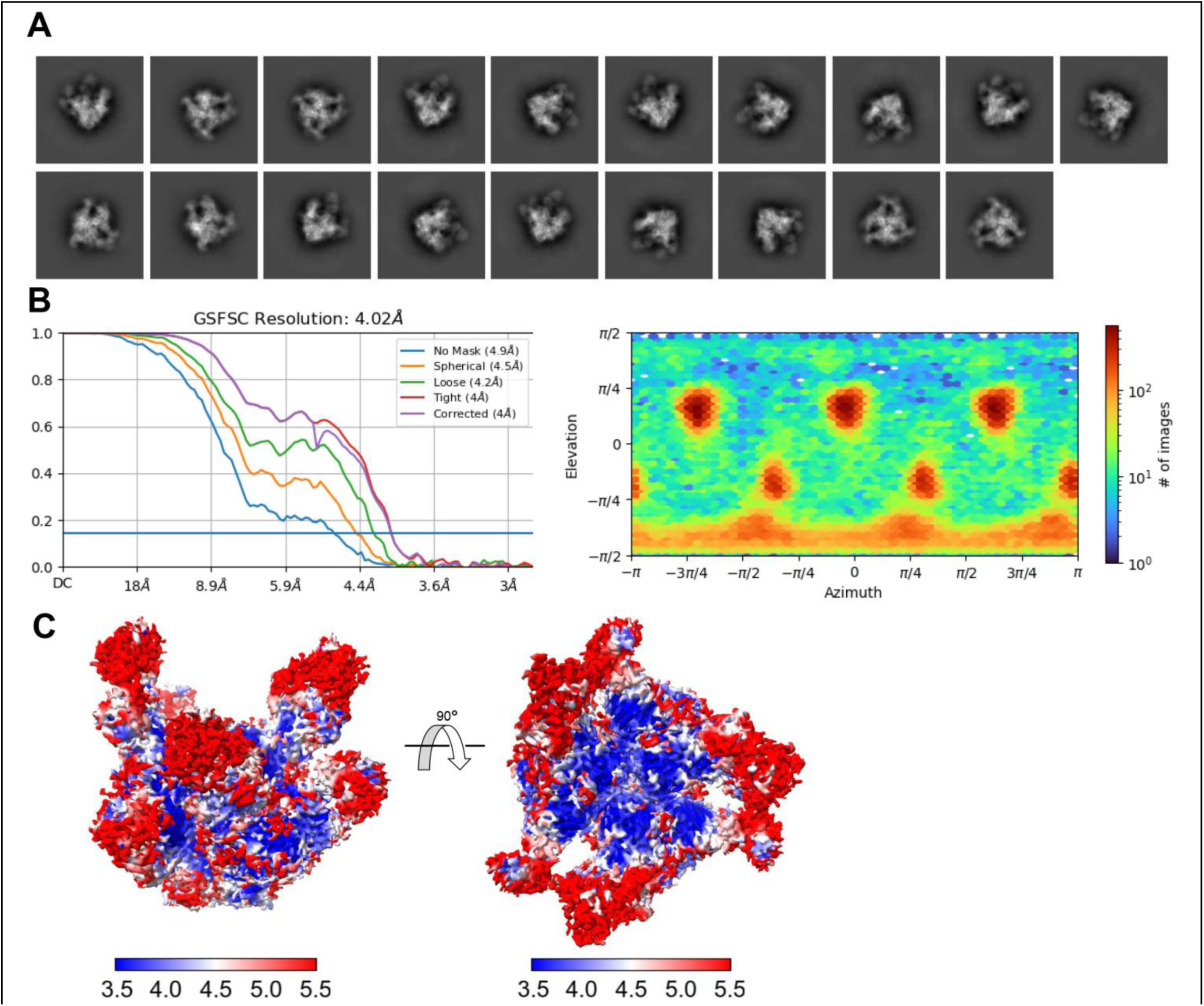
Cryo-EM structure of BG505 SOSIP.664 HIV-1 bound complex N49P9.6-FR and PGT121. (**a**) Selected 2D classes for *ab initio* map reconstruction(**b**) The Fourier shell correlation curves with spherical mask indicate the overall resolution (FSC cutoff 0.143) as determined by CryoSPARC and the direction distribution plot of all particles used in the final refinement. (**c**) Local resolution estimation plots with top and side views of the complex.

**Figure S5.**
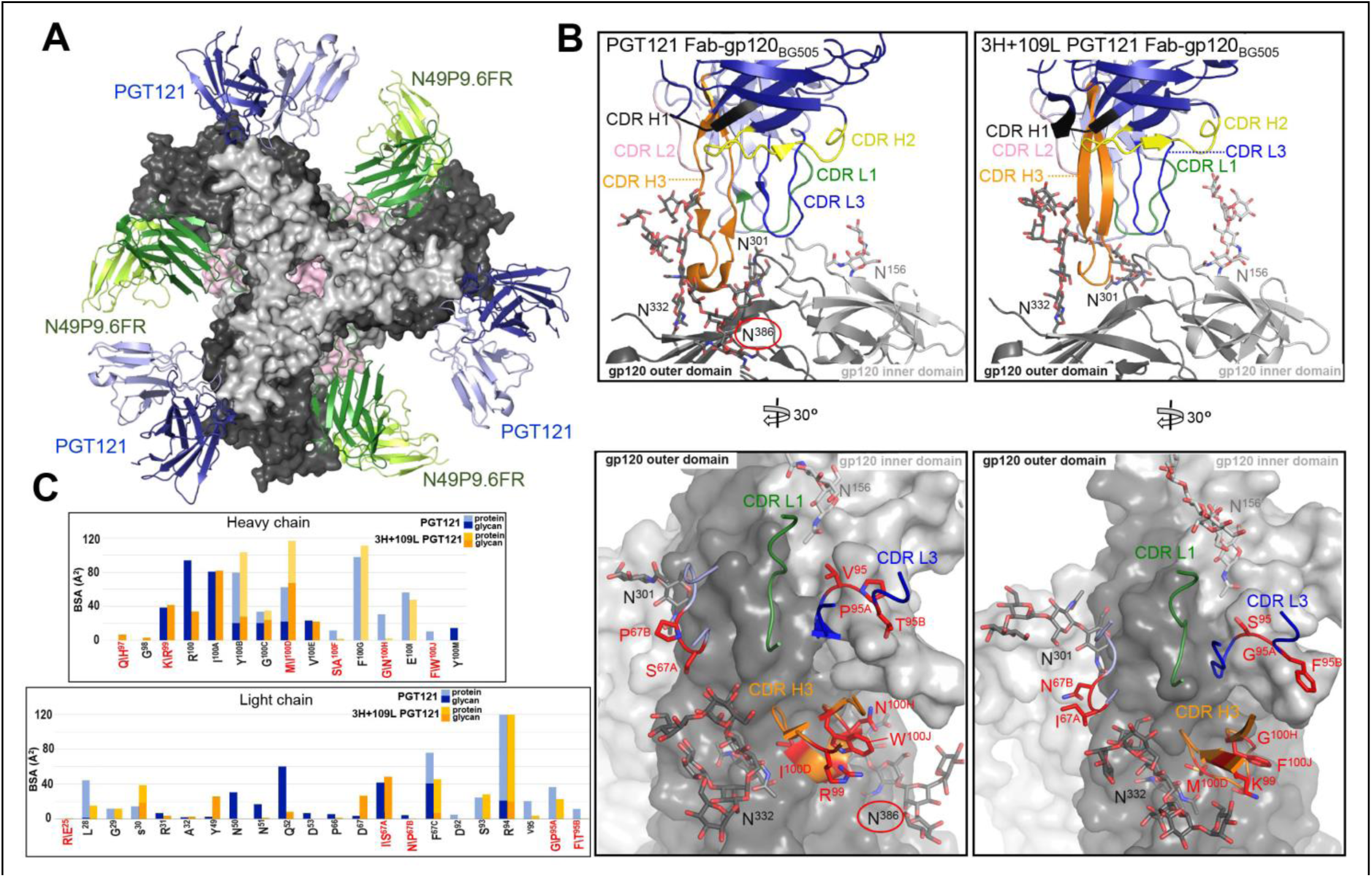
Molecular details of PGT121 bound to BG505 SOSIP.664. (**a**) Top view of the complex of BG505 SOSIP.664 HIV-1 Env trimer with N49P9.6-FR and PGT121 bound. N49P9.6-FR Fabs and gp120 are colored as in Figure 3. PGT121 (only the variable regions of the Fabs were built into the Cryo-EM model) are shown in blue and light blue ribbons. (**b**) Side-by-side comparison of PGT121 and the inferred germline PGT121 precursor, 3H+109L PGT121 (PDB ID: 5CEZ). CDRs are colored as in Figure 2. (**c**) BSA plot of Fab residues contributing to binding for PGT121 and 3H+109L PGT121. Protein or glycan portions are as indicated with residues that differ in sequence in red, 3H+109L PGT121 residue bottom and PGT121 residue top. One potential additional glycan interaction between PGT121’s light chain and the glycan attached to N^386^ that is absent in 3H+109L PGT121 is encircled in red in PGT121. The gp120 contact residues are largely identical between PGT121 and 3H+109L PGT121 but interactions PGT121 are more protein dependent. The total BSA for 3H+109L PGT121 is 2275 Å^2^ (1211 Å^2^ for gp120 and 1064 Å^2^ for Fab); total BSA for PGT121 is 2455 Å^2^ (1236 Å^2^ for gp120 and 1219 Å^2^ for Fab). Interactions between protein residues explain the total BSA of 1126 Å^2^ for 3H+109L PGT121 and the total BSA of 1308 Å^2^ for PGT121. Contributions from glycan are roughly comparible for both, 1126 Å^2^ for 3H+109L PGT121 and 1147 Å^2^ for PGT121. Thus one way PGT121 seems to have increased its affinity to Env during maturation is to have maximized its interactions with protein residues while maintaining similar interactions with key glycans.

**Figure S6.**
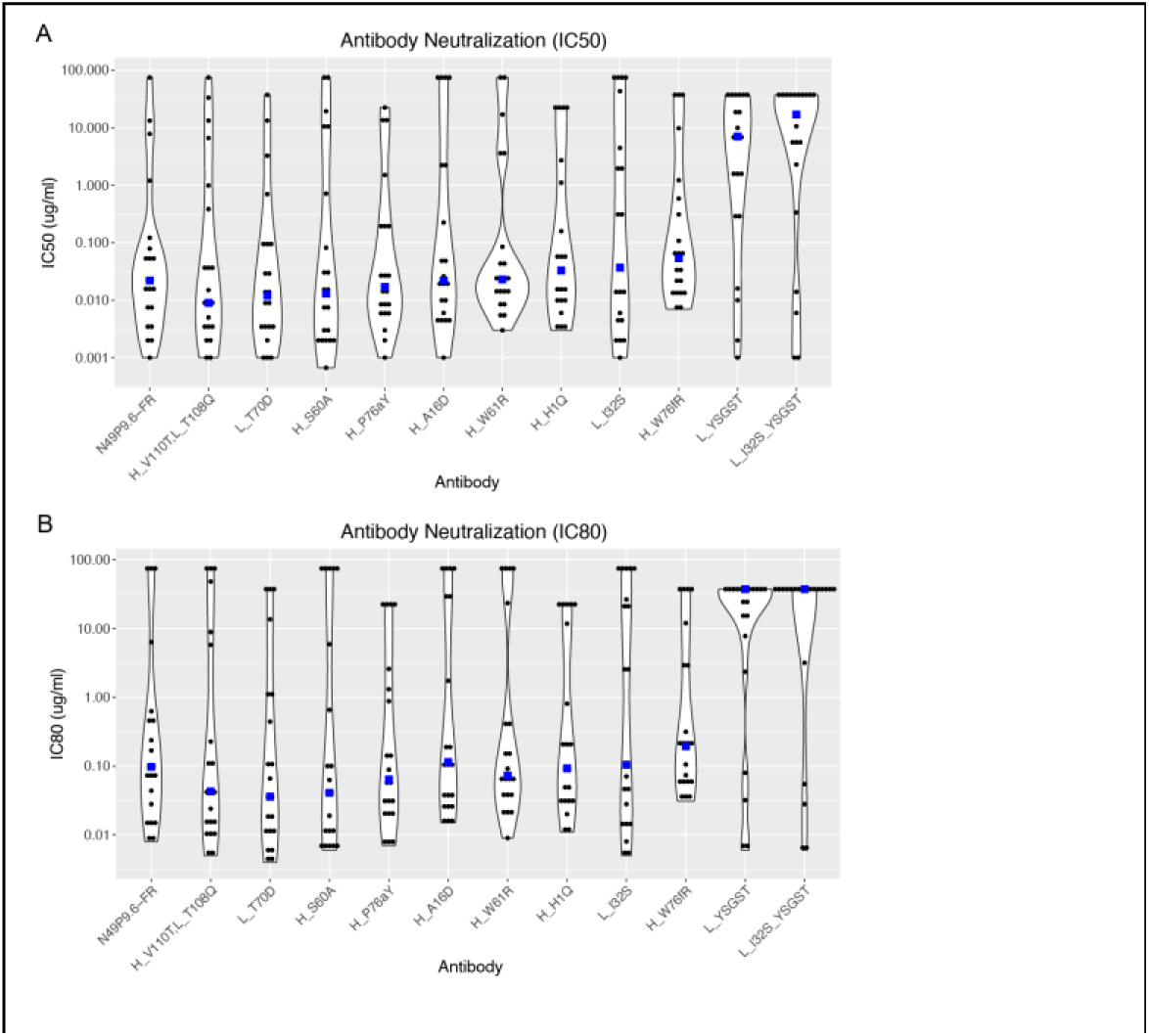
Neutralization assessment of N49P9.6-FR designs. Designed antibodies and N49P9.6-FR wild-type were tested for neutralization using a 21 virus global HIV panel, and measured (A) IC50 and (B) IC80 levels are shown as violin plots. Individual neutralization values are shown as black points, with median values shown as blue squares, and designs are ordered from lower to higher median IC50 levels (with wild-type on left for reference). In cases with unquantified neutralization, neutralization values are represented as 1.5-fold versus the unquantified level (e.g. 75 μg/ml for >50 μg/ml). Figure generated with ggplot2 [70].

**Figure S7.**
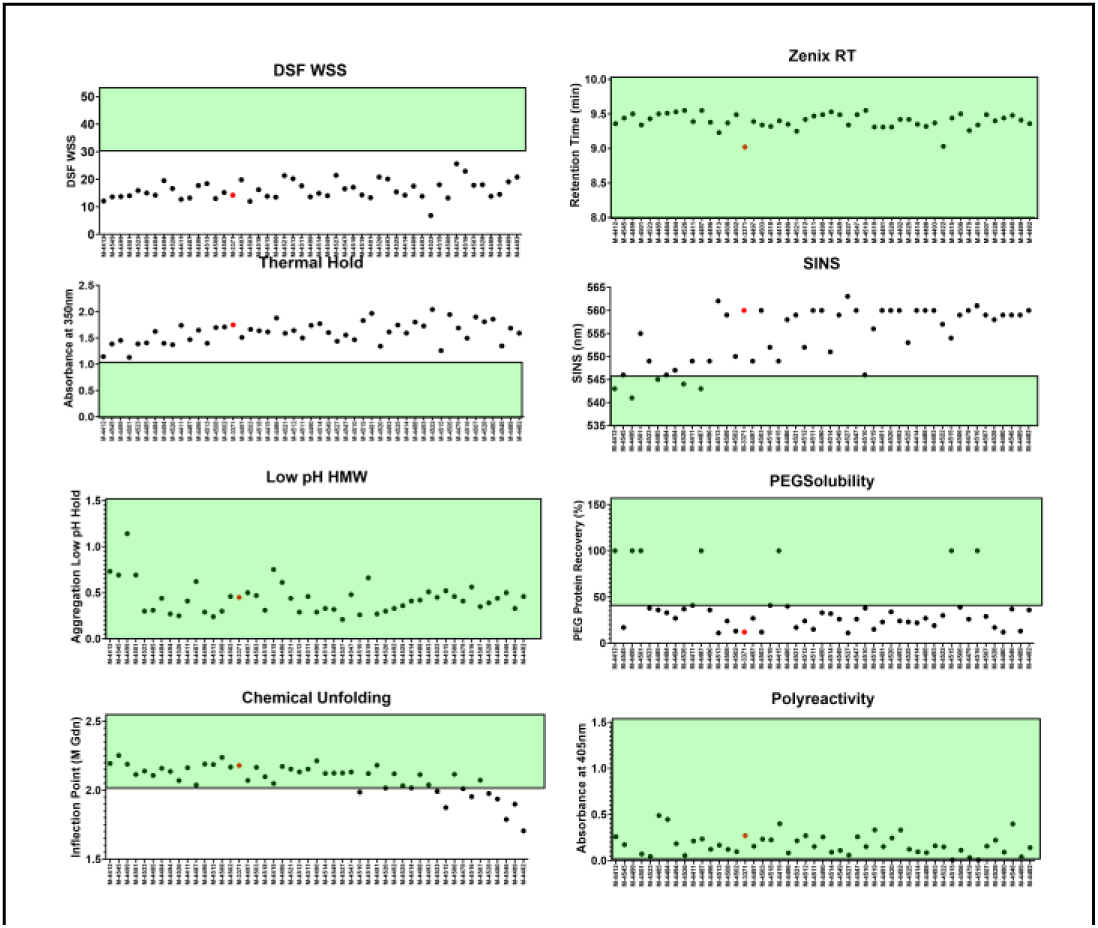
Conformational and colloidal stability results for variants selected from library panning. Regions of generally acceptable developability results are highlighted in green, N49P9.6-FR-LS are indicated in red circles compared with variants in black circles. Significant change in conformational stability was not observed while an increase colloidal stability was observed for multiple variants.

**Figure S8.**
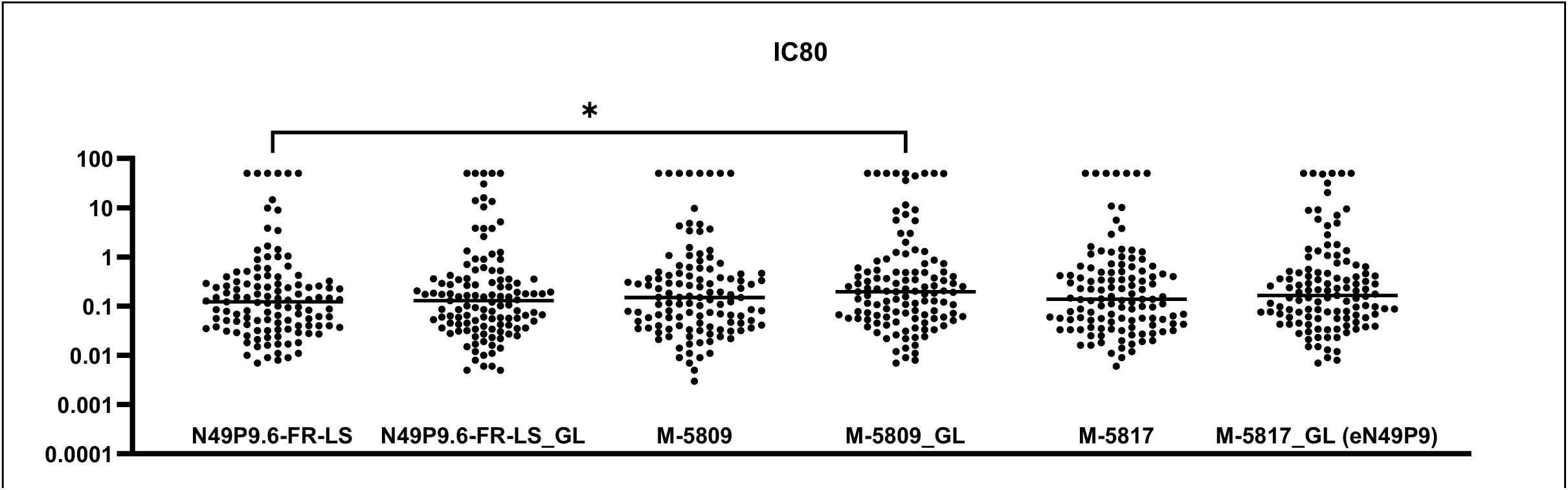
Neutralizing activity of N49P9.6-FR-LS variants selected for manufacturability characteristics. Six selected variants (see narrative) were expressed from stable cells pools and tested for neutralization against a 119 multi-tier multi-clade neutralization panel. All the variants tested had less than two fold difference in median IC80 compared to the parental N49P9.6-FR-LS, although this was significant only for M-5809_GL compared to N490P9.6-FR-LS (P=.04 by Mann-Whitney test). Variant M-5817_GL was the final variant chosen for clinical development (eN49P9). “-GL”: use of light chain with uncleaved N-terminus.

**Figure S9.**
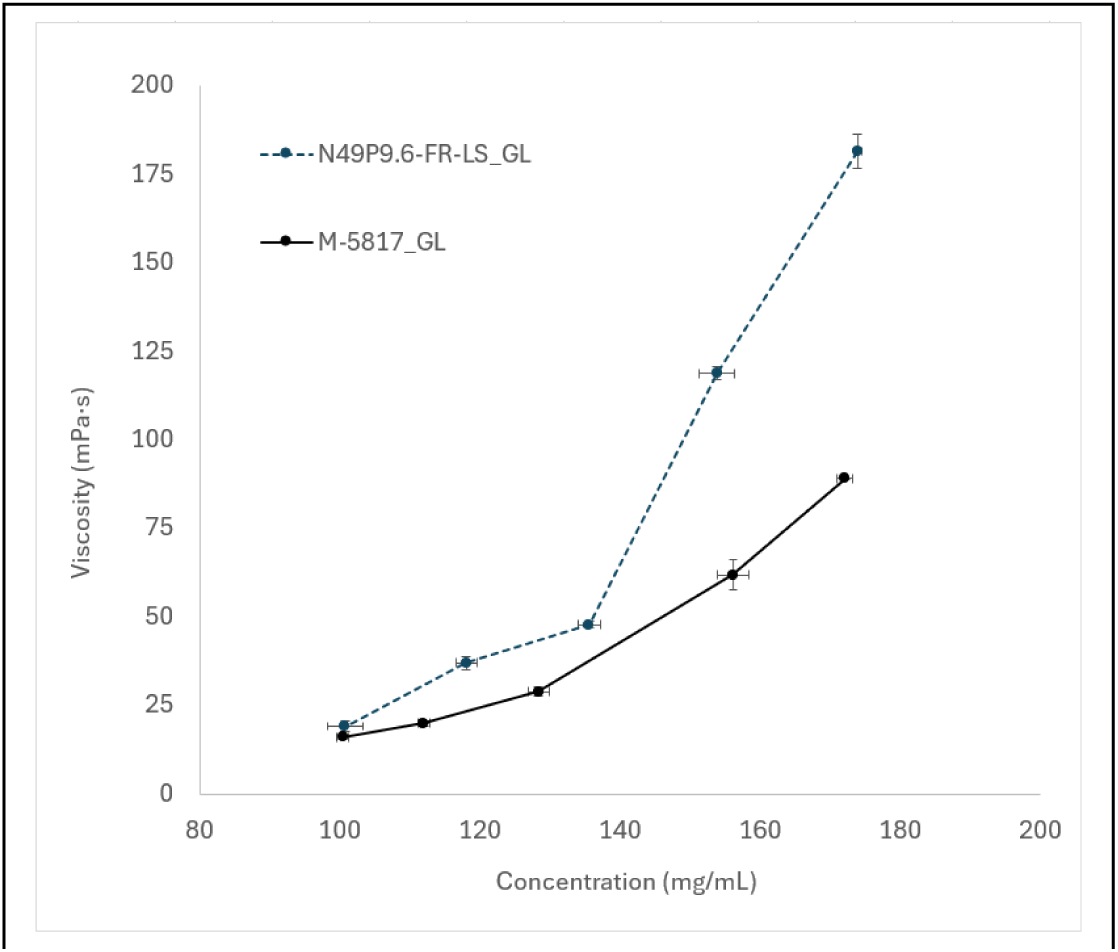
**Viscosity measurement of final variant and parental form**. Increased colloidal stability as measured by biophysical analysis was confirmed by viscosity measurement across increasing protein concentration. Decrease in viscosity was observed in the lead candidate, M-5817_GL (black line) as compared with the parental molecule N49P9.6-FR-LS (germline N-term) (blue dotted line). Standard deviation of triplicate protein concentration and viscosity measurements are capture in hash marks for each axis.

**Table S1.**
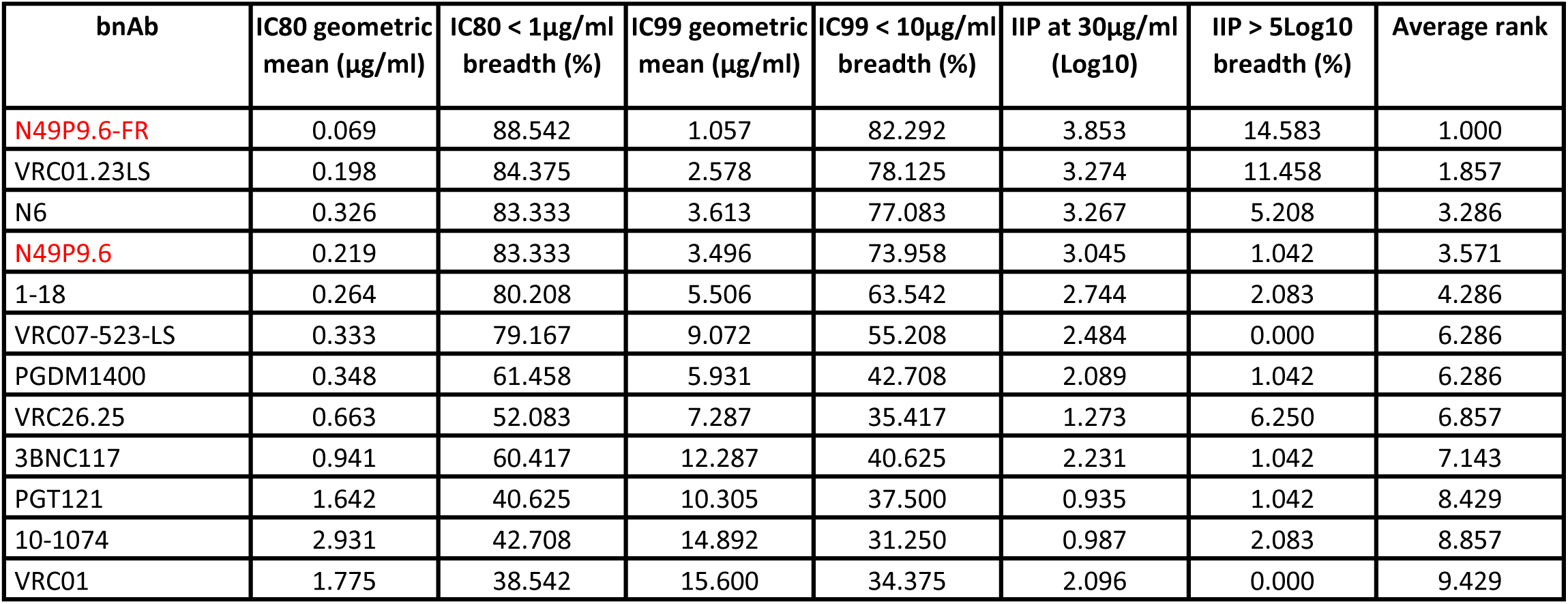
Single bnAb neutralization characteristics and ranking.

**Table S2.**
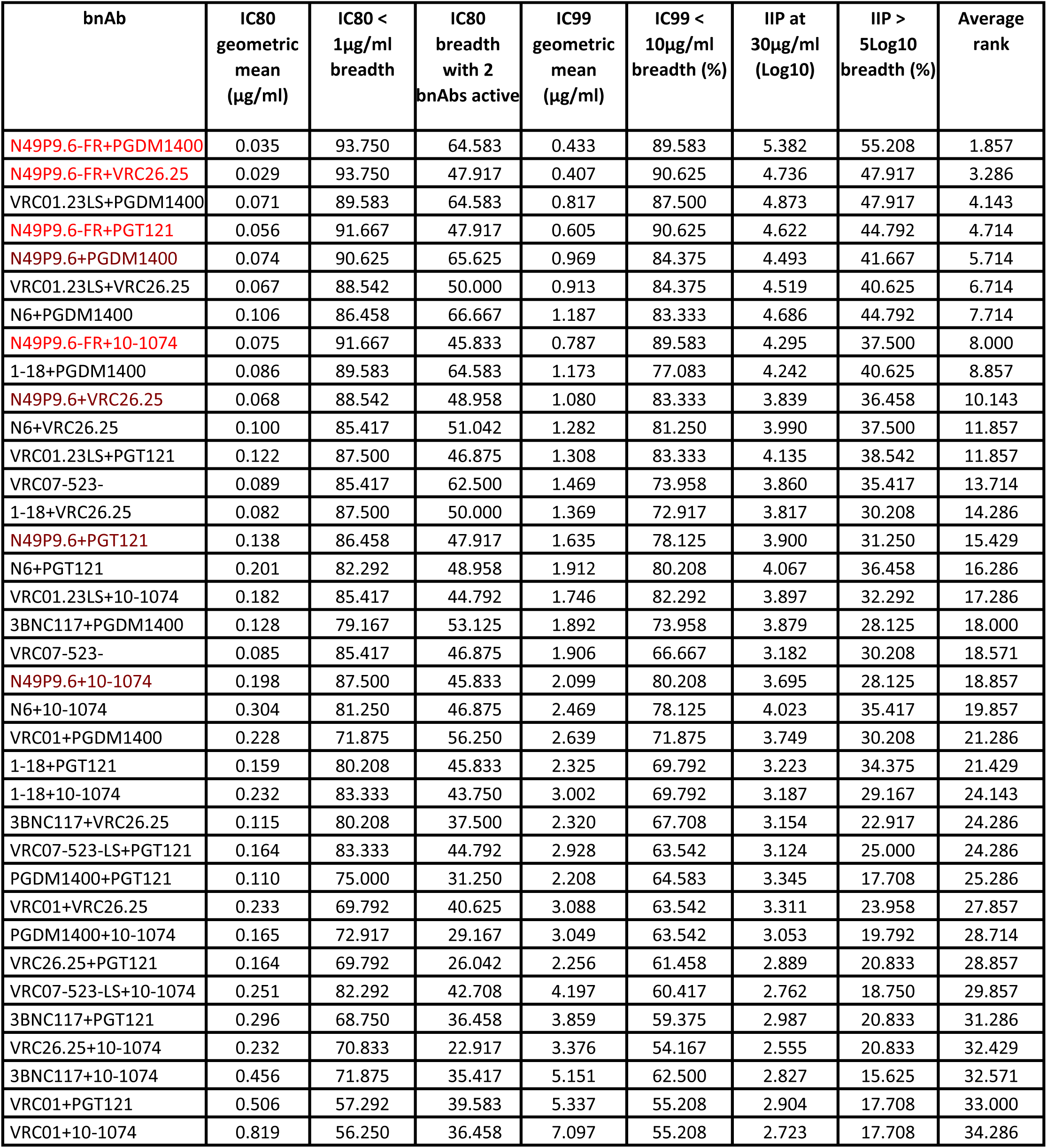
Dual bnAb neutralization characteristics and ranking.

**Table S3.**
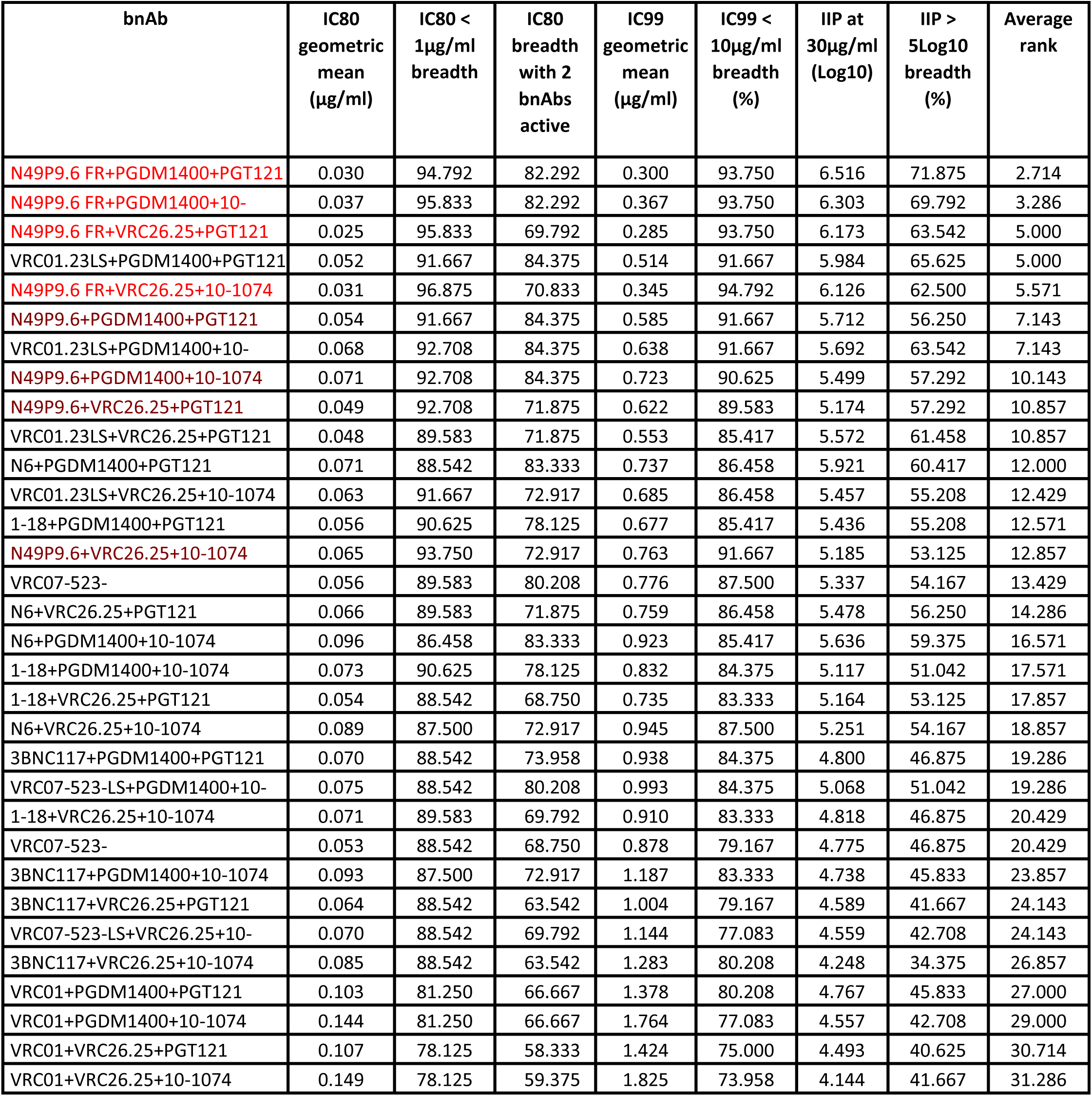
Triple bnAb neutralization characteristics and ranking.

**Table S4.**
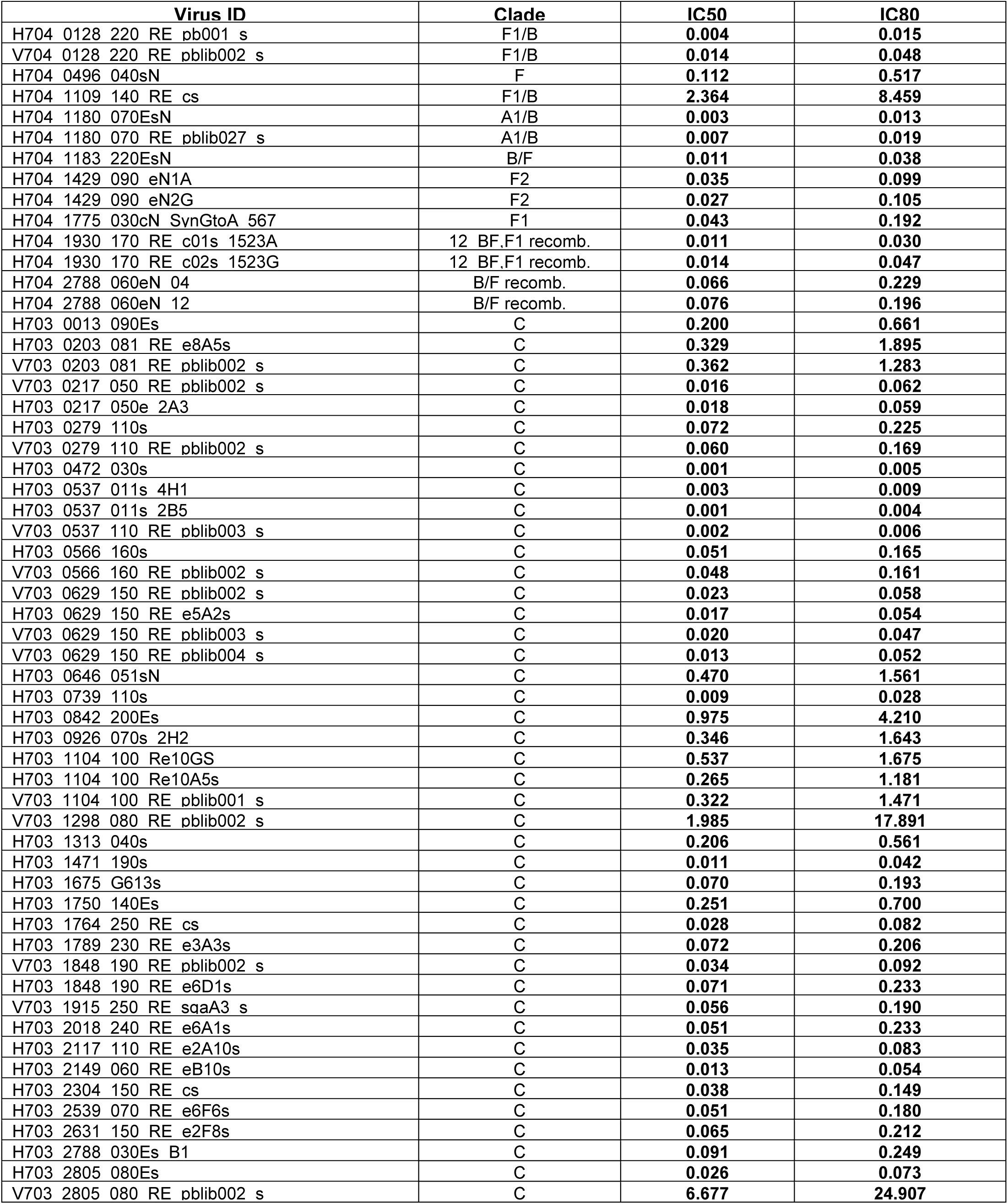
N49P9.6-FR-LS neutralization against AMP panel pseudoviruses in TZM.bl cells.

**Table S5.**
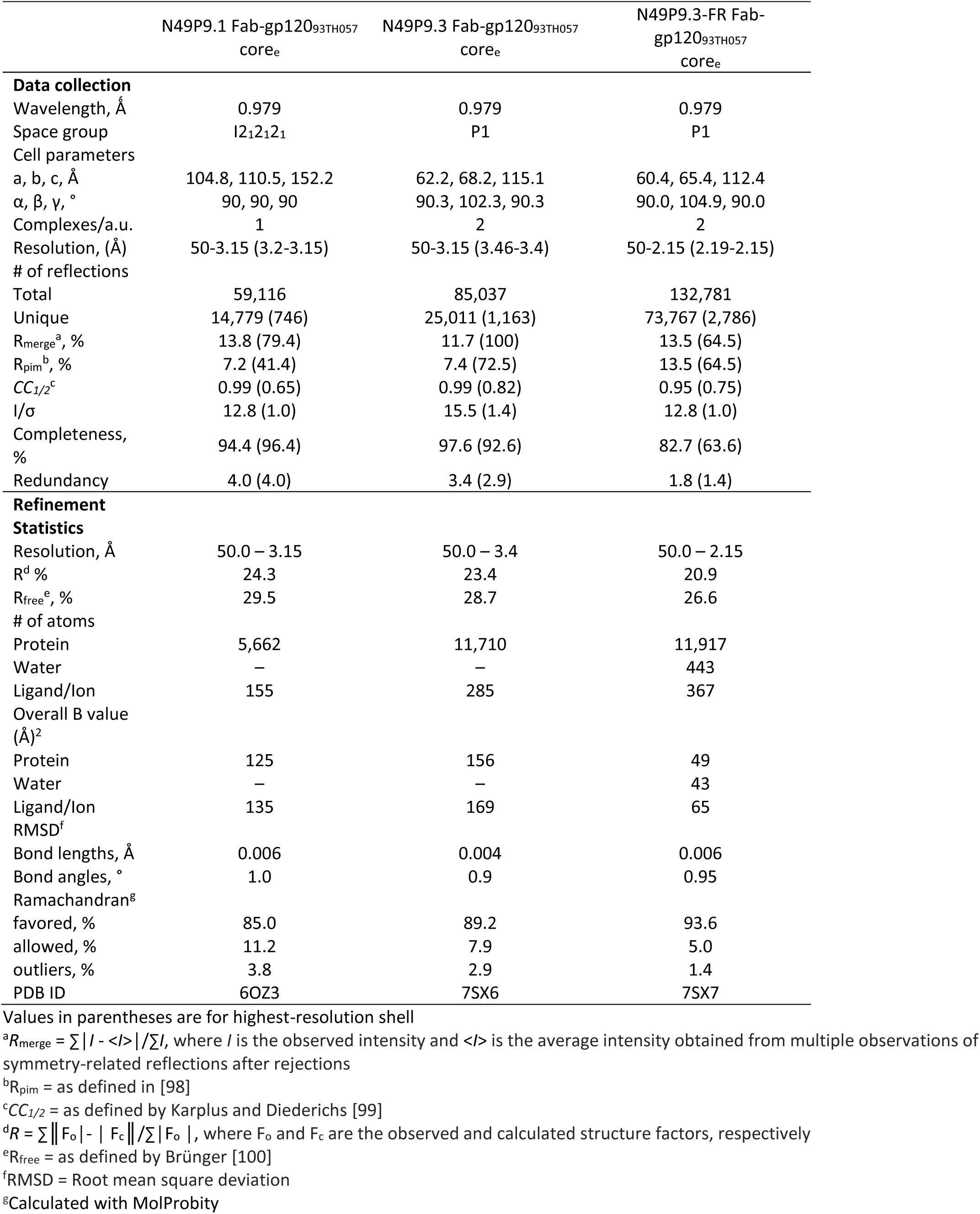
Crystallographic Data collection and refinement statistics.

**Table S6.**
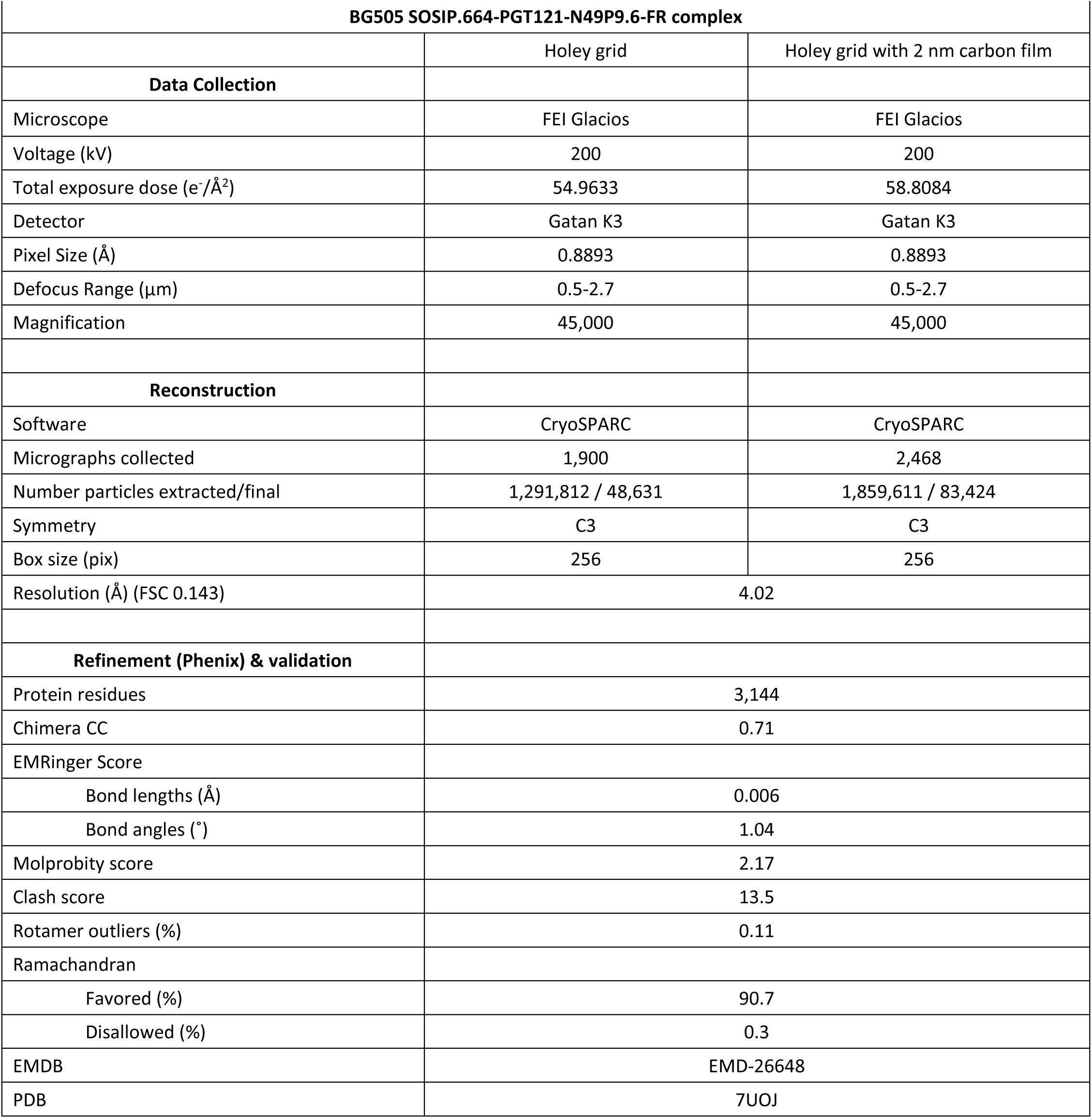
CryoEM Data collection and refinement statistics.

**Table S7.**
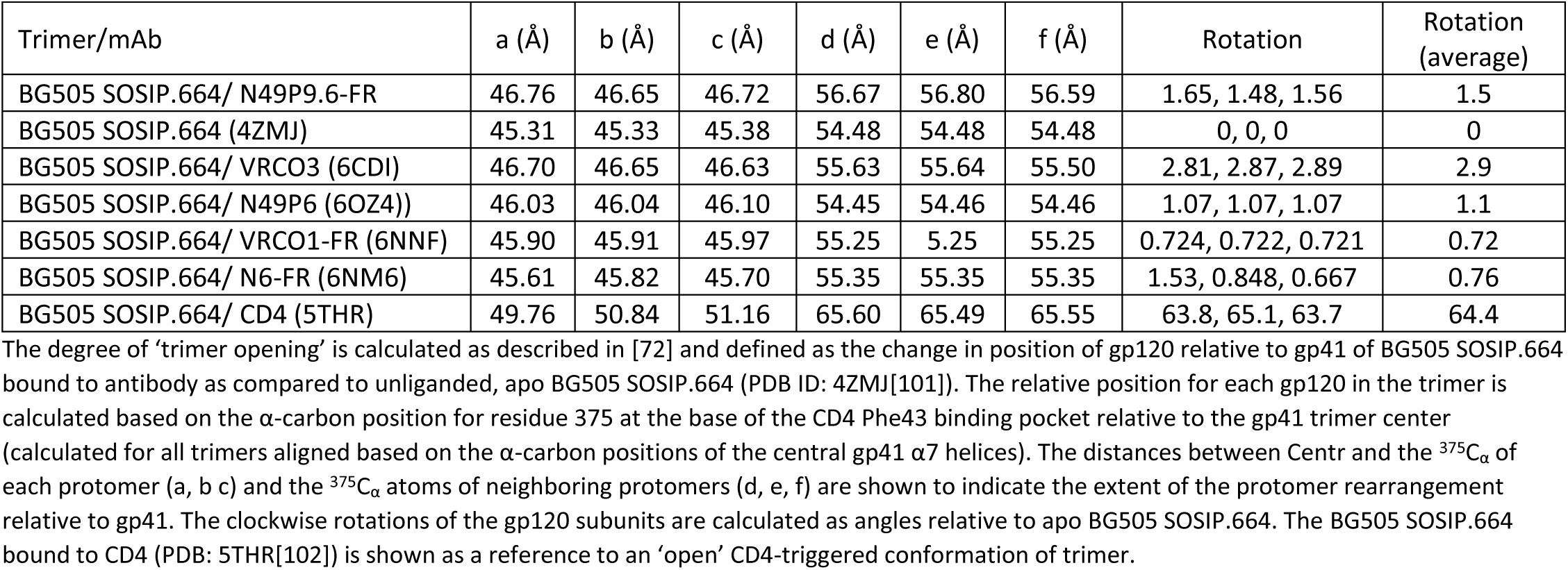
Changes to the BG505 SOSIP.664 trimer upon binding to CD4bs mAbs that contact the adjacent gp120 protomer.

**Table S8.**
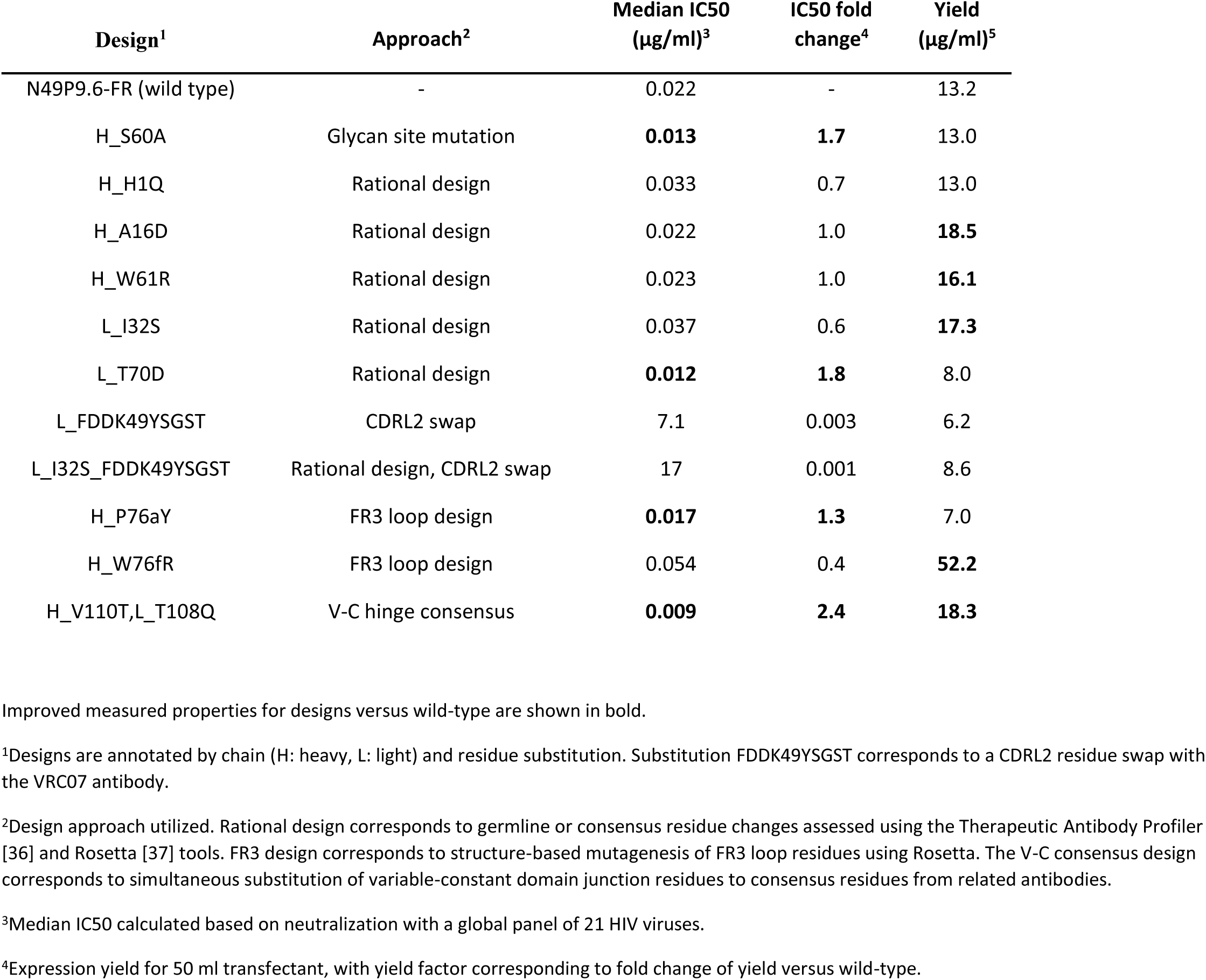
Viral neutralization and expression yield for N49P9.6-FR3 antibody designs.

**Table S9.**
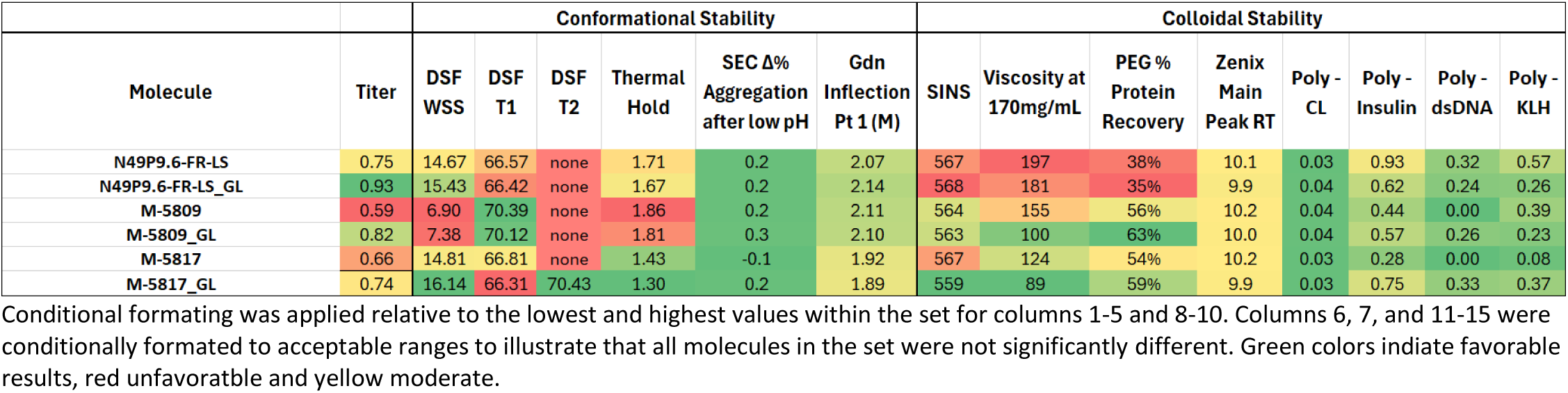
Final variant set showing biophsyical characterization analysis.

| Molecule | Titer | Conformational Stability |  |  |  |  |  | Colloidal Stability |  |  |  |  |  |  |  |
| --- | --- | --- | --- | --- | --- | --- | --- | --- | --- | --- | --- | --- | --- | --- | --- |
|  |  | DSF WSS | DSF T1 | DSF T2 | Thermal Hold | SEC Δ% Aggregation after low pH | Gdn Inflection Pt 1 (M) | SINS | Viscosity at 170mg/mL | PEG % Protein Recovery | Zenix Main Peak RT | Poly - CL | Poly - Insulin | Poly - dsDNA | Poly - KLH |
| N49P9.6-FR-LS | 0.75 | 14.67 | 66.57 | none | 1.71 | 0.2 | 2.07 | 567 | 197 | 38% | 10.1 | 0.03 | 0.93 | 0.32 | 0.57 |
| N49P9.6-FR-LS_GL | 0.93 | 15.43 | 66.42 | none | 1.67 | 0.2 | 2.14 | 568 | 181 | 35% | 9.9 | 0.04 | 0.62 | 0.24 | 0.26 |
| M-5809 | 0.59 | 6.90 | 70.39 | none | 1.86 | 0.2 | 2.11 | 564 | 155 | 56% | 10.2 | 0.04 | 0.44 | 0.00 | 0.39 |
| M-5809_GL | 0.82 | 7.38 | 70.12 | none | 1.81 | 0.3 | 2.10 | 563 | 100 | 63% | 10.0 | 0.04 | 0.57 | 0.26 | 0.23 |
| M-5817 | 0.66 | 14.81 | 66.81 | none | 1.43 | -0.1 | 1.92 | 567 | 124 | 54% | 10.2 | 0.03 | 0.28 | 0.00 | 0.08 |
| M-5817_GL | 0.74 | 16.14 | 66.31 | 70.43 | 1.30 | 0.2 | 1.89 | 559 | 89 | 59% | 9.9 | 0.03 | 0.75 | 0.33 | 0.37 |

Conditional formatting was applied relative to the lowest and highest values within the set for columns 1-5 and 8-10. Columns 6, 7, and 11-15 were conditionally formatted to acceptable ranges to illustrate that all molecules in the set were not significantly different. Green colors indicate favorable results, red unfavorable and yellow moderate.

## Notes

### Competing Interest Statement

The authors have declared no competing interest.

