## Supplementary material for "A comprehensive engineering strategy improves potency and manufacturability of a near pan-neutralizing antibody against HIV": Figure S1

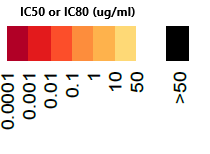

|  | **N49 plasma** | | **N49P9** | | **N49P9.3** | | **N49P9.6** | | **N49P9.6-FR** | | **N49P9.6-FR-LS** | |
| --- | --- | --- | --- | --- | --- | --- | --- | --- | --- | --- | --- | --- |
| **Virus ID** | **ID50** | **ID80** | **IC50** | **IC80** | **IC50** | **IC80** | **IC50** | **IC80** | **IC50** | **IC80** | **IC50** | **IC80** |
| CNE58 | 1,078 | 467 |  |  |  |  |  |  |  |  |  |  |
| HIV-001428-2.42 | 1,430 | 546 |  |  |  |  |  |  |  |  |  |  |
| CNE52 | 984 | 379 |  |  |  |  |  |  |  |  |  |  |
| 6480.v4.c25 | 1,317 | 281 |  |  |  |  |  |  |  |  |  |  |
| Q842.d12 | 591 | 170 |  |  |  |  |  |  |  |  |  |  |
| 6041.v3.c23 | 316 | 88 |  |  |  |  |  |  |  |  |  |  |
| Q259.d2.17 | 521 | 193 |  |  |  |  |  |  |  |  |  |  |
| ZM109F.PB4 | 328 | 91 |  |  |  |  |  |  |  |  |  |  |
| 235-47 | 1,519 | 355 |  |  |  |  |  |  |  |  |  |  |
| Q23.17 | 1,810 | 764 |  |  |  |  |  |  |  |  |  |  |
| Du156.12 | 1,025 | 352 |  |  |  |  |  |  |  |  |  |  |
| R2184.c04 | 496 | 184 |  |  |  |  |  |  |  |  |  |  |
| 0815.v3.c3 | 173 | <40 |  |  |  |  |  |  |  |  |  |  |
| 3365.v2.c2 | 2,504 | 535 |  |  |  |  |  |  |  |  |  |  |
| 3415.v1.c1 | 677 | 186 |  |  |  |  |  |  |  |  |  |  |
| BJOX028000.10.3 | 2,249 | 346 |  |  |  |  |  |  |  |  |  |  |
| HIV-16055-2.3 | 830 | 346 |  |  |  |  |  |  |  |  |  |  |
| Ce0682_E4 | 774 | 200 |  |  |  |  |  |  |  |  |  |  |
| RHPA4259.7 | 496 | 79 |  |  |  |  |  |  |  |  |  |  |
| C2101.c01 | 1,378 | 246 |  |  |  |  |  |  |  |  |  |  |
| ZM249M.PL1 | 1,228 | 255 |  |  |  |  |  |  |  |  |  |  |
| TRO.11 | 578 | 102 |  |  |  |  |  |  |  |  |  |  |
| WITO4160.33 | 993 | 185 |  |  |  |  |  |  |  |  |  |  |
| Ce0393_C3 | 345 | 116 |  |  |  |  |  |  |  |  |  |  |
| 249M B10 | 296 | 95 |  |  |  |  |  |  |  |  |  |  |
| SC422661.8 | 1,095 | 289 |  |  |  |  |  |  |  |  |  |  |
| R1166.c01 | 552 | 184 |  |  |  |  |  |  |  |  |  |  |
| C3347.c11 | 1,943 | 252 |  |  |  |  |  |  |  |  |  |  |
| X1193_c1 | 1,772 | 405 |  |  |  |  |  |  |  |  |  |  |
| 191084 B7-19 | 3,724 | 175 |  |  |  |  |  |  |  |  |  |  |
| Q769.d22 | 258 | 94 |  |  |  |  |  |  |  |  |  |  |
| 6952.v1.c20 | 1,006 | 268 |  |  |  |  |  |  |  |  |  |  |
| BJOX025000.01.1 | 427 | 125 |  |  |  |  |  |  |  |  |  |  |
| X1254_c3 | 568 | 208 |  |  |  |  |  |  |  |  |  |  |
| 3016.v5.c45 | 709 | 271 |  |  |  |  |  |  |  |  |  |  |
| BF1266.431a | 498 | 169 |  |  |  |  |  |  |  |  |  |  |
| WEAU_d15_410_787 | 1,091 | 321 |  |  |  |  |  |  |  |  |  |  |
| CNE19 | 512 | 113 |  |  |  |  |  |  |  |  |  |  |
| 3301.v1.c24 | 457 | 151 |  |  |  |  |  |  |  |  |  |  |
| P0402_c2_11 | 783 | 202 |  |  |  |  |  |  |  |  |  |  |
| 9004SS_A3_4 | 97 | <40 |  |  |  |  |  |  |  |  |  |  |
| MS208.A1 | 164 | 48 |  |  |  |  |  |  |  |  |  |  |
| 263-8 | 641 | 170 |  |  |  |  |  |  |  |  |  |  |
| 231966.c02 | 783 | 133 |  |  |  |  |  |  |  |  |  |  |
| BJOX015000.11.5 | 422 | 59 |  |  |  |  |  |  |  |  |  |  |
| 3103.v3.c10 | 172 | 63 |  |  |  |  |  |  |  |  |  |  |
| CNE53 | 234 | 52 |  |  |  |  |  |  |  |  |  |  |
| X1632_S2_B10 | 324 | 87 |  |  |  |  |  |  |  |  |  |  |
| TRJO4551.58 | 687 | 100 |  |  |  |  |  |  |  |  |  |  |
| X2088_c9 | 398 | 67 |  |  |  |  |  |  |  |  |  |  |
| CAP45.2.00.G3 | 174 | 43 |  |  |  |  |  |  |  |  |  |  |
| C4118.c09 | 403 | 85 |  |  |  |  |  |  |  |  |  |  |
| REJO4541.67 | 877 | 150 |  |  |  |  |  |  |  |  |  |  |
| ZM247v1(Rev-) | 925 | 244 |  |  |  |  |  |  |  |  |  |  |
| T255-34 | 688 | 244 |  |  |  |  |  |  |  |  |  |  |
| Du172.17 | 133 | 46 |  |  |  |  |  |  |  |  |  |  |
| CNE8 | 78 | <40 |  |  |  |  |  |  |  |  |  |  |
| 211-9 | 356 | 134 |  |  |  |  |  |  |  |  |  |  |
| 6811.v7.c18 | 354 | 83 |  |  |  |  |  |  |  |  |  |  |
| 1012_11_TC21_3257 | 295 | 41 |  |  |  |  |  |  |  |  |  |  |
| Q461.e2 | NT | NT |  |  |  |  |  |  |  |  |  |  |
| Ce2010_F5 | 233 | 40 |  |  |  |  |  |  |  |  |  |  |
| PVO.4 | 1,005 | 295 |  |  |  |  |  |  |  |  |  |  |
| 6244_13_B5_4576 | 600 | 97 |  |  |  |  |  |  |  |  |  |  |
| T251-18 | 233 | 54 |  |  |  |  |  |  |  |  |  |  |
| QH0692.42 | 274 | 60 |  |  |  |  |  |  |  |  |  |  |
| CNE21 | 199 | 68 |  |  |  |  |  |  |  |  |  |  |
| Ce703010054_2A2 |  |  |  |  |  |  |  |  |  |  |  |  |
| Ce1086_B2 | 534 | 139 |  |  |  |  |  |  |  |  |  |  |
| ZM214M.PL15 | 491 | 170 |  |  |  |  |  |  |  |  |  |  |
| X2131_C1_B5 | 543 | 143 |  |  |  |  |  |  |  |  |  |  |
| 1006_11_C3_1601 | 879 | 293 |  |  |  |  |  |  |  |  |  |  |
| 928-28 | 386 | 83 |  |  |  |  |  |  |  |  |  |  |
| 0260.v5.c36 | 178 | 50 |  |  |  |  |  |  |  |  |  |  |
| A07412M1.vrc12 | 401 | 93 |  |  |  |  |  |  |  |  |  |  |
| R3265.c06 | 251 | 63 |  |  |  |  |  |  |  |  |  |  |
| CNE20 | 159 | 57 |  |  |  |  |  |  |  |  |  |  |
| CNE5 | 263 | 83 |  |  |  |  |  |  |  |  |  |  |
| ZM197M.PB7 | 146 | <40 |  |  |  |  |  |  |  |  |  |  |
| CNE30 | 355 | 90 |  |  |  |  |  |  |  |  |  |  |
| Ce704809221_1B3 | 314 | 51 |  |  |  |  |  |  |  |  |  |  |
| 1056_10_TA11_1826 | 318 | 78 |  |  |  |  |  |  |  |  |  |  |
| Ce2060_G9 | 250 | 56 |  |  |  |  |  |  |  |  |  |  |
| SC05_8C11_2344 | 207 | 71 |  |  |  |  |  |  |  |  |  |  |
| 89-F1_2_25 | 680 | 148 |  |  |  |  |  |  |  |  |  |  |
| ZM233M.PB6 | 837 | 155 |  |  |  |  |  |  |  |  |  |  |
| C1080.c03 | 852 | 106 |  |  |  |  |  |  |  |  |  |  |
| 62357_14_D3_4589 | 172 | <40 |  |  |  |  |  |  |  |  |  |  |
| AC10.0.29 | 363 | 112 |  |  |  |  |  |  |  |  |  |  |
| HIV-0013095-2.11 | 230 | 54 |  |  |  |  |  |  |  |  |  |  |
| 6240_08_TA5_4622 | 160 | <40 |  |  |  |  |  |  |  |  |  |  |
| T257-31 | 139 | 43 |  |  |  |  |  |  |  |  |  |  |
| Ce1176_A3 | 114 | <40 |  |  |  |  |  |  |  |  |  |  |
| CNE17 | 327 | 52 |  |  |  |  |  |  |  |  |  |  |
| BJOX010000.06.2 | 330 | 105 |  |  |  |  |  |  |  |  |  |  |
| 7030102001E5(Rev-) | 330 | 96 |  |  |  |  |  |  |  |  |  |  |
| P1981_C5_3 | 54 | <40 |  |  |  |  |  |  |  |  |  |  |
| 6405.v4.c34 | 293 | 91 |  |  |  |  |  |  |  |  |  |  |
| 231965.c01 | 127 | <40 |  |  |  |  |  |  |  |  |  |  |
| BJOX009000.02.4 | 515 | 161 |  |  |  |  |  |  |  |  |  |  |
| ZM53M.PB12 | 136 | 46 |  |  |  |  |  |  |  |  |  |  |
| 1054_07_TC4_1499 | 727 | 240 |  |  |  |  |  |  |  |  |  |  |
| 1394C9G1(Rev-) | 163 | <40 |  |  |  |  |  |  |  |  |  |  |
| 246F C1G | 230 | 59 |  |  |  |  |  |  |  |  |  |  |
| Du422.1 | 121 | 42 |  |  |  |  |  |  |  |  |  |  |
| HIV-16845-2.22 | 254 | 52 |  |  |  |  |  |  |  |  |  |  |
| ZM135M.PL10a | 438 | 68 |  |  |  |  |  |  |  |  |  |  |
| 6535.3 | 460 | 93 |  |  |  |  |  |  |  |  |  |  |
| 3817.v2.c59 | 1,140 | 246 |  |  |  |  |  |  |  |  |  |  |
| Ce1172_H1 | 112 | <40 |  |  |  |  |  |  |  |  |  |  |
| CAAN5342.A2 | 342 | 103 |  |  |  |  |  |  |  |  |  |  |
| THRO4156.18 | 144 | 46 |  |  |  |  |  |  |  |  |  |  |
| 191955_A11 | 151 | 44 |  |  |  |  |  |  |  |  |  |  |
| CAP210.2.00.E8 | 104 | <40 |  |  |  |  |  |  |  |  |  |  |
| T250-4 | 631 | 167 |  |  |  |  |  |  |  |  |  |  |
| T278-50 | 45 | <40 |  |  |  |  |  |  |  |  |  |  |
| 620345.c01 | <40 | <40 |  |  |  |  |  |  |  |  |  |  |
| 6540.v4.c1 | 766 | 245 |  |  |  |  |  |  |  |  |  |  |
| 6545.v4.c1 | 165 | 49 |  |  |  |  |  |  |  |  |  |  |

**Figure S1. Neutralization testing of N49P9 family and variants in 118 multi-tier multiclade HIV pseudovirus assay.** IC50 and IC80 values given as colors and heat map (refer to color key).
