## Supplementary material for "A comprehensive engineering strategy improves potency and manufacturability of a near pan-neutralizing antibody against HIV": Figure S2

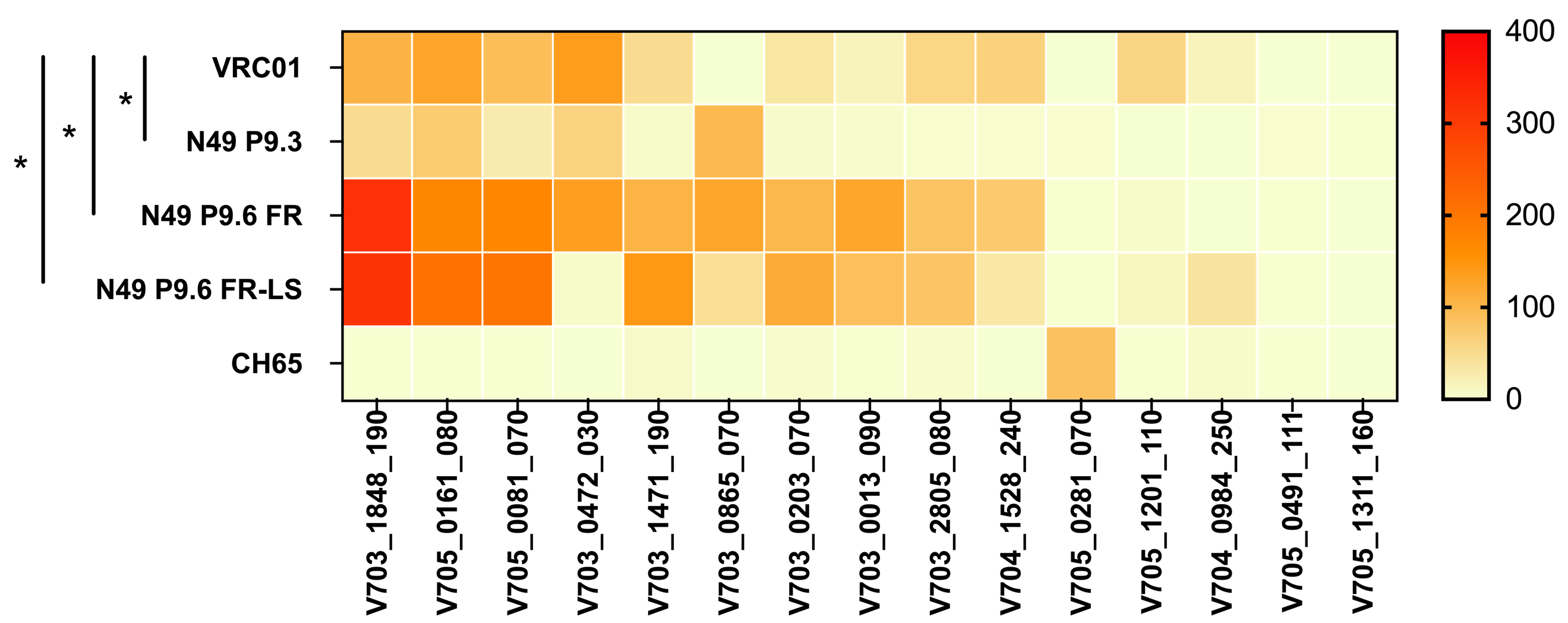

**Figure S2. ADCC activity of N49 P series variants.** The heat map represents the magnitude of responses against infectious molecular clones expressing recent Envelopes from the placebo group of phase II/III trials (HVTN703/704/705) as area under the curve (AUC) from 0 (no activity) to ≥400 (highest) as indicated by the scale on the right y-axis. Antibody-specific killing of each mAb listed on the left y-axis was conducted against each HIV IMC (x-axis) starting at 50µg/mL using a 5-fold dilution. Area-under-the-curves (AUCs) were then calculated using the trapezoid rule after subtracting the background activity. The anti-flu CH65 mAb was used as negative control. The parental antibody mAb N49P9.3 displayed significantly less ADCC activity than VRC01, while the engineered variants N49P9.6-FR, and N49P9.6-FR-LS demonstrated significantly more ADCC activity than VRC01, when tested by paired t test. * = P < .05

= P < .05
