## Supplementary material for "A comprehensive engineering strategy improves potency and manufacturability of a near pan-neutralizing antibody against HIV": Figure S3

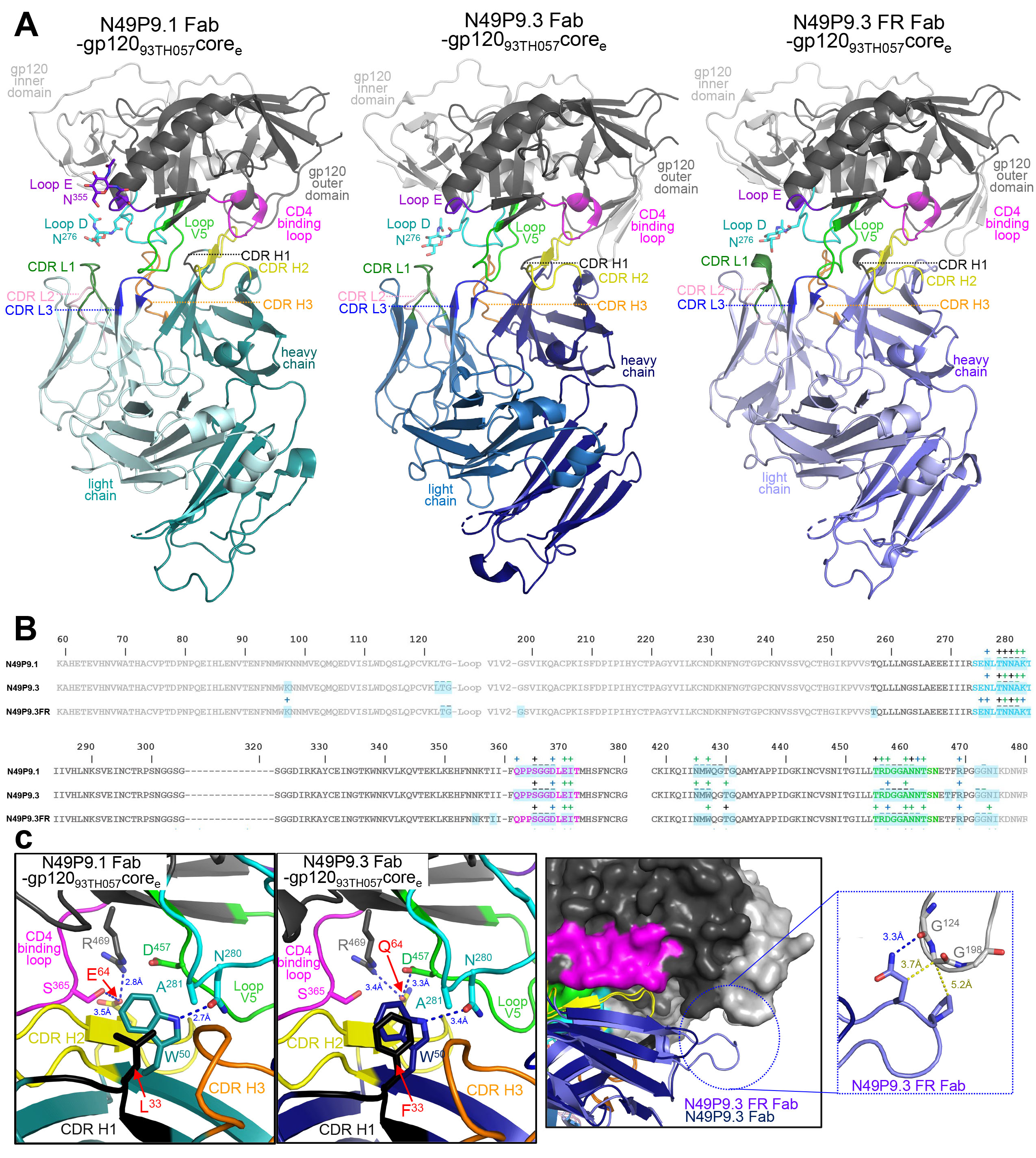

**Figure S3. Crystal structures of N49P9.1Fab-, N49P9.3-Fab- and N49P9.3FR Fab-gp120_93TH057_core_e_ complexes** (**a**) The overall structure of the complexes are shown as ribbon diagrams. The complementary-determining regions (CDRs) of Fabs are colored: CDR L1, green; CDR L2 light pink; CDR L3, blue; CDR H1, black; CDR H2, yellow and CDR H3, light orange. Outer and inner domains of gp120 are dark and light grey, respectively. The outer domain loops: D, E, CD4 binding, and V5 are colored in cyan, purple blue, magenta, and light green, respectively. Carbohydrates at position N^276^ (loop D) and N^355^ (loop E) are shown as sticks. (**b**) Epitope footprints of Fabs N49P9.1 and N49P9.3 mapped onto the gp120 primary sequences. Contact residues are defined by a 5 Å cutoff and marked above the sequence with (+) for side chain and (-) for main chain to indicate the type of contact: hydrophilic (blue), hydrophobic (green) and both (black). Buried surface residues determined by PISA are shaded blue for primary (**c**) Details of N49P9.3 Fab-gp120_93TH057_core_e_ interface with a blow-up view into the Fab contacts mediated by CDR H1 and 2 of N49P9.1 and N49P9.3 with colors indicated as in (a) and hydrogen bonds shown as dotted blue lines *(left panel)*. Interaction network of the frame region of N49P9.3-FR to gp120_93TH057_core_e_ *(right panel)*. H-bonds and Van der Waals contacts are shown as blue and yellow dotted lines, respectively. Residues that differ between N49P9.1 and N49P9.3 are labeled in red. Structures of N49P9.3 with and without the VRC03 FR insertion allowed us to assess the framework’s role in binding monomer gp120. Analysis of the N49P9.1 and N49P9.3 structures reveal that only two gp120 contact residues differ between the two, one in CDRH1 and one in CDRH2 (b and c). The first, Phe33 in CDRH1, packs against gp120 Ala281 in place of Leu33 in N49P9.1, but with slightly better van der Waals contacts, and modulates heavy chain Trp50, which also packs against gp120 Ala281 and forms a hydrogen bond to gp120 Asn280. The second Gln64 in CDRH2 can form hydrogen bonds with both gp120 Arg469 and Asp457 versus Glu64 in N49P9.1 which can only form a hydrogen bond with Arg469. Gln at position 64 is also what is seen in both N49P7 and VRC01 implying that Gln can make an important contribution to binding when placed here [22, 54]. These two amino acid changes likely account for much of N49P9.3’s increase in neutralization potency and breadth relative to N49P9.1. With the addition of the N49P9.3-FR structure We were able to confirm that the insertion in N49P9.3-FR makes little direct contact to gp120 core in the N49P9.3-FR Fab-gp12093TH057coree complex as expected and that the primary gp120 contacts remain intact; the VRC03 framework insertion is only expected to contact the adjacent gp120 protomer in the Env trimer [44]. The only direct contact the insert makes are to two glycines in the truncated V1V2 loop in the gp12093TH057coree construct which are not present in the original V1V2 loop sequence (Figure S3c).
