## Supplementary material for "A comprehensive engineering strategy improves potency and manufacturability of a near pan-neutralizing antibody against HIV": Figure S4

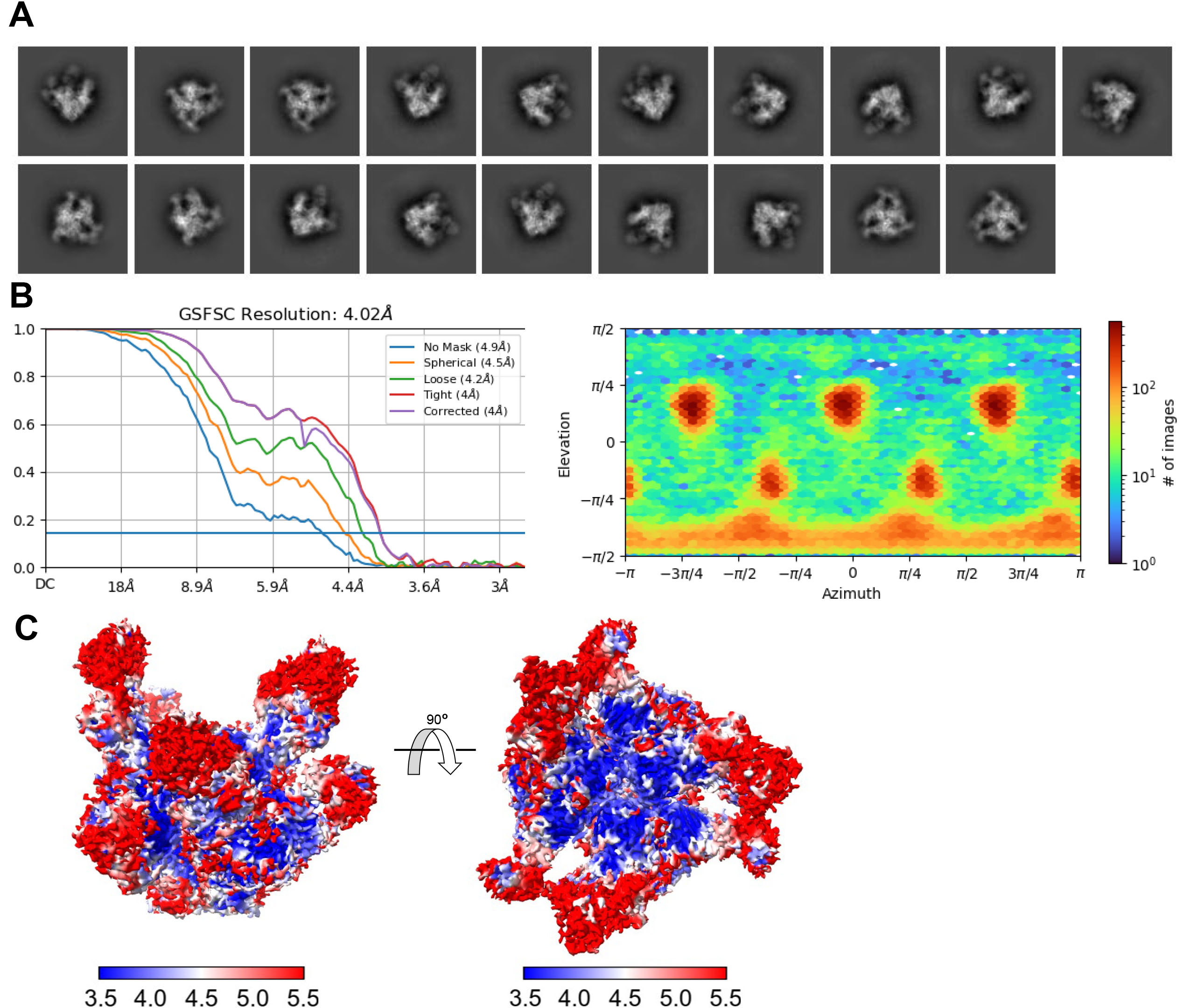

**Figure S4. Cryo-EM structure of BG505 SOSIP.664 HIV-1 bound complex N49P9.6-FR and PGT121** (**a**) Selected 2D classes for *ab initio* map reconstruction(**b**) The Fourier shell correlation curves with spherical mask indicate the overall resolution (FSC cutoff 0.143) as determined by CryoSPARC and the direction distribution plot of all particles used in the final refinement. (**c**) Local resolution estimation plots with top and side views of the complex.
