## Supplementary material for "A comprehensive engineering strategy improves potency and manufacturability of a near pan-neutralizing antibody against HIV": Figure S5

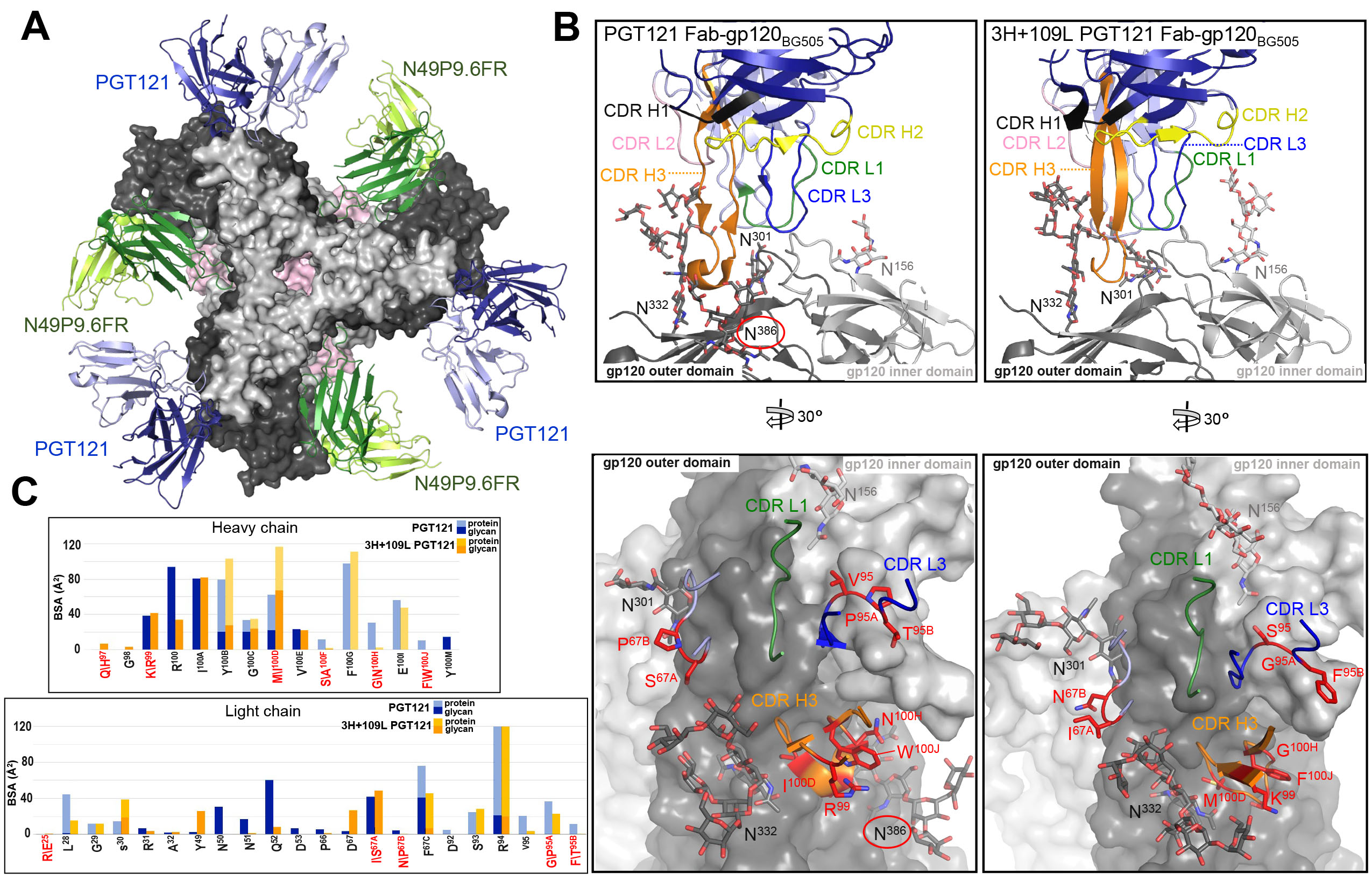

**Figure S5. Molecular details of PGT121 bound to BG505 SOSIP.664.** (**a**) Top view of the complex of BG505 SOSIP.664 HIV-1 Env trimer with N49P9.6-FR and PGT121 bound. N49P9.6-FR Fabs and gp120 are colored as in Figure 3. PGT121 (only the variable regions of the Fabs were built into the Cryo-EM model) are shown in blue and light blue ribbons. (**b**) Side-by-side comparison of PGT121 and the inferred germline PGT121 precursor, 3H+109L PGT121 (PDB ID: 5CEZ). CDRs are colored as in Figure 2. (**c**) BSA plot of Fab residues contributing to binding for PGT121 and 3H+109L PGT121. Protein or glycan portions are as indicated with residues that differ in sequence in red, 3H+109L PGT121 residue bottom and PGT121 residue top. One potential additional glycan interaction between PGT121’s light chain and the glycan attached to N^386^ that is absent in 3H+109L PGT121 is encircled in red in PGT121. The gp120 contact residues are largely identical between PGT121 and 3H+109L PGT121 but interactions PGT121 are more protein dependent. The total BSA for 3H+109L PGT121 is 2275 Å^2^ (1211 Å^2^ for gp120 and 1064 Å^2^ for Fab); total BSA for PGT121 is 2455 Å^2^ (1236 Å^2^ for gp120 and 1219 Å^2^ for Fab). Interactions between protein residues explain the total BSA of 1126 Å^2^ for 3H+109L PGT121 and the total BSA of 1308 Å^2^ for PGT121. Contributions from glycan are roughly comparible for both, 1126 Å^2^ for 3H+109L PGT121 and 1147 Å^2^ for PGT121. Thus one way PGT121 seems to have increased its affinity to Env during maturation is to have maximized its interactions with protein residues while maintaining similar interactions with key glycans.
