## Supplementary material for "A comprehensive engineering strategy improves potency and manufacturability of a near pan-neutralizing antibody against HIV": Figure S6

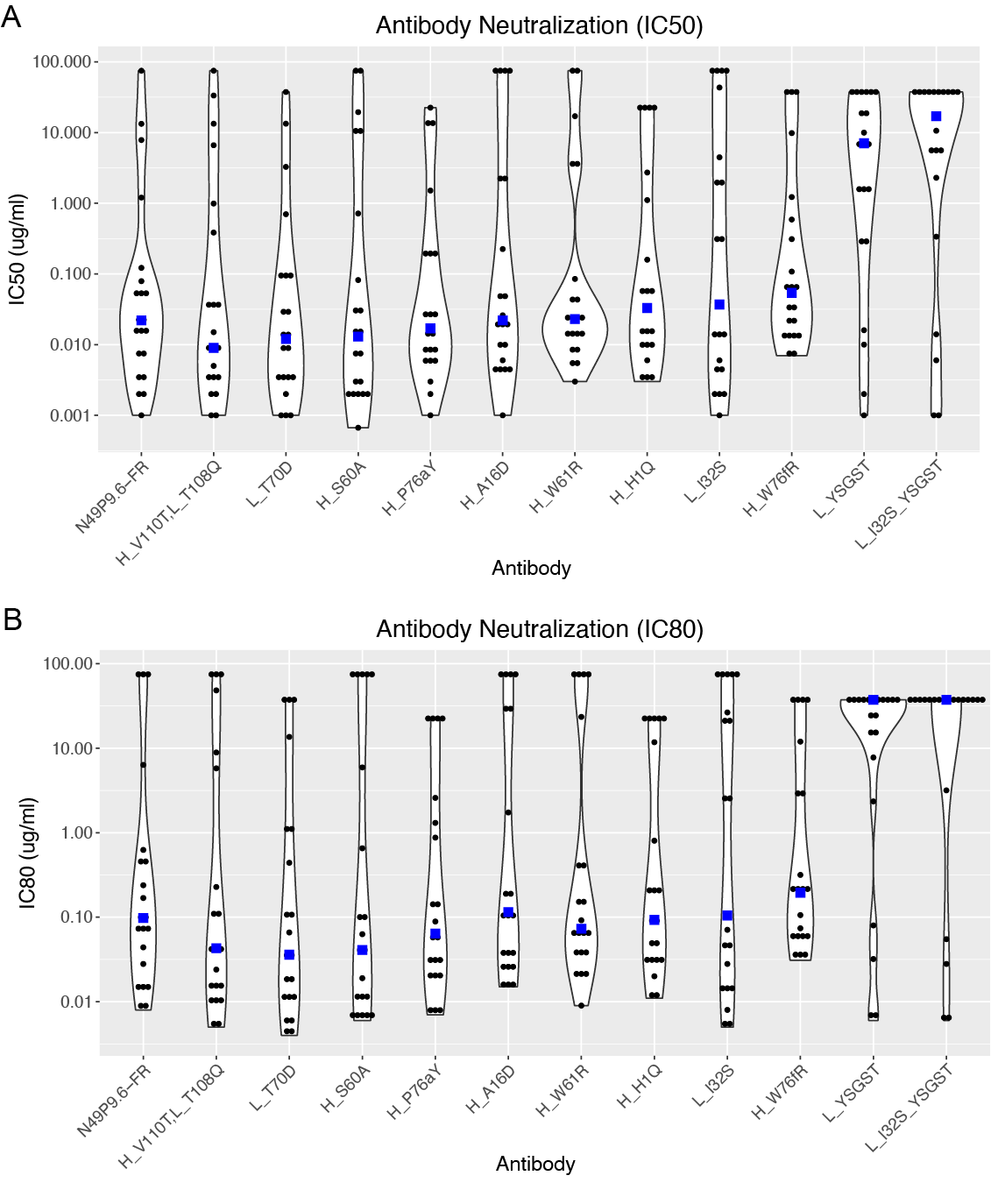

**Figure S6. Neutralization assessment of N49P9.6-FR designs.** Designed antibodies and N49P9.6-FR wild-type were tested for neutralization using a 21 virus global HIV panel, and measured (A) IC50 and (B) IC80 levels are shown as violin plots. Individual neutralization values are shown as black points, with median values shown as blue squares, and designs are ordered from lower to higher median IC50 levels (with wild-type on left for reference). In cases with unquantified neutralization, neutralization values are represented as 1.5-fold versus the unquantified level (e.g. 75 μg/ml for >50 μg/ml). Figure generated with ggplot2 [70].
