## Supplementary material for "A comprehensive engineering strategy improves potency and manufacturability of a near pan-neutralizing antibody against HIV": Figure S7

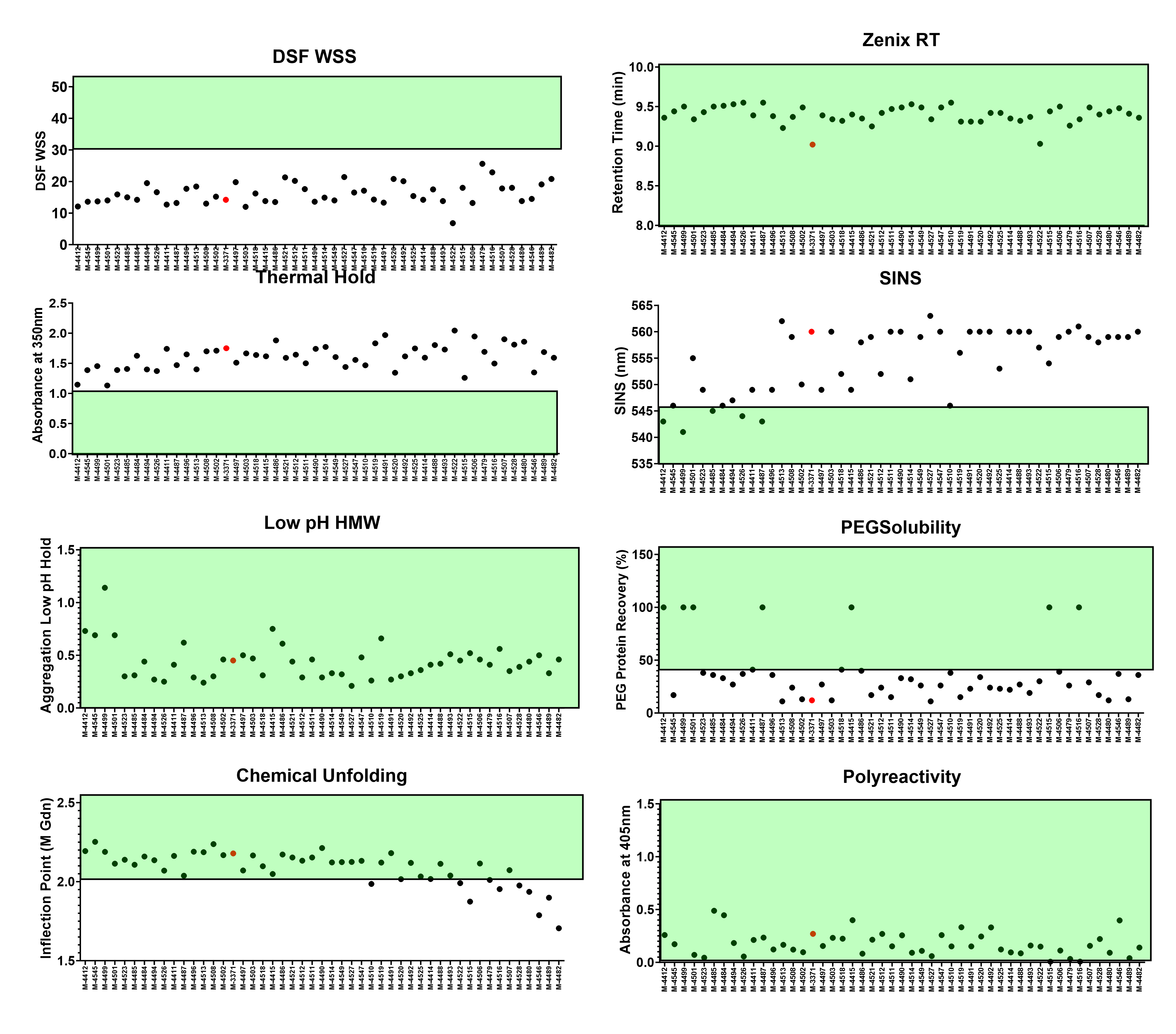

**Figure S7. Conformational and colloidal stability results for variants selected from library panning.** Regions of generally acceptable developability results are highlighted in green, N49P9.6-FR-LS are indicated in red circles compared with variants in black circles. Significant change in conformational stability was not observed while an increase colloidal stability was observed for multiple variants.
