## Supplementary material for "A comprehensive engineering strategy improves potency and manufacturability of a near pan-neutralizing antibody against HIV": Figure S8

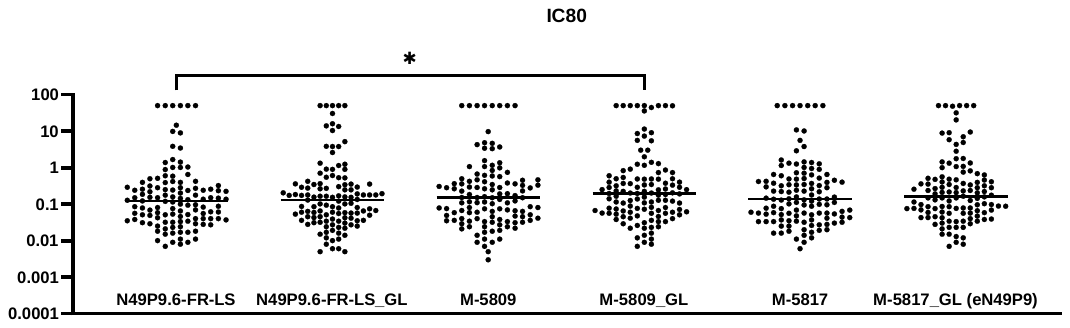

**Figure S8. Neutralizing activity of N49P9.6-FR-LS variants selected for manufacturability characteristics.** Six selected variants (see narrative) were expressed from stable cells pools and tested for neutralization against a 119 multi-tier multi-clade neutralization panel. All the variants tested had less than two fold difference in median IC80 compared to the parental N49P9.6-FR-LS, although this was significant only for M-5809_GL compared to N490P9.6-FR-LS (P=.04 by Mann-Whitney test). Variant M-5817_GL was the final variant chosen for clinical development (eN49P9).

“-GL”: use of light chain with uncleaved N-terminus.
