## Supplementary material for "A comprehensive engineering strategy improves potency and manufacturability of a near pan-neutralizing antibody against HIV": Figure S9

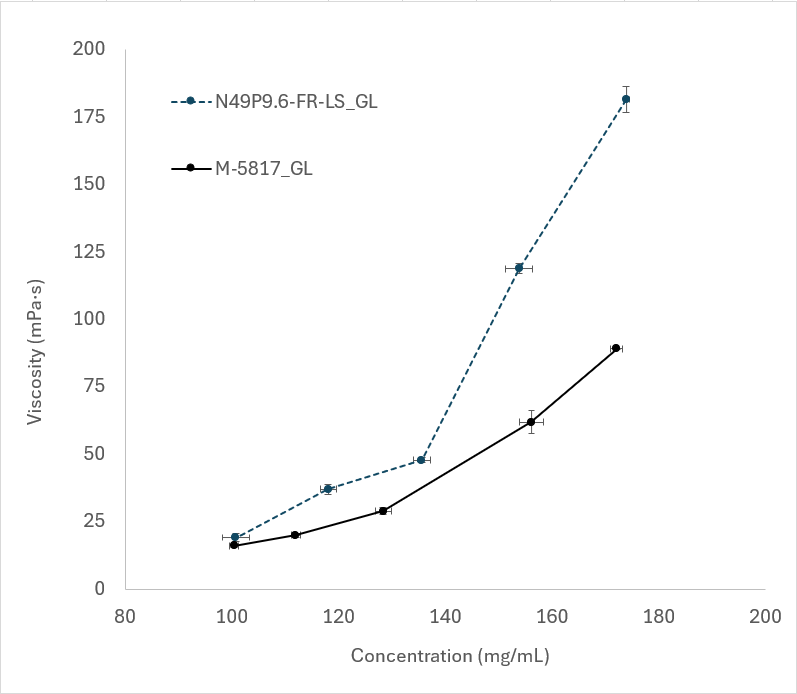

**Figure S9.** **Viscosity measurement of final variant and parental form.** Increased colloidal stability as measured by biophysical analysis was confirmed by viscosity measurement across increasing protein concentration. Decrease in viscosity was observed in the lead candidate, M-5817_GL (black line) as compared with the parental molecule N49P9.6-FR-LS (germline N-term) (blue dotted line). Standard deviation of triplicate protein concentration and viscosity measurements are capture in hash marks for each axis.
