## Supplementary material for "A comprehensive engineering strategy improves potency and manufacturability of a near pan-neutralizing antibody against HIV": Table S1

**Table S1. Single bnAb neutralization characteristics and ranking**

| **bnAb** | **IC80 geometric mean (µg/ml)** | **IC80 < 1µg/ml breadth (%)** | **IC99 geometric mean (µg/ml)** | **IC99 < 10µg/ml breadth (%)** | **IIP at 30µg/ml (Log10)** | **IIP > 5Log10 breadth (%)** | **Average rank** |
| --- | --- | --- | --- | --- | --- | --- | --- |
| N49P9.6-FR | 0.069 | 88.542 | 1.057 | 82.292 | 3.853 | 14.583 | 1.000 |
| VRC01.23LS | 0.198 | 84.375 | 2.578 | 78.125 | 3.274 | 11.458 | 1.857 |
| N6 | 0.326 | 83.333 | 3.613 | 77.083 | 3.267 | 5.208 | 3.286 |
| N49P9.6 | 0.219 | 83.333 | 3.496 | 73.958 | 3.045 | 1.042 | 3.571 |
| 1-18 | 0.264 | 80.208 | 5.506 | 63.542 | 2.744 | 2.083 | 4.286 |
| VRC07-523-LS | 0.333 | 79.167 | 9.072 | 55.208 | 2.484 | 0.000 | 6.286 |
| PGDM1400 | 0.348 | 61.458 | 5.931 | 42.708 | 2.089 | 1.042 | 6.286 |
| VRC26.25 | 0.663 | 52.083 | 7.287 | 35.417 | 1.273 | 6.250 | 6.857 |
| 3BNC117 | 0.941 | 60.417 | 12.287 | 40.625 | 2.231 | 1.042 | 7.143 |
| PGT121 | 1.642 | 40.625 | 10.305 | 37.500 | 0.935 | 1.042 | 8.429 |
| 10-1074 | 2.931 | 42.708 | 14.892 | 31.250 | 0.987 | 2.083 | 8.857 |
| VRC01 | 1.775 | 38.542 | 15.600 | 34.375 | 2.096 | 0.000 | 9.429 |
