## Supplementary material for "A comprehensive engineering strategy improves potency and manufacturability of a near pan-neutralizing antibody against HIV": Table S2

**Table S2. Dual bnAb neutralization characteristics and ranking**

| **bnAb** | **IC80 geometric mean (µg/ml)** | **IC80 < 1µg/ml breadth** | **IC80 breadth with 2 bnAbs active** | **IC99 geometric mean (µg/ml)** | **IC99 < 10µg/ml breadth (%)** | **IIP at 30µg/ml (Log10)** | **IIP > 5Log10 breadth (%)** | **Average rank** |
| --- | --- | --- | --- | --- | --- | --- | --- | --- |
| N49P9.6-FR+PGDM1400 | 0.035 | 93.750 | 64.583 | 0.433 | 89.583 | 5.382 | 55.208 | 1.857 |
| N49P9.6-FR+VRC26.25 | 0.029 | 93.750 | 47.917 | 0.407 | 90.625 | 4.736 | 47.917 | 3.286 |
| VRC01.23LS+PGDM1400 | 0.071 | 89.583 | 64.583 | 0.817 | 87.500 | 4.873 | 47.917 | 4.143 |
| N49P9.6-FR+PGT121 | 0.056 | 91.667 | 47.917 | 0.605 | 90.625 | 4.622 | 44.792 | 4.714 |
| N49P9.6+PGDM1400 | 0.074 | 90.625 | 65.625 | 0.969 | 84.375 | 4.493 | 41.667 | 5.714 |
| VRC01.23LS+VRC26.25 | 0.067 | 88.542 | 50.000 | 0.913 | 84.375 | 4.519 | 40.625 | 6.714 |
| N6+PGDM1400 | 0.106 | 86.458 | 66.667 | 1.187 | 83.333 | 4.686 | 44.792 | 7.714 |
| N49P9.6-FR+10-1074 | 0.075 | 91.667 | 45.833 | 0.787 | 89.583 | 4.295 | 37.500 | 8.000 |
| 1-18+PGDM1400 | 0.086 | 89.583 | 64.583 | 1.173 | 77.083 | 4.242 | 40.625 | 8.857 |
| N49P9.6+VRC26.25 | 0.068 | 88.542 | 48.958 | 1.080 | 83.333 | 3.839 | 36.458 | 10.143 |
| N6+VRC26.25 | 0.100 | 85.417 | 51.042 | 1.282 | 81.250 | 3.990 | 37.500 | 11.857 |
| VRC01.23LS+PGT121 | 0.122 | 87.500 | 46.875 | 1.308 | 83.333 | 4.135 | 38.542 | 11.857 |
| VRC07-523-LS+PGDM1400 | 0.089 | 85.417 | 62.500 | 1.469 | 73.958 | 3.860 | 35.417 | 13.714 |
| 1-18+VRC26.25 | 0.082 | 87.500 | 50.000 | 1.369 | 72.917 | 3.817 | 30.208 | 14.286 |
| N49P9.6+PGT121 | 0.138 | 86.458 | 47.917 | 1.635 | 78.125 | 3.900 | 31.250 | 15.429 |
| N6+PGT121 | 0.201 | 82.292 | 48.958 | 1.912 | 80.208 | 4.067 | 36.458 | 16.286 |
| VRC01.23LS+10-1074 | 0.182 | 85.417 | 44.792 | 1.746 | 82.292 | 3.897 | 32.292 | 17.286 |
| 3BNC117+PGDM1400 | 0.128 | 79.167 | 53.125 | 1.892 | 73.958 | 3.879 | 28.125 | 18.000 |
| VRC07-523-LS+VRC26.25 | 0.085 | 85.417 | 46.875 | 1.906 | 66.667 | 3.182 | 30.208 | 18.571 |
| N49P9.6+10-1074 | 0.198 | 87.500 | 45.833 | 2.099 | 80.208 | 3.695 | 28.125 | 18.857 |
| N6+10-1074 | 0.304 | 81.250 | 46.875 | 2.469 | 78.125 | 4.023 | 35.417 | 19.857 |
| VRC01+PGDM1400 | 0.228 | 71.875 | 56.250 | 2.639 | 71.875 | 3.749 | 30.208 | 21.286 |
| 1-18+PGT121 | 0.159 | 80.208 | 45.833 | 2.325 | 69.792 | 3.223 | 34.375 | 21.429 |
| 1-18+10-1074 | 0.232 | 83.333 | 43.750 | 3.002 | 69.792 | 3.187 | 29.167 | 24.143 |
| 3BNC117+VRC26.25 | 0.115 | 80.208 | 37.500 | 2.320 | 67.708 | 3.154 | 22.917 | 24.286 |
| VRC07-523-LS+PGT121 | 0.164 | 83.333 | 44.792 | 2.928 | 63.542 | 3.124 | 25.000 | 24.286 |
| PGDM1400+PGT121 | 0.110 | 75.000 | 31.250 | 2.208 | 64.583 | 3.345 | 17.708 | 25.286 |
| VRC01+VRC26.25 | 0.233 | 69.792 | 40.625 | 3.088 | 63.542 | 3.311 | 23.958 | 27.857 |
| PGDM1400+10-1074 | 0.165 | 72.917 | 29.167 | 3.049 | 63.542 | 3.053 | 19.792 | 28.714 |
| VRC26.25+PGT121 | 0.164 | 69.792 | 26.042 | 2.256 | 61.458 | 2.889 | 20.833 | 28.857 |
| VRC07-523-LS+10-1074 | 0.251 | 82.292 | 42.708 | 4.197 | 60.417 | 2.762 | 18.750 | 29.857 |
| 3BNC117+PGT121 | 0.296 | 68.750 | 36.458 | 3.859 | 59.375 | 2.987 | 20.833 | 31.286 |
| VRC26.25+10-1074 | 0.232 | 70.833 | 22.917 | 3.376 | 54.167 | 2.555 | 20.833 | 32.429 |
| 3BNC117+10-1074 | 0.456 | 71.875 | 35.417 | 5.151 | 62.500 | 2.827 | 15.625 | 32.571 |
| VRC01+PGT121 | 0.506 | 57.292 | 39.583 | 5.337 | 55.208 | 2.904 | 17.708 | 33.000 |
| VRC01+10-1074 | 0.819 | 56.250 | 36.458 | 7.097 | 55.208 | 2.723 | 17.708 | 34.286 |
