## Supplementary material for "A comprehensive engineering strategy improves potency and manufacturability of a near pan-neutralizing antibody against HIV": Table S3

**Table S3. Triple bnAb neutralization characteristics and ranking**

| **bnAb** | **IC80 geometric mean (µg/ml)** | **IC80 < 1µg/ml breadth** | **IC80 breadth with 2 bnAbs active** | **IC99 geometric mean (µg/ml)** | **IC99 < 10µg/ml breadth (%)** | **IIP at 30µg/ml (Log10)** | **IIP > 5Log10 breadth (%)** | **Average rank** |
| --- | --- | --- | --- | --- | --- | --- | --- | --- |
| N49P9.6 FR+PGDM1400+PGT121 | 0.030 | 94.792 | 82.292 | 0.300 | 93.750 | 6.516 | 71.875 | 2.714 |
| N49P9.6 FR+PGDM1400+10-1074 | 0.037 | 95.833 | 82.292 | 0.367 | 93.750 | 6.303 | 69.792 | 3.286 |
| N49P9.6 FR+VRC26.25+PGT121 | 0.025 | 95.833 | 69.792 | 0.285 | 93.750 | 6.173 | 63.542 | 5.000 |
| VRC01.23LS+PGDM1400+PGT121 | 0.052 | 91.667 | 84.375 | 0.514 | 91.667 | 5.984 | 65.625 | 5.000 |
| N49P9.6 FR+VRC26.25+10-1074 | 0.031 | 96.875 | 70.833 | 0.345 | 94.792 | 6.126 | 62.500 | 5.571 |
| N49P9.6+PGDM1400+PGT121 | 0.054 | 91.667 | 84.375 | 0.585 | 91.667 | 5.712 | 56.250 | 7.143 |
| VRC01.23LS+PGDM1400+10-1074 | 0.068 | 92.708 | 84.375 | 0.638 | 91.667 | 5.692 | 63.542 | 7.143 |
| N49P9.6+PGDM1400+10-1074 | 0.071 | 92.708 | 84.375 | 0.723 | 90.625 | 5.499 | 57.292 | 10.143 |
| N49P9.6+VRC26.25+PGT121 | 0.049 | 92.708 | 71.875 | 0.622 | 89.583 | 5.174 | 57.292 | 10.857 |
| VRC01.23LS+VRC26.25+PGT121 | 0.048 | 89.583 | 71.875 | 0.553 | 85.417 | 5.572 | 61.458 | 10.857 |
| N6+PGDM1400+PGT121 | 0.071 | 88.542 | 83.333 | 0.737 | 86.458 | 5.921 | 60.417 | 12.000 |
| VRC01.23LS+VRC26.25+10-1074 | 0.063 | 91.667 | 72.917 | 0.685 | 86.458 | 5.457 | 55.208 | 12.429 |
| 1-18+PGDM1400+PGT121 | 0.056 | 90.625 | 78.125 | 0.677 | 85.417 | 5.436 | 55.208 | 12.571 |
| N49P9.6+VRC26.25+10-1074 | 0.065 | 93.750 | 72.917 | 0.763 | 91.667 | 5.185 | 53.125 | 12.857 |
| VRC07-523-LS+PGDM1400+PGT121 | 0.056 | 89.583 | 80.208 | 0.776 | 87.500 | 5.337 | 54.167 | 13.429 |
| N6+VRC26.25+PGT121 | 0.066 | 89.583 | 71.875 | 0.759 | 86.458 | 5.478 | 56.250 | 14.286 |
| N6+PGDM1400+10-1074 | 0.096 | 86.458 | 83.333 | 0.923 | 85.417 | 5.636 | 59.375 | 16.571 |
| 1-18+PGDM1400+10-1074 | 0.073 | 90.625 | 78.125 | 0.832 | 84.375 | 5.117 | 51.042 | 17.571 |
| 1-18+VRC26.25+PGT121 | 0.054 | 88.542 | 68.750 | 0.735 | 83.333 | 5.164 | 53.125 | 17.857 |
| N6+VRC26.25+10-1074 | 0.089 | 87.500 | 72.917 | 0.945 | 87.500 | 5.251 | 54.167 | 18.857 |
| 3BNC117+PGDM1400+PGT121 | 0.070 | 88.542 | 73.958 | 0.938 | 84.375 | 4.800 | 46.875 | 19.286 |
| VRC07-523-LS+PGDM1400+10-1074 | 0.075 | 88.542 | 80.208 | 0.993 | 84.375 | 5.068 | 51.042 | 19.286 |
| 1-18+VRC26.25+10-1074 | 0.071 | 89.583 | 69.792 | 0.910 | 83.333 | 4.818 | 46.875 | 20.429 |
| VRC07-523-LS+VRC26.25+PGT121 | 0.053 | 88.542 | 68.750 | 0.878 | 79.167 | 4.775 | 46.875 | 20.429 |
| 3BNC117+PGDM1400+10-1074 | 0.093 | 87.500 | 72.917 | 1.187 | 83.333 | 4.738 | 45.833 | 23.857 |
| 3BNC117+VRC26.25+PGT121 | 0.064 | 88.542 | 63.542 | 1.004 | 79.167 | 4.589 | 41.667 | 24.143 |
| VRC07-523-LS+VRC26.25+10-1074 | 0.070 | 88.542 | 69.792 | 1.144 | 77.083 | 4.559 | 42.708 | 24.143 |
| 3BNC117+VRC26.25+10-1074 | 0.085 | 88.542 | 63.542 | 1.283 | 80.208 | 4.248 | 34.375 | 26.857 |
| VRC01+PGDM1400+PGT121 | 0.103 | 81.250 | 66.667 | 1.378 | 80.208 | 4.767 | 45.833 | 27.000 |
| VRC01+PGDM1400+10-1074 | 0.144 | 81.250 | 66.667 | 1.764 | 77.083 | 4.557 | 42.708 | 29.000 |
| VRC01+VRC26.25+PGT121 | 0.107 | 78.125 | 58.333 | 1.424 | 75.000 | 4.493 | 40.625 | 30.714 |
| VRC01+VRC26.25+10-1074 | 0.149 | 78.125 | 59.375 | 1.825 | 73.958 | 4.144 | 41.667 | 31.286 |
