## Supplementary material for "A comprehensive engineering strategy improves potency and manufacturability of a near pan-neutralizing antibody against HIV": Table S4

**Table S4. N49P9.6-FR-LS neutralization against AMP panel pseudoviruses in TZM.bl cells**

| **Virus ID** | **Clade** | **IC50** | **IC80** |
| --- | --- | --- | --- |
| H704_0128_220_RE_pb001_s | F1/B | **0.004** | **0.015** |
| V704_0128_220_RE_pblib002_s | F1/B | **0.014** | **0.048** |
| H704_0496_040sN | F | **0.112** | **0.517** |
| H704_1109_140_RE_cs | F1/B | **2.364** | **8.459** |
| H704_1180_070EsN | A1/B | **0.003** | **0.013** |
| H704_1180_070_RE_pblib027_s | A1/B | **0.007** | **0.019** |
| H704_1183_220EsN | B/F | **0.011** | **0.038** |
| H704_1429_090_eN1A | F2 | **0.035** | **0.099** |
| H704_1429_090_eN2G | F2 | **0.027** | **0.105** |
| H704_1775_030cN_SynGtoA_567 | F1 | **0.043** | **0.192** |
| H704_1930_170_RE_c01s_1523A | 12_BF,F1 recomb. | **0.011** | **0.030** |
| H704_1930_170_RE_c02s_1523G | 12_BF,F1 recomb. | **0.014** | **0.047** |
| H704_2788_060eN_04 | B/F recomb. | **0.066** | **0.229** |
| H704_2788_060eN_12 | B/F recomb. | **0.076** | **0.196** |
| H703_0013_090Es | C | **0.200** | **0.661** |
| H703_0203_081_RE_e8A5s | C | **0.329** | **1.895** |
| V703_0203_081_RE_pblib002_s | C | **0.362** | **1.283** |
| V703_0217_050_RE_pblib002_s | C | **0.016** | **0.062** |
| H703_0217_050e_2A3 | C | **0.018** | **0.059** |
| H703_0279_110s | C | **0.072** | **0.225** |
| V703_0279_110_RE_pblib002_s | C | **0.060** | **0.169** |
| H703_0472_030s | C | **0.001** | **0.005** |
| H703_0537_011s_4H1 | C | **0.003** | **0.009** |
| H703_0537_011s_2B5 | C | **0.001** | **0.004** |
| V703_0537_110_RE_pblib003_s | C | **0.002** | **0.006** |
| H703_0566_160s | C | **0.051** | **0.165** |
| V703_0566_160_RE_pblib002_s | C | **0.048** | **0.161** |
| V703_0629_150_RE_pblib002_s | C | **0.023** | **0.058** |
| H703_0629_150_RE_e5A2s | C | **0.017** | **0.054** |
| V703_0629_150_RE_pblib003_s | C | **0.020** | **0.047** |
| V703_0629_150_RE_pblib004_s | C | **0.013** | **0.052** |
| H703_0646_051sN | C | **0.470** | **1.561** |
| H703_0739_110s | C | **0.009** | **0.028** |
| H703_0842_200Es | C | **0.975** | **4.210** |
| H703_0926_070s_2H2 | C | **0.346** | **1.643** |
| H703_1104_100_Re10GS | C | **0.537** | **1.675** |
| H703_1104_100_Re10A5s | C | **0.265** | **1.181** |
| V703_1104_100_RE_pblib001_s | C | **0.322** | **1.471** |
| V703_1298_080_RE_pblib002_s | C | **1.985** | **17.891** |
| H703_1313_040s | C | **0.206** | **0.561** |
| H703_1471_190s | C | **0.011** | **0.042** |
| H703_1675_G613s | C | **0.070** | **0.193** |
| H703_1750_140Es | C | **0.251** | **0.700** |
| H703_1764_250_RE_cs | C | **0.028** | **0.082** |
| H703_1789_230_RE_e3A3s | C | **0.072** | **0.206** |
| V703_1848_190_RE_pblib002_s | C | **0.034** | **0.092** |
| H703_1848_190_RE_e6D1s | C | **0.071** | **0.233** |
| V703_1915_250_RE_sgaA3_s | C | **0.056** | **0.190** |
| H703_2018_240_RE_e6A1s | C | **0.051** | **0.233** |
| H703_2117_110_RE_e2A10s | C | **0.035** | **0.083** |
| H703_2149_060_RE_eB10s | C | **0.013** | **0.054** |
| H703_2304_150_RE_cs | C | **0.038** | **0.149** |
| H703_2539_070_RE_e6F6s | C | **0.051** | **0.180** |
| H703_2631_150_RE_e2F8s | C | **0.065** | **0.212** |
| H703_2788_030Es_B1 | C | **0.091** | **0.249** |
| H703_2805_080Es | C | **0.026** | **0.073** |
| V703_2805_080_RE_pblib002_s | C | **6.677** | **24.907** |
