## Supplementary material for "A comprehensive engineering strategy improves potency and manufacturability of a near pan-neutralizing antibody against HIV": Table S5

**Table S5. Crystallographic Data collection and refinement statistics**

|  | N49P9.1 Fab-gp120_93TH057_  core_e_ | N49P9.3 Fab-gp120_93TH057_  core_e_ | N49P9.3-FR Fab-gp120_93TH057_  core_e_ |
| --- | --- | --- | --- |
| **Data collection** |  |  |  |
| Wavelength, Ǻ | 0.979 | 0.979 | 0.979 |
| Space group | I2_1_2_1_2_1_ | P1 | P1 |
| Cell parameters |  |  |  |
| a, b, c, Å | 104.8, 110.5, 152.2 | 62.2, 68.2, 115.1 | 60.4, 65.4, 112.4 |
| α, β, γ, ° | 90, 90, 90 | 90.3, 102.3, 90.3 | 90.0, 104.9, 90.0 |
| Complexes/a.u. | 1 | 2 | 2 |
| Resolution, (Å) | 50-3.15 (3.2-3.15) | 50-3.15 (3.46-3.4) | 50-2.15 (2.19-2.15) |
| # of reflections |  |  |  |
| Total | 59,116 | 85,037 | 132,781 |
| Unique | 14,779 (746) | 25,011 (1,163) | 73,767 (2,786) |
| R_merge_^a^, % | 13.8 (79.4) | 11.7 (100) | 13.5 (64.5) |
| R_pim_^b^, % | 7.2 (41.4) | 7.4 (72.5) | 13.5 (64.5) |
| *CC_1/2_*^c^ | 0.99 (0.65) | 0.99 (0.82) | 0.95 (0.75) |
| I/σ | 12.8 (1.0) | 15.5 (1.4) | 12.8 (1.0) |
| Completeness, % | 94.4 (96.4) | 97.6 (92.6) | 82.7 (63.6) |
| Redundancy | 4.0 (4.0) | 3.4 (2.9) | 1.8 (1.4) |
| **Refinement Statistics** |  |  |  |
| Resolution, Å | 50.0 – 3.15 | 50.0 – 3.4 | 50.0 – 2.15 |
| R^d^ % | 24.3 | 23.4 | 20.9 |
| R_free_^e^, % | 29.5 | 28.7 | 26.6 |
| # of atoms |  |  |  |
| Protein | 5,662 | 11,710 | 11,917 |
| Water | – | – | 443 |
| Ligand/Ion | 155 | 285 | 367 |
| Overall B value (Å)^2^ |  |  |  |
| Protein | 125 | 156 | 49 |
| Water | – | – | 43 |
| Ligand/Ion | 135 | 169 | 65 |
| RMSD^f^ |  |  |  |
| Bond lengths, Å | 0.006 | 0.004 | 0.006 |
| Bond angles, ° | 1.0 | 0.9 | 0.95 |
| Ramachandran^g^ |  |  |  |
| favored, % | 85.0 | 89.2 | 93.6 |
| allowed, % | 11.2 | 7.9 | 5.0 |
| outliers, % | 3.8 | 2.9 | 1.4 |
| PDB ID | 6OZ3 | 7SX6 | 7SX7 |

Values in parentheses are for highest-resolution shell

^a^*R*_merge_ = ∑│*I* - <*I*>│/∑*I*, where *I* is the observed intensity and <*I*> is the average intensity obtained from multiple observations of symmetry-related reflections after rejections

^b^R_pim_ = as defined in [98]

^c^*CC_1/2_* = as defined by Karplus and Diederichs [99]

^d^*R* = ∑║F_o_│- │ F_c_║/∑│F_o_ │, where F_o_ and F_c_ are the observed and calculated structure factors, respectively

^e^R_free_ = as defined by Brünger [100]

^f^RMSD = Root mean square deviation

^g^Calculated with MolProbity
