## Supplementary material for "A comprehensive engineering strategy improves potency and manufacturability of a near pan-neutralizing antibody against HIV": Table S6

**Table S6. CryoEM Data collection and refinement statistics**

| **BG505 SOSIP.664-PGT121-N49P9.6-FR complex** | | |
| --- | --- | --- |
|  | Holey grid | Holey grid with 2 nm carbon film |
| **Data Collection** |  |  |
| Microscope | FEI Glacios | FEI Glacios |
| Voltage (kV) | 200 | 200 |
| Total exposure dose (e^-^/Å^2^) | 54.9633 | 58.8084 |
| Detector | Gatan K3 | Gatan K3 |
| Pixel Size (Å) | 0.8893 | 0.8893 |
| Defocus Range (µm) | 0.5-2.7 | 0.5-2.7 |
| Magnification | 45,000 | 45,000 |
| **Reconstruction** |  |  |
| Software | CryoSPARC | CryoSPARC |
| Micrographs collected | 1,900 | 2,468 |
| Number particles extracted/final | 1,291,812 / 48,631 | 1,859,611 / 83,424 |
| Symmetry | C3 | C3 |
| Box size (pix) | 256 | 256 |
| Resolution (Å) (FSC 0.143) | 4.02 | |
| **Refinement (Phenix) & validation** |  | |
| Protein residues | 3,144 | |
| Chimera CC | 0.71 | |
| EMRinger Score |  | |
| Bond lengths (Å) | 0.006 | |
| Bond angles (˚) | 1.04 | |
| Molprobity score | 2.17 | |
| Clash score | 13.5 | |
| Rotamer outliers (%) | 0.11 | |
| Ramachandran |  | |
| Favored (%) | 90.7 | |
| Disallowed (%) | 0.3 | |
| EMDB | EMD-26648 | |
| PDB | 7UOJ | |
