## Supplementary material for "A comprehensive engineering strategy improves potency and manufacturability of a near pan-neutralizing antibody against HIV": Table S7

**Table S7. Changes to the BG505 SOSIP.664 trimer upon binding to CD4bs mAbs that contact the adjacent gp120 protomer.**

| Trimer/mAb | a (Å) | b (Å) | c (Å) | d (Å) | e (Å) | f (Å) | Rotation | Rotation (average) |
| --- | --- | --- | --- | --- | --- | --- | --- | --- |
| BG505 SOSIP.664/ N49P9.6-FR | 46.76 | 46.65 | 46.72 | 56.67 | 56.80 | 56.59 | 1.65, 1.48, 1.56 | 1.5 |
| BG505 SOSIP.664 (4ZMJ) | 45.31 | 45.33 | 45.38 | 54.48 | 54.48 | 54.48 | 0, 0, 0 | 0 |
| BG505 SOSIP.664/ VRCO3 (6CDI) | 46.70 | 46.65 | 46.63 | 55.63 | 55.64 | 55.50 | 2.81, 2.87, 2.89 | 2.9 |
| BG505 SOSIP.664/ N49P6 (6OZ4)) | 46.03 | 46.04 | 46.10 | 54.45 | 54.46 | 54.46 | 1.07, 1.07, 1.07 | 1.1 |
| BG505 SOSIP.664/ VRCO1-FR (6NNF) | 45.90 | 45.91 | 45.97 | 55.25 | 5.25 | 55.25 | 0.724, 0.722, 0.721 | 0.72 |
| BG505 SOSIP.664/ N6-FR (6NM6) | 45.61 | 45.82 | 45.70 | 55.35 | 55.35 | 55.35 | 1.53, 0.848, 0.667 | 0.76 |
| BG505 SOSIP.664/ CD4 (5THR) | 49.76 | 50.84 | 51.16 | 65.60 | 65.49 | 65.55 | 63.8, 65.1, 63.7 | 64.4 |

The degree of ‘trimer opening’ is calculated as described in [72] and defined as the change in position of gp120 relative to gp41 of BG505 SOSIP.664 bound to antibody as compared to unliganded, apo BG505 SOSIP.664 (PDB ID: 4ZMJ[101]). The relative position for each gp120 in the trimer is calculated based on the α-carbon position for residue 375 at the base of the CD4 Phe43 binding pocket relative to the gp41 trimer center (calculated for all trimers aligned based on the α-carbon positions of the central gp41 α7 helices). The distances between Centr and the ^375^C_α_ of each protomer (a, b c) and the ^375^C_α_ atoms of neighboring protomers (d, e, f) are shown to indicate the extent of the protomer rearrangement relative to gp41. The clockwise rotations of the gp120 subunits are calculated as angles relative to apo BG505 SOSIP.664. The BG505 SOSIP.664 bound to CD4 (PDB: 5THR[102]) is shown as a reference to an ‘open’ CD4-triggered conformation of trimer.
