## Supplementary material for "A comprehensive engineering strategy improves potency and manufacturability of a near pan-neutralizing antibody against HIV": Table S8

| **Table S8. Viral neutralization and expression yield for N49P9.6-FR3 antibody designs.** | | | | |
| --- | --- | --- | --- | --- |
| **Design^1^** | **Approach^2^** | **Median IC50 (μg/ml)^3^** | **IC50 fold change^4^** | **Yield (μg/ml)^5^** |
| N49P9.6-FR (wild type) | - | 0.022 | - | 13.2 |
| H_S60A | Glycan site mutation | **0.013** | **1.7** | 13.0 |
| H_H1Q | Rational design | 0.033 | 0.7 | 13.0 |
| H_A16D | Rational design | 0.022 | 1.0 | **18.5** |
| H_W61R | Rational design | 0.023 | 1.0 | **16.1** |
| L_I32S | Rational design | 0.037 | 0.6 | **17.3** |
| L_T70D | Rational design | **0.012** | **1.8** | 8.0 |
| L_FDDK49YSGST | CDRL2 swap | 7.1 | 0.003 | 6.2 |
| L_I32S_FDDK49YSGST | Rational design, CDRL2 swap | 17 | 0.001 | 8.6 |
| H_P76aY | FR3 loop design | **0.017** | **1.3** | 7.0 |
| H_W76fR | FR3 loop design | 0.054 | 0.4 | **52.2** |
| H_V110T,L_T108Q | V-C hinge consensus | **0.009** | **2.4** | **18.3** |

Improved measured properties for designs versus wild-type are shown in bold.

^1^Designs are annotated by chain (H: heavy, L: light) and residue substitution. Substitution FDDK49YSGST corresponds to a CDRL2 residue swap with the VRC07 antibody.

^2^Design approach utilized. Rational design corresponds to germline or consensus residue changes assessed using the Therapeutic Antibody Profiler [36] and Rosetta [37] tools. FR3 design corresponds to structure-based mutagenesis of FR3 loop residues using Rosetta. The V-C consensus design corresponds to simultaneous substitution of variable-constant domain junction residues to consensus residues from related antibodies.

^3^Median IC50 calculated based on neutralization with a global panel of 21 HIV viruses.

^4^Expression yield for 50 ml transfectant, with yield factor corresponding to fold change of yield versus wild-type.
