## Supplementary material for "A comprehensive engineering strategy improves potency and manufacturability of a near pan-neutralizing antibody against HIV": Table S9

**Table S9. Final variant set showing biophsyical characterization analysis**

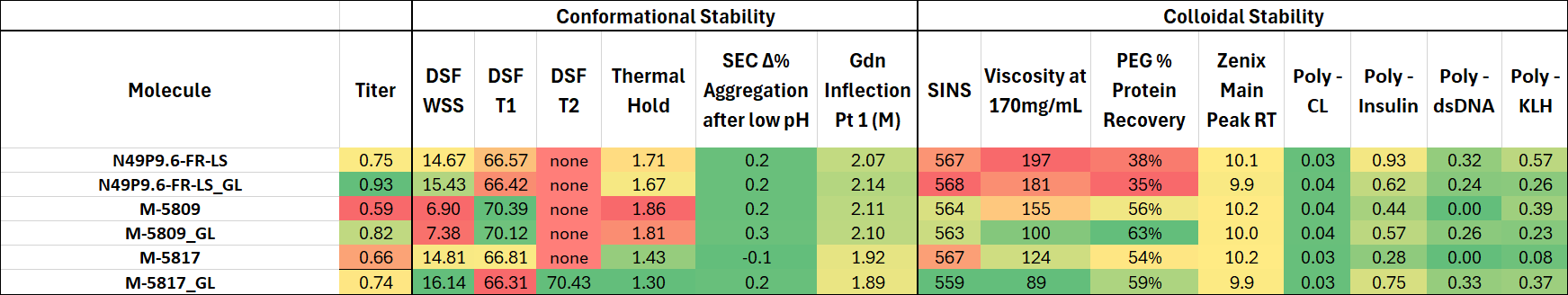

Conditional formating was applied relative to the lowest and highest values within the set for columns 1-5 and 8-10. Columns 6, 7, and 11-15 were conditionally formated to acceptable ranges to illustrate that all molecules in the set were not significantly different. Green colors indiate favorable results, red unfavoratble and yellow moderate.
